# Genetic architecture of collective behaviors in zebrafish

**DOI:** 10.1101/350314

**Authors:** Wenlong Tang, Guoqiang Zhang, Fabrizio Serluca, Jingyao Li, Xiaorui Xiong, Matthew Coble, Tingwei Tsai, Zhuyun Li, Gregory Molind, Peixin Zhu, Mark C. Fishman

## Abstract

Collective behaviors of groups of animals, such as schooling and shoaling of fish, are central to species survival, but genes that regulate these activities are not known. Here we parsed collective behavior of groups of adult zebrafish using computer vision and unsupervised machine learning into a set of highly reproducible, unitary, several hundred millisecond states and transitions, which together can account for the entirety of relative positions and postures of groups of fish. Using CRISPR-Cas9 we then targeted for knockout 35 genes associated with autism and schizophrenia. We found mutations in three genes had distinctive effects on the amount of time spent in the specific states or transitions between states. Mutation in *immp2l* (inner mitochondrial membrane peptidase 2-like gene) enhances states of cohesion, so increases shoaling; mutation in in the Nav1.1 sodium channel, *scn1lab*+/− causes the fish to remain scattered without evident social interaction; and mutation in the adrenergic receptor, *adra1aa*−/−, keeps fish close together and retards transitions between states, leaving fish motionless for long periods. Motor and visual functions seemed relatively well-preserved. This work shows that the behaviors of fish engaged in collective activities are built from a set of stereotypical states. Single gene mutations can alter propensities to collective actions by changing the proportion of time spent in these states or the tendency to transition between states. This provides an approach to begin dissection of the molecular pathways used to generate and guide collective actions of groups of animals.

## Introduction

Darwin recognized that survival depends not only upon the attributes of individuals but also upon coordination of larger groups, such as schools of fish, flocks of birds, and armies of ants (Darwin, 1871). The benefits of collective movement include improved predator avoidance (Ioannou, Guttal, & Couzin, 2012; Ioannou, Morrell, Ruxton, & Krause, 2009), foraging success (Krause, Hartmann, & Pritchard, 1999; Torney, Berdahl, & Couzin, 2011), and reduction in energy consumption due to favorable local currents for fish and birds (Krause, 1994; Spedding, 2011). It is not known whether and how genetic changes in individuals affect group dynamics.

The schooling, shoaling, and other collective dynamics of fish have been beautifully described (Jolles, Boogert, Sridhar, Couzin, & Manica; Katz, Tunstrom, Ioannou, Huepe, & Couzin, 2011; Tunstrom et al., 2013). It appears that the sometimes rapid changes in direction and mode result from propagation of local interactions (Katz et al., 2011). The dynamics can be mathematically modelled as self-propelled particles subjected to forces such as attraction, repulsion, and alignment (Katz et al., 2011; Lukeman, Li, & Edelstein-Keshet, 2010; Sumpter, 2010). No genes regulating such forces have been identified, although QTL linkage analysis suggests that schooling at least is under genetic control (Greenwood, Wark, Yoshida, & Peichel, 2013).

Machine learning has begun to make more tractable the studies of complex behaviors (Anderson & Perona, 2014), such as *C. elegans* foraging (Greene, Brown, et al., 2016), hydra basal behaviors (Han, Taralova, Dupre, & Yuste, 2018), *Drosophila* mating and aggression (Dankert, Wang, Hoopfer, Anderson, & Perona, 2009; Kabra, Robie, Rivera-Alba, Branson, & Branson, 2013; Kravitz & Huber, 2003), and mouse social behaviors (Hong et al., 2015; Kabra et al., 2013; Wiltschko et al., 2015). Unsupervised machine learning approaches, in particular, have the power to capture unknown patterns with as few *a priori* assumptions as possible (Todd, Kain, & de Bivort, 2017), and are widely used in pattern learning, (Feng et al., 2017; Todd et al., 2017) even in cases where data is dense and not readily separable (Chuang, Tzeng, Chen, Wu, & Chen, 2006; Siew, 2013).

Here we have found that unsupervised machine learning can describe fairly completely the collective motion of groups of adult zebrafish as a set of 17 stereotypical states and transitions between states. (Although most studies of zebrafish behavior use larvae (Baier, 2000; Brockerhoff et al., 1995; Friedrich, Jacobson, & Zhu, 2010; Granato et al., 1996; Muto et al., 2005; Portugues & Engert, 2009; Rihel et al., 2010; Zhu et al., 2009), we had to use adult fish because we found robust social behaviors developed late.) We mutated by CRISPR-Cas9 35 genes linked to human disorders of social behavior, such as autism and schizophrenia, with the hope that these genes might play a conserved social behavioral role through evolution. Overall these mutations did not completely add or delete states. Three mutations in particular, however, changed overall usage of states or transitions, with dramatic and consistent effects upon the collective group dynamics.

## Results

We first defined patterns of wild-type behavior. We find that the collective activity became more robust and the behavioral repertoire expanded as the animals grew (Supplemental Figure 1), so it was essential for our studies to use adult fish, at least 3 months of age. We tracked groups of adult fish by continuous video monitoring over half hour assays in an environmentally sheltered tank. As shown in the video attached to Figure 1 (Supplemental Video 1), wild-type fish tend to swim continuously and dynamically in relatively tight groups.

**Figure 1.**
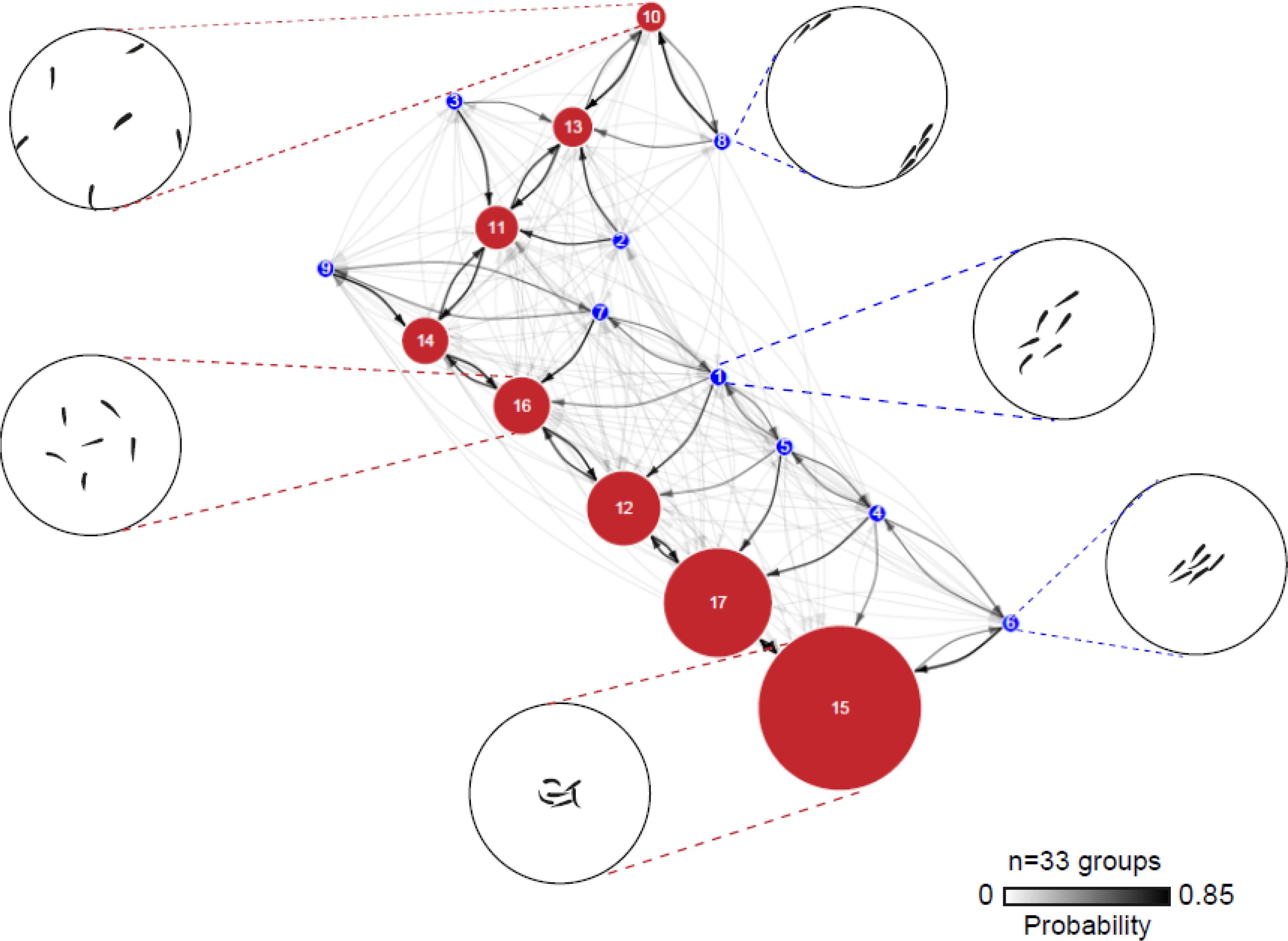
Ethogram of collective behavior in wild-type fish. Circle sizes correspond to state usage ratios, and density of arrows to transition probabilities, with polarized states in blue and unpolarized states in red (n= 33 groups). Dendrogram of state usage is shown in Supplemental Figure 2. Schematic representations (not to scale) of selected states from both categories are shown as blow-ups.

### Description of collective activity

Ethograms are convenient ways to display the amount of time spent in each state as well as the probability of transition between states (Anderson & Perona, 2014; Dankert et al., 2009). The ethogram of group behavior of wild-type fish is shown in Figure 1. Each circle represents a different state (with selected examples shown in blow-ups). For clarity, we have subdivided polarized (school) states (blue) from unpolarized (shoal) (red). The thickness of arrows shows transition probabilities between states. Certain states and transitions are used far more frequently than others (Supplemental Figures 2 and 3). The mean time spent in each state as a block is 0.78 seconds (Supplemental Figure 4). In this assay, fish spend far more time shoaling than schooling, and generally in more cohesive shoals, as shown by the prominence of states in the lower portion of the ethogram.

#### Mutations

For mutational analysis, we elected to mutate genes putatively related to autism or schizophrenia (Supplemental Table 1 and 2) because of the profound defects in social behavior that characterize these disorders, with the hope that their roles in social behaviors might be evolutionarily conserved, even if the behaviors themselves are species-specific. In order to provide sufficiently accurate information for design of effective CRISPR-Cas9 guide RNAs, we found it necessary to sequence and reassemble the complete genome of the AB strain (data available upon request).

We generated mutations in 35 genes (Supplemental file 1) and screened for changes to collective behavior in adult fish homozygous for the mutation (with the exception of *scn1lab* and *slc18a2* which were not viable as homozygous animals so were evaluated as heterozygous animals). Behavior for each mutation was quantified on ethograms generated as above, and then all 35 compared to wild type AB. Using the Kullback-Leibler divergence approach (Kullback & Leibler, 1951) we identified three mutations with particularly distinctive behavioral changes. Differences from wild-type were cross-validated by multi-class Support Vector Machines (SVM) (Supplemental Figure 5), and finally assessed by human expert validation using double blinded datasets. The ethograms for these three mutant fish are shown in Figure 2 and 3, along with exemplary videos. Some other mutations changed state usage in manner similar to the three described, but to a lesser degree (Supplemental Figure 6).

##### Enhanced cohesion

Fish mutant in the inner mitochondrial membrane peptidase 2-like gene, *immp2l*−/−, tend to aggregate tightly in small shoals, as shown in the video attached to Figure 2b (Supplemental Video 2). The increased shoaling is reflected in enhancement of two tight cohesion states in the lower part of the ethogram of Figure 2b (n=9 groups, p<0.001; Mann-Whitney U-test; Supplemental Figure 6a). This is the only mutation to completely abolish individual states (states 8 and 10 are missing). In the contrast with *scn1lab*+/−the two missing states are the most dispersed polarized and unpolarized states._The fish can move dynamically between the limited states, and rarely polarize (usage ratios shown in Supplemental Figure 3). Cross validation of this robust phenotype was performed by multi-class SVM (Prediction Accuracy (PA) =0.95; Leave-One-Out; Supplemental Figure 5). These fish feed well and spawn (Figure 4), and in the home tank movements are indistinguishable from normal.

**Figure 2.**
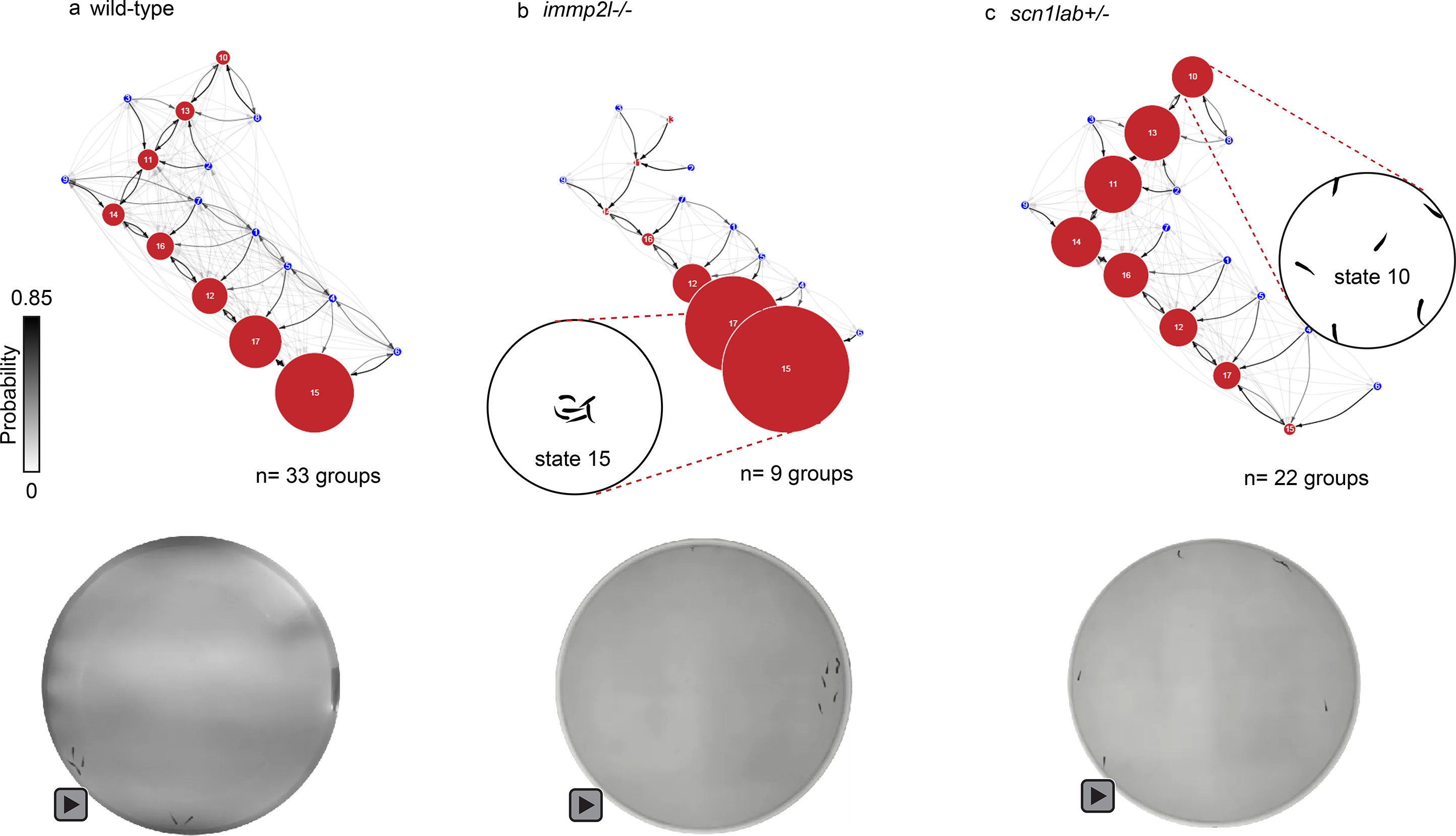
Ethograms of (a) wild-type fish; (b) *immp2l*−/− mutant fish, showing enhanced cohesion (note that dispersed polarized state 8 and unpolarized state10 are missing); (c) *scn1lab*+/− mutant fish, showing enhanced dispersion. Videos can be accessed by clicking where shown, or via Supplemental Video 1 (wild-type), Supplemental Video 2 (*immp2l*−/−), and Supplemental Video 3 (*scn1lab +/−*). Usage ratios are quantified in Supplemental Figure 3 and 4. (wild-type, n=33 groups; *immp2l*−/−, n=9 groups; *scn1lab*+/−, n=22 groups). Color bar indicates transition probability: [0 0.85].

**Figure 3.**
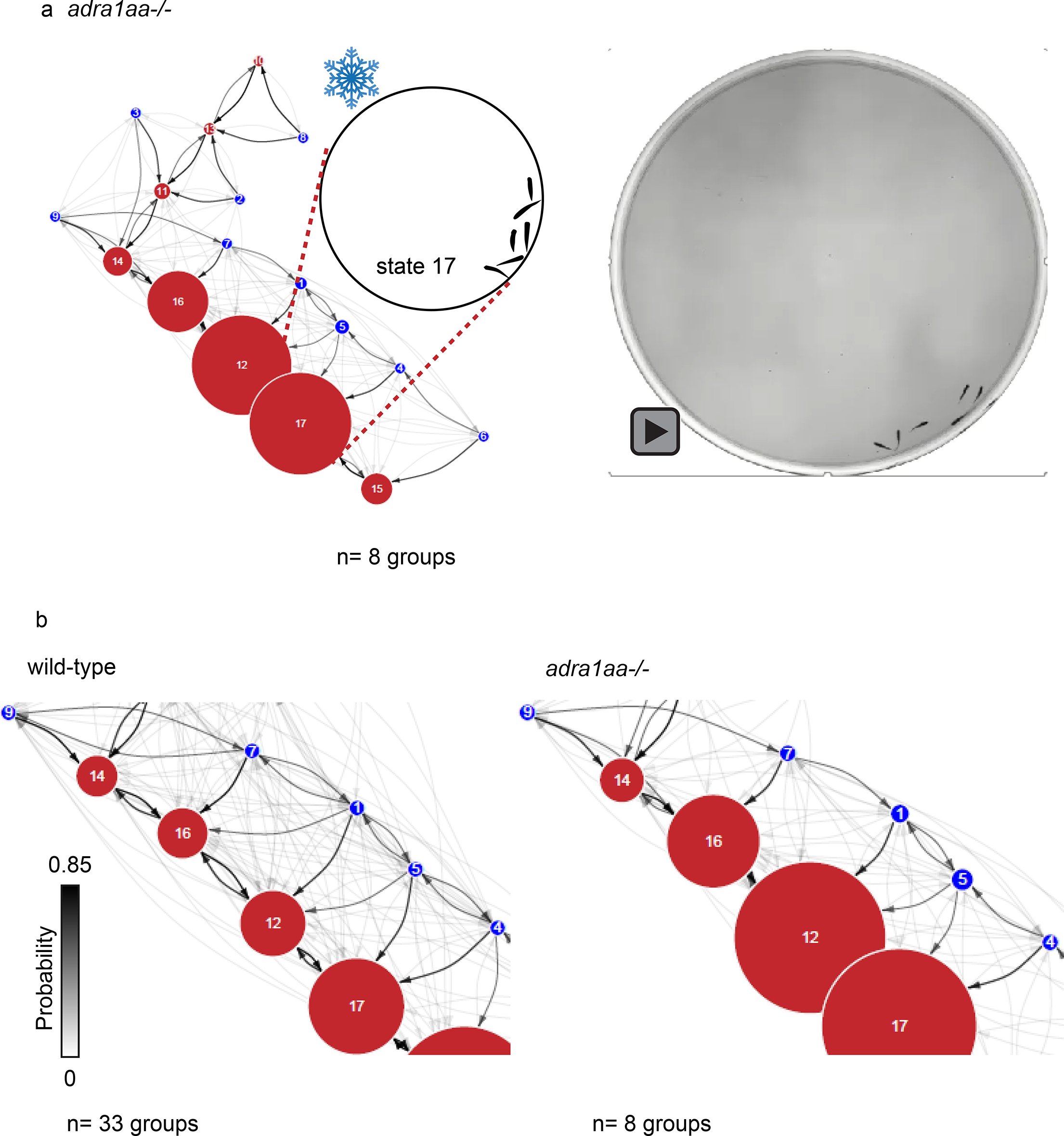
(a) Ethogram of *adralaa*−/− mutant fish (n = 8 groups) showing reduced transitions. Video (Supplemental Video 4) is clickable, and shows bouts of “freezing”. (b) blow-ups of the lower part of the ethograms, comparing wild-type to *adra1aa*−/− fish, to illustrate the relative paucity of transitions between states of *adra1aa*−/− fish. Color bar indicates transition probability: [0 0.85].

##### Enhanced dispersion

Fish mutant in the Nav1.1 sodium channel, *scn1lab*+/−, remain dispersed throughout the chamber, rarely aggregating, as shown in Figure 2c and its attached video (Supplemental Video 3). This is represented by the shift in predominant states of the ethogram to states to the upper right compared to wild-type (n=22 groups; p<0.001; Mann-Whitney U-test; Supplemental Figure 6b). In fact, the states lost in *immp2l*−/− (states 8 and 10) are enhanced in *scn1lab*+/−. Cross validation of this robust phenotype was performed by multi-class SVM (PA=0.88; Leave-One-Out; Supplemental Figure 5). The group centroid speed, pairwise distance, and polarization have little correlation with each other (Supplemental Figure 7). The fish feed, spawn and grow normally.

##### Enhanced freezing

Fish mutant in the adrenergic receptor, *adra1aa*−/−, have many more periods of inactivity than do wild-type, especially towards the end of the half hour test. The state usage of the selected top two states for each comparison is not very distinct from wild type (n=8 groups; Tight shoaling: p=0.59; Medium schooling: p=0.86 and Dispersion: p=0.07; Supplemental Figure 6a-b) as shown in Figure 3a. Compared to wild-type, *adra1aa*−/− mutant fish tend to have significantly more “freeze” motifs in the last third of the recording time (p<0.001), as shown in Supplemental Figure 6c. This difference in dynamics is shown by the relative paucity of arrows in the ethogram blow up of Figure 3b and the attached video (Supplemental video 4). Cross validation of this robust phenotype was performed by multi-class SVM (PA=0.92; Leave-One-Out; Supplemental Figure 5). The fish feed and spawn normally, and their behavior in the home tank is indistinguishable from wild-type.

##### Effects on other behaviors

We were curious as to whether the mutations that perturb collective activity also markedly affect motor control or vision. In terms of motor function, we measured speed, the frequency of acceleration, and turns of individual fish in mating behavior. The three mutations with collective behavioral phenotypes noted above can reach the same level or even higher in all three motion measurements compared to wild type fish. We explored mating, another social behavior, a dance with strong and close proximate interactions, on the assumption that subtle visual or motor defects would perturb the effectiveness of this rapid and highly stereotyped interaction (Nasiadka & Clark, 2012; Spence, Gerlach, Lawrence, & Smith, 2008). It also has a quantitative output, i.e., number of fertilized eggs. The spawning consequences of the mutations were modest, *immp2l*−/− and *adra1aa*−/− slightly increasing and *scn1lab*+/− slightly reducing spawning (Figure 4). Hence the genes that dictate collective behavior have only modest effects on the individual behaviors or mating capabilities that we measured.

**Figure 4.**
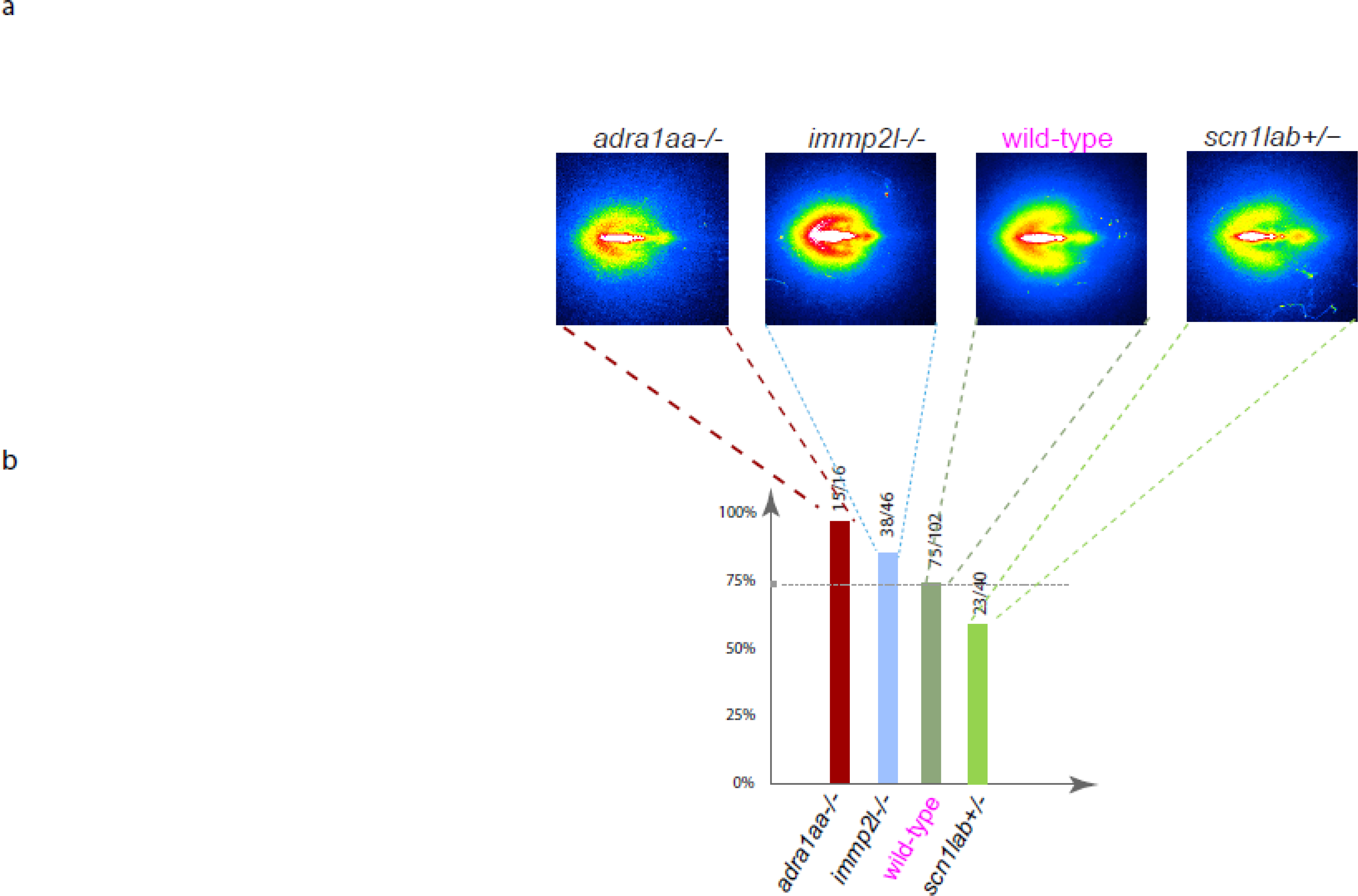
Courtship and spawning is relatively preserved in mutant fish. (a) Positional maps of chasing behavior, with female fish centered facing right and male fish chasing position frequency plotted as a heat map, shows chasing behavior is relatively retained in the mutations of interest. (b) Success of courtship behaviors from mutants and wild-type ranked by the ratio of pairs yielding fertilized embryos, showing modest effects on spawning of the mutations of interest.

## Discussion

Darwin (Darwin, 1871) and Tinbergen (Tinbergen, 1952) speculated as to why some species live in groups and others as solitary animals. Presumably, the benefits of living and moving in a collective provide some advantages (Couzin, 2009; Sumpter, 2010). It is believed that fish school and birds flock in part to help avoid or confuse predators, to improve foraging, and to reduce energy expenditure, especially during long haul migration. But such activities come at a cost to individuals, for example, the need to share food and the exposure of those positioned on the edge of the school to attack (Krause, 1994). Hence, how group activity has arisen though evolution, where selection acts on the genome of the individual, has been a source of intense debate.

Here we show that collective behavior is stereotypical and can be described as the consequence of time spent in, and transitions between, 17 different states, each lasting around. By clustering cost curve (Supplemental Figure 8b), we found these were the fewest number of states needed to define the full repertoire of collective actions for wild-type and mutant fish.

We found that single gene changes can modify collective activity in interpretable manner, using the same states. In other words, mutations did not add states. Mutation in the mitochondrial membrane peptidase 2-like gene *immp2l* increases the tendency to aggregate in tight cohesion, and caused the loss of two highly dispersed states. Mutation in the sodium channel Nav1.1 gene *scn1lab* appears to prevent any group interaction, abolishing cohesion and polarization. Mutation in the adrenergic receptor gene *adra1aa* reduces the transitions between states, and causes the animals often to freeze. Freezing and tight cohesion both are seen as responses to threat and conspecific injury (Chicoli & Paley, 2016; Faustino, Tacao-Monteiro, & Oliveira, 2017; Rennekamp et al., 2016).

It is noticeable that these effects correspond to some of the forces (repulsion, attraction, and spontaneity) applied to individual particles in mathematical models to generate coordinated movements of groups (Katz et al., 2011; Krause et al., 1999; Li & Xiao, 2010; Sumpter, 2010). (We have not seen, so far, mutations with selective effects on polarization, a fourth such force). In these models, individual fish need only to manifest these forces between immediate neighbors for the larger scale behavior to emerge as a consequence.

One might argue that widespread changes in neural function could affect a range of behaviors. Although we cannot rule out subtle changes in these parameters, these mutations did not dramatically affect vision or motor function. Of course, other sensory cues might be changed, such as pheromones, so critical to mating (Spence et al., 2008; Yabuki et al., 2016) and to aggregation and dispersion of *C. elegans* (Greene, Dobosiewicz, Butcher, McGrath, & Bargmann, 2016). However, mating remains intact, in terms both of the ornate dance and number of fertilized eggs produced, and we noted no deficiencies in feeding. We also recognize that different environments will elicit distinct behaviors, as will other variables, such as numbers of fish participating (Tunstrom et al., 2013; Yates et al., 2009) and familiarity of the fish with the test chamber. In our studies, the assay was done on fish which had never experienced the test chamber before.

To our knowledge these are the first genes shown to regulate group collective behavior in vertebrates. Some of these have been analyzed for effects in mice, but for other attributes. A mutation in the Inner Mitochondrial Membrane Peptidase 2-Like Gene (*immp2l*) reduces fertility (Lu et al., 2008). One where there is some similarity is haploinsufficiency of the voltage-gated sodium channel, Nav1.1 (Han et al., 2012), which causes hyperactivity and loss of preference for another mouse over empty chambers and curiosity about a novel mouse.

We selected these genes for mutation because they have been reported, with differing degrees of certainty, to have genetic association with autism and schizophrenia, disorders marked by impairments in social interaction. Rare genetic mutations of the IMMP2L gene have been identified in Tourette syndrome and autism spectrum disorders (ASD), and this gene is in the 7q31 translocation region of a Tourette syndrome cohort (Bartnik et al., 2012; Petek et al., 2001). Haploinsufficiency of Nav1.1 in human causes Dravet syndrome, with autism proceeded in infancy by seizures. The gene for the adrenergic receptor ADRA1A is in a large genomic region associated with autism, but no behavioral role has previously been speculated.

Of course, whether these mutations are good models for behavioral deficits in humans remains to be seen. Heterozygous mutation of Nav1.1 in humans, fish, and mice all reduce social interactions. But with the exception of Nav1.1, we have evaluated homozygous effects of mutations, while in humans the mutations are studied in patients with heterozygous deficiencies. To the extent there is duplication of function that accompanies the duplication of many genes in teleost compared to humans or mice (Meyer & Schartl, 1999; Sztal, McKaige, Williams, Ruparelia, & Bryson-Richardson, 2018) certain effects may be compensated. In fact, adra1a is split into adra1aa and adra1ab, knockout of each which causes enhanced freezing, suggesting that there may be duplication of function and compensation. This may also explain why certain candidate autism genes with striking behavioral disturbances when knocked out in mice, such as dlg4a and shank3b (Feyder et al., 2010; Peca et al., 2011), have less striking phenotypes in the fish. Additionally, our assays observe only a small biopsy of potential collective activities under specific controlled circumstances. Nonetheless, zebrafish are amenable to chemical screens for restoration of mutant to normal phenotype (Peterson et al., 2004),

It seems remarkable that some of the genes we discovered here to regulate collective behavior of fish also have been proposed to affect social behavior of humans. This suggests that, perhaps, the role of genes for social behavior, writ large, is conserved through evolution, of course playing out distinctively in different species. The relative convenience of the fish for therapeutic compound screening (Peterson et al., 2004), may be used to advantage in phenotypic screens for reparative agents for disorders such as autism and schizophrenia.

## Materials and methods

### Animals

Zebrafish (*Danio rerio*; AB strain) were housed with mixed gender in 3L tanks on a recirculating Aquatic Habitats facility (Custom design, Pentair, USA). The fish were kept at a density of 12 fish per 3L tank. Facility water temperature was kept at 28±0.5°C, and water sourced from deionized water conditioned with sodium bicarbonate (Catalog #SC12-Pentair, USA) and Instant Ocean sea salt (Catalog #IS160-Pentair, USA) to a pH of 7.2±0.5 and conductivity of 420±50 μS. Fish were maintained on a 14-hour light/10-hour dark cycle with light turning on at 07:00 AM. Fish were fed a diet of brine shrimp (Catalog #BSEPCA-Brine Shrimp Direct, USA) twice daily and supplemented with flake fish food (Tetramin Catalog# 98525-Pentair, USA) every other day. All animals were maintained and procedures were performed in accordance with the Institutional Animal Care and Use Committees (IACUC) of NIBR.

### Genome-wide CRISPR-Cas9 sgRNA design

Because the current work was carried out in the AB strain and the public genome assembly (GRCz11) is based on TU strain, we established a new genome assembly for the AB strain (unpublished, sequences available upon request). For designing CRISPR/Cas9 sgRNAs, we re-trained sgRNA-efficiency models using Random Forest and Naïve Bayes methods based on previously published sgRNAs. 1280 sgRNAs sequences and their editing efficiencies were obtained from previous study in TU-strain by Moreno-Mateos, M. A. et al., (Moreno-Mateos et al., 2015), and mapped to the AB genomic sequence. About 150 features were used to train the two Machine Learning models, Random Forest and Naïve Bayes, in classification mode, including genomic strand of sgRNA, GC%, identity of ±4 bp of sgRNA targeting sequence, thermodynamics parameter, ΔG of sgRNA-genomic DNA heteroduplex for sliding windows of different sizes (Sugimoto, 1995), the free-energy of stem-loops 1, 2, and 3, tetraloop, repeat-anti repeat and linker structures of the full-length sgRNA predicted by UNA-fold (Hiroshi Nishimasu, 2014; Markham & Zuker, 2008) and etc. A training accuracy of > 0.7 was achieved by both models, and the efficacies of all sgRNAs in the AB genomic sequence was predicted. Guide RNAs with high predicted efficacy were selected to target the 5′ end of the coding exons or Pfam domains of the genes in current study (for the full gene list, citations for disease indications, please see Supplemental. Table 1 and for the guide RNA sequences used for CRISPR-cas9, please see Supplemental. Table 2).

### Micro-injections, CRISPR/Cas9 mutant founder identification

sgRNAs were synthesized using T7 *in vitro* transcription using the MEGAshortscript™ T7 Transcription Kit (ThermoFisher, AM1354). Cas9/sgRNAs were co-injected into 1-cell stage fertilized zebrafish embryos. Conditions were optimized to maximize the CRISPR efficiency in our settings. High indel rates (>90%) were usually observed in fully developed embryos injected with 500 ng/μL Cas9 protein (PNABio, CP01) and 125 ng/μL sgRNA purified with MEGAclear™-96 Transcription Clean-Up Kit (ThermoFisher, AM1909).

Indel rate in the injected zebrafish were measured using NGS at 2 days post fertilization (dpf). PCR were performed on the genomic DNA extracted from zebrafish larvae using the HotSHOT method(Meeker, Hutchinson, Ho, & Trede, 2007), and NGS libraries for PCR products were generated using Nextera DNA Library Preparation Kit (Illumina, FC-121-1031). Subsequently, Nextera libraries were sequenced on Illumina MiSeq, and > avg. 2000 reads for each amplicon were obtained. Sequencing reads were mapped to the reference sequence using BLAT (Kent, 2002), and indels were extracted from the .psl file using a bioinformatic pipeline developed in-house (available upon request). The gene editing efficiency in the injected embryos were calculated as the maximum indel rate within the ±30 bp regions of PAM sites.

We usually sequence five larvae per CRISPR injected clutch to confirm the gene editing efficiency. The rest fish of the confirmed clutch were raised to adulthood and crossed with wild-type AB fish, and founder fish carrying the desired frameshift mutation were screened from the F1 generation. F1 heterozygotes were inter-crossed, and homozygotes, if viable, were identified and inter-crossed again to obtain sufficient gene knockout fish for behavioral assays. Each CRISPR line was at least an F2 stable line before being run in any assay. All founders and homozygote identification were carried out using fin-clipping PCR, and sequencing the PCR product using NGS as described above. The indel sequence of homozygous mutations and heterozygous mutations that could not be bred to homozygous (*scn1lab* and *slc18a2*) are illustrated in Supplementary File 1, and also the predicted protein sequence aligned with that of the wild-type fish.

### Experiment setup for behavioral assays

Attention was paid to ensure fish gender balance and matching of size and age, and to conduct experiments at similar times of days and feeding cycle, and to monitor by video without human presence. All behavioral rooms were fed with water directly from the main fish facility and room temperature and light cycle was consistent with the main facility. The behavioral setups consisted of 20” diameter acrylic circular arena (Custom Design: Acrylic Tank – Clear - 20” OD x 19.25” ID x 8” height – Open Top, Plastic Supply, Inc., USA) filled to a depth of 1 3/4” (~9 L total volume) with system water fed directly from the main housing unit to ensure all water parameters were identical to the housing conditions. The circular arenas were coated on the outside (I00810, Frosted Glass Finish, Krylon, USA) to prevent the fish from being able to see outside of the arenas but to allow IR light transfer. Underneath the tanks were 24 inch adjustable IR panels (with 940 nm IR LEDs made by Shenzhen VICO). Basler Ace 2040-90um Near Infrared (NIR) cameras (Order#-106541, Graftek Imaging, USA) were mounted 23 inches above the arena to collect a dorsal view of the fish. Infrared long pass filters (Midopt LP780-62, Graftek Imaging, USA) were attached to the lens (Scheider Cinegon 1.9, Graftek Imaging, USA) and were set to an aperture of 6 (Supplemental Figure 9a). All trials were recorded after 10 minutes habituation to allow recovery from any stress due to netting. Briefly, shoaling assays were run with six fish (3 males and 3 females), which were randomly combined from multiple tanks of siblings (with the exception of *chd8*−/− which had only male homozygous animals so were paired with 3 AB females) (Supplemental Figure 9c). Each trial was a recording of 30 min at 60 fps. Arenas were rinsed clean with system water at the end of the day and put through a cabinet washer (Type: 9LAV65, IWT Tecniplast Inc., Italy) once a week on a hot water only cycle.

Courtship assays (1 male and 1 female) were all conducted in Aquatic Habitat 2L group mating tanks (Catalog# Breeder2-Pentair, USA) positioned carefully within arenas used for the above shoaling assays (Supplemental Figure 9b). The main arena was still filled with water to ensure all fish’s safety as they infrequently escape the mating tanks, which remain lidless due the need to eliminate optical obstructions. Image size was fitted to exactly cover the two tanks in the overnight assays (Supplemental Figure 9d). Three 30-min videos were recorded for each pair of animals: the first recording was initiated at 16:00 PM while fish were separated by a clear divider; the second recording at 22:00 PM while the lights were off and divider still in place; the third recording was done at 08:00 AM (the following day) with the divider removed. After each recording, spawning records were documented and confirmed only if more than 10 embryos survive till 1 dpf with normal course. All mating tanks used in assays were washed daily in under counter washers (Type: GG05/Model PG 8583; Miele AG, Germany).

### Automated data collection

We used a virtual instrument console designed within LabVIEW (National Instruments, USA) to control all cameras. The acquisition software was designed with four prioritized functions: 1) Saving the recorded video; 2) Logging all relevant metadata accurately and automatically; 3) Enabling real time user visualization, verification and modification; 4) Generating associated files to streamline downstream analysis.

Non-default parameters were set within the NI-MAX (Measurement and Automation Explorer) including dimensions and offsets for centering (1880×1880 pixels (offset 84/84) for Shoaling assays, 1000×1000 pixels (offset 420/600) for Courtship assays) and frame rate (60 fps). One workstation (X2D65UT#ABA, Z440, Hewlett Packard Inc., USA) was dedicated for simultaneous recordings of two USB3.0 Basler ACE cameras. The resulting videos which are approximately 360 GB each for Shoaling assays, and 108 GB each for Courtship assays are saved on a local 4×2 TB SSD RAID0 (Samsung EVO 850, B&H Photos Inc., USA). All cameras were named uniquely to allow us to trace back which videos are generated by corresponding setups.

Relevant metadata includes time of trial, user, and fish information (genetic background, age, genetic knockouts, genotype, compound treatment (concentration/duration) and other manipulations), and assay information (duration, fps, number of male and female fish). All software developed in house is available upon request, excluding licenses.

### Data preprocessing

The processing involved building of fingerprint libraries to distinguish fish accurately even after repeated crossings, using locomotion quantifications such as trajectory, angles, speeds, pairwise distances, and convex hull area. (software based on MATLAB 2013a; MathWorks, U.S.A.) inspired by Perez-Escudero *et al.* (Perez-Escudero, Vicente-Page, Hinz, Arganda, & de Polavieja, 2014). In addition, we used supervised behavioral annotation of simple motifs, such as nonsocial turn, dive, rest, and cruise, or social chase, cross, and parallel rotation, with previously trained classifiers as described by Kabra *et al* (Kabra et al., 2013). These processing procedures are integrated into an overarching program that manages data dependencies to automatically start the next stages when ready without needing additional user inputs (scripts available upon request).

### Unsupervised collective behavior state hierarchical structure clustering

For this analysis we used 5 frame epochs. Polarization was determined by the swimming direction angle. For the *i*^*th*^ fish in epoch *n*, this was represented by the tangent angle of the *i*^*th*^ fish’s trajectory from the 1^st^ frame of the *n*^*th*^ epoch to the 1^st^ frame of the *n*+*1*^th^ epoch in the Cartesian coordinates where the trajectory was represented. If the standard deviation of the angles of the six fish in an epoch was lower than the threshold 0.6 radians and the minimum swimming speed of the 6 fish in the epoch faster than 1 inch per second, the epoch was classified as “polarized swimming”; otherwise the epoch will be classified as “unpolarized swimming”.

The pairwise distances among the six fish’s centroids was calculated from the 1^st^ frame to represent the pairwise distances in each epoch. The pairwise distances within each epoch was ranked in a descend order. A principle component analysis (PCA) dimension reduction was applied to the ranked pairwise distance matrices (with a dimension of 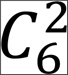 by number of epochs) in polarized swimming and unpolarized swimming epochs, respectively. The top 2 components that capture more than 95% of variance were kept and *fuzzy* C-means clustering was implemented on the top 2 PCA components with *k=9* states, for polarized swimming and *k=8* for unpolarized swimming components, respectively. The flow chart of the unsupervised learning approach we performed is shown in Supplemental Figure 9a. A majority filter is applied to the clustered labels to eliminate cases that only a single epoch has a different label from previous epoch labels and post epoch labels, since there is less interest in super short states with duration less than 1/6 seconds. Change points identified state block duration distribution is shown in Supplemental Figure 9. The mean of block duration is 0.78 seconds with a standard deviation of 0.96 seconds.

A t-distributed stochastic neighbor embedding (t-SNE) (Laurens van der Maaten, 2008) plot was used to visualize the high dimensional data in a two dimensional space for the wild-type fish collective behavior (Supplemental Figure 10a-b). The collective behavior pattern is shown by clickable videos for the selected states.

The *k* value*s* were determined by optimizing the clustering cost (D T Pham, 2005) (Supplemental Figure 9b). Moreover, to increase the resolution of the state quantification, the state number was pushed to *k=50* for both polarized and unpolarized swimming. The overall results remain consistent.

Self-Organizing map (SOM) clustering and k means were applied as orthogonal approaches to *fuzzy C*-means. The performance of SOM and k means is equivalent to *fuzzy C*-means (data not shown). All the quantifications are implemented in Matlab R2014a (Mathworks. Inc, U.S.A.). Figure 1 to 3 and supplementary figure 1, 2, 3, 5, and 10 were plotted in *R 3.4.1*, the rest figures were plotted in Matlab. Silhouette analysis was performed for both polarized and un-polarized swimming to ensure no significant over classification (data not shown). The Matlab and *R* codes to replot all the figures in this manuscript are available upon request.

### Ethogram visualizations

Ethograms are used to represent the different states that occur during collective behaviors. The ethogram indicates the frequency and transition probability with which one state is followed by another state (Anderson & Perona, 2014; Dankert et al., 2009). To represent the relationship among behavioral states and probability distribution of states using the ethogram, two type of analysis were performed. First, we computed the one-step transition probability which is the conditional distribution of the current state given by the previous state (Bishop, 2006). Second, we computed the probability distribution of states which is the frequency of the occurrence. In practice, The ethogram is generated by *igraph R* package (Csardi, 2006). In the ethogram, the edge and vertex are corresponding to the one-step transition probability and probability distribution of states, respectively. The thicker and darker the edge is, the higher transition probability is; and the bigger the vertex is, the higher the state appearance probability is. All attributes (e.g., vertex color, vertex size, arrow size, edge size, and edge curvature, provided in the scripts) were adjusted by parameters of the package to generate current figures.

### Kullback-Leibler divergence

Kullback-Leibler divergence (relative entropy) (Kullback & Leibler, 1951) is a mathematical method to measure the differences of two probability distributions. The difference *D* is a function of the two discrete probability distributions *P* and *Q* as shown in the following equation (1):

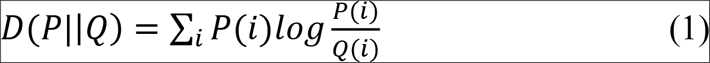

Kullback-Leibler divergence was applied to the polarized and unpolarized state usages, respectively. The differences between each mutation and wild type were ranked in descending order (Supplementary File 2) and the top ranked mutations were selected.

### Supervised behavioral phenotype category classification

#### Feature extraction

State usage, transition matrix and swimming speed in each trial are employed as features (also called measurements or attributes), including the averaged aggregation (unpolarized) state usages, dispersion (unpolarized) state usages, schooling (polarized) state usages, overall state usages, and transition matrix. Speed features include mean speed of all the 6 fish each trial in the last 10 minutes of the recording, number of stop epochs from all 6 fish in each trial. PCA then is implemented to reduce the dimensionality of the feature space. Top 12 PCs are kept capturing above 95% of the variance. In addition, an alternative feature pool is tested, including only swimming speed distribution in each unpolarized state from all the trials. PCA dimensional reduction is followed and 3 top PCs are selected to capture 95% variance of the data.

#### Support vector machine

The classifier we implement is the support vector machine (SVM) (Vapnick, 1995), which is widely used for pattern classification (Ma, Randolph, & Drish, 2001). For the classical binary formulation of SVM classification, the output of an SVM classifier yields:

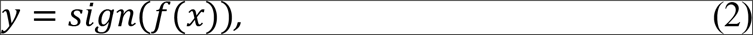

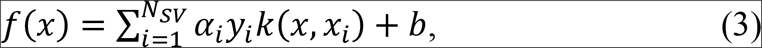

where *α_*i*_* are the Lagrange multipliers from solving the quadratic optimization problem, *y_*i*_* are the class targets, *x*_*i*_ are the input data points, *k*(*x*, *x*_*i*_)is a kernel function (Chang, 2011), *b* is the bias, and *N*_*sv*_ is the number of support vectors. Linear kernel is used here since it is more capable of avoiding possible overfitting in the classification than other kernels (Keerthi & Lin, 2003).

To extend a binary classification to a multi-class classification problem such as the problem at hand, several binary SVM classifiers need to be utilized. One-against-one classification method trains *k*(*k*−*1*)/2 binary classifiers (where *k* is the number of classes) to solve *k* class classification problem. During prediction the class with the most votes becomes the winner. One-against-one outperforms other multi-class SVM classification methods (Hsu & Lin, 2002) and is the default choice in the libSVM package (Chang, 2011) that has been utilized in this study.

#### Leave-one-out

The Leave-One-Out (LOO) (Efron, 1993) validation method is used to evaluate the performance of the classifiers described in this study. Random one trial from each category is taken as the testing data and the remaining as the training data to train SVM multi-class classifiers. This procedure was applied for 100 times (most trial numbers are less than 10) by randomly choosing training and testing trials. All the test results are then combined into a cumulative confusion matrix.

## Acknowledgements

We thank Ricardo Dolmetsch, Rainer Friedrich, Joseph Loureiro, Mark Borowsky, Brant Peterson, Gerald Sun, Lingling Shen, Ajeet Singh, Vibhas Aravamuthan, Stephen Litster, Guangliang Wang, and Jian Fang for discussion. We also want to thank Gerlinde Wussler, Kara Moloney, Meghan Aguirre, Franki Vetrano-Olsen, KarenJ Lee, Haley Clark, Johnathan Tobin, and Joseph Beaton for their support in the fish facility, also Stacey Gearin, Stephanie Wiessner, and Jessica Garver for CRISPR injection work, Caroline Fawcett for fish database management; thank Michael Steeves and Michael Derby for their help in computation resource; thank Michael Paolucci and Aaron Bickel for their help in hardware customizing; thank Andrea Schwarz and Lauren Goldfinger for project coordination. X.X. is supported by NIBR postdoctoral scholar program. Funding is provided by the Novartis Institutes for Biomedical Research (NIBR).

---

**Supplemental Figure 1.**
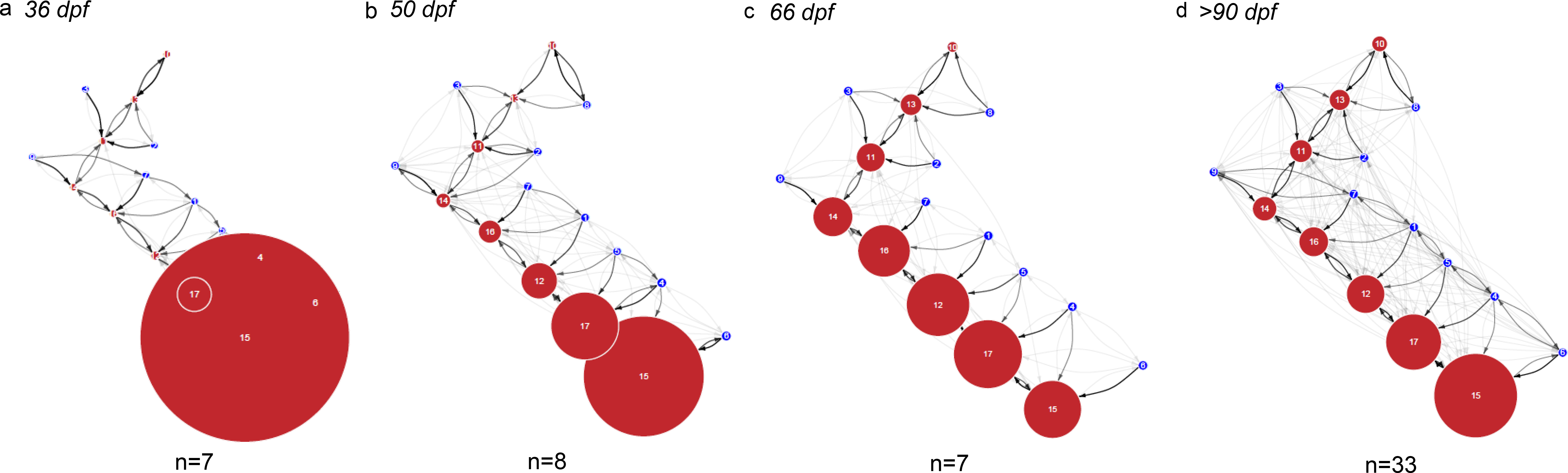
Development of collective behavior, showing ethograms of wild-type fish (a) 36 dpf (note that most of the usage concentrates in state 15, the tightest cohesion state, and that the dispersed schooling state 8 is missing; (b) 50 dpf (tight cohesion states 15 and 17 are still the majority of state usage); (c) 66 dpf (the collective behavior state usage is similar to adult fish, although the transitions between states are less dense); (d) adult fish. Collective behavior is not mature until the fish are about 3 months old.

**Supplemental Figure 2.**
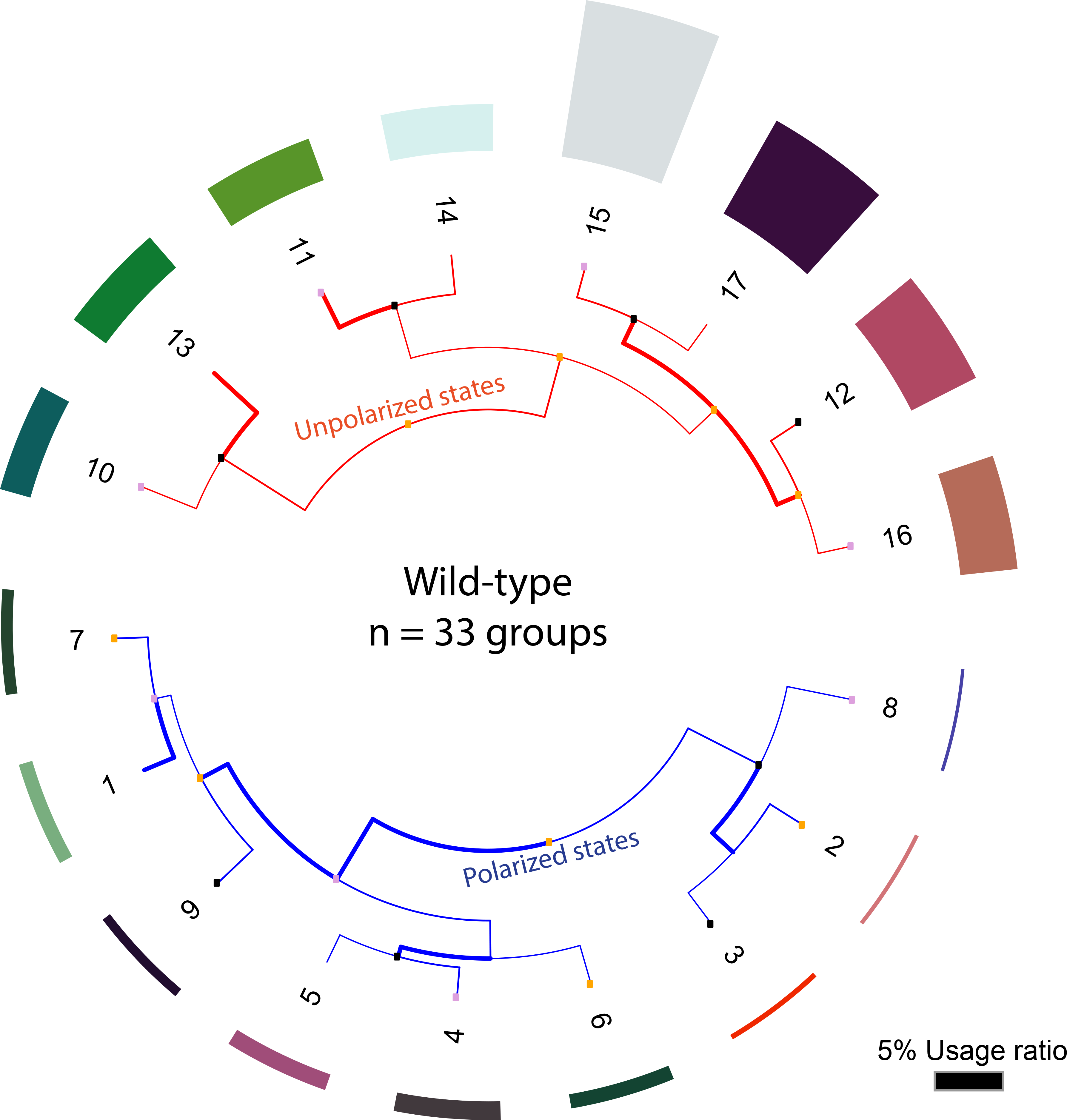
Dendrogram of wild-type fish collective behavioral states defined using hierarchical clustering.

**Supplemental Figure 3.**
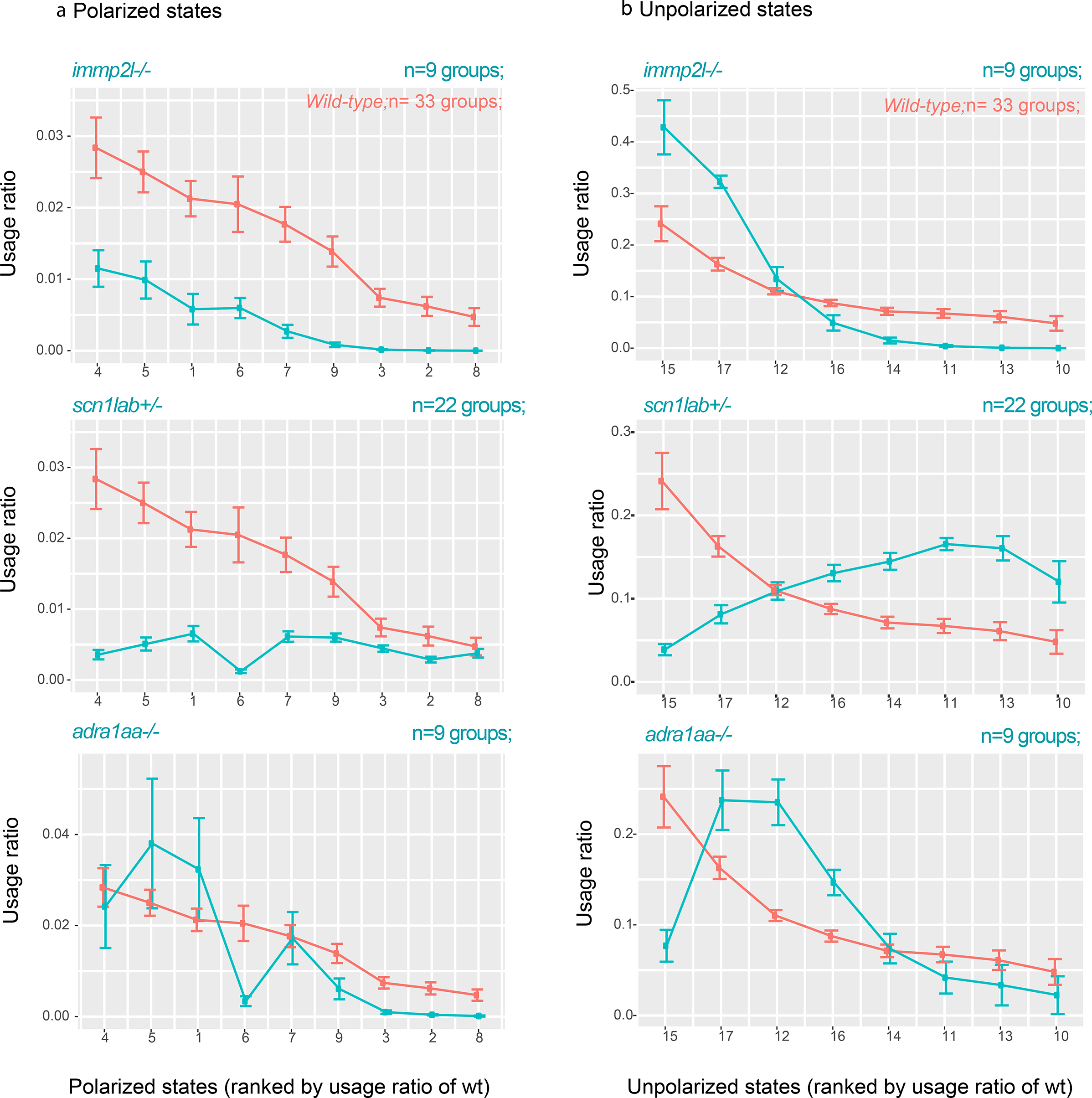
Usage ratios of mutant states. Usage ratios of all (a) polarized and (b) unpolarized states are compared to corresponding states in wild-type (error bars indicate ±SEM). The usage ratios are compatible with observations described in the text for the ethograms of Figure 2, and with significance test in Supplemental Figure 6.

**Supplemental Figure 4.**
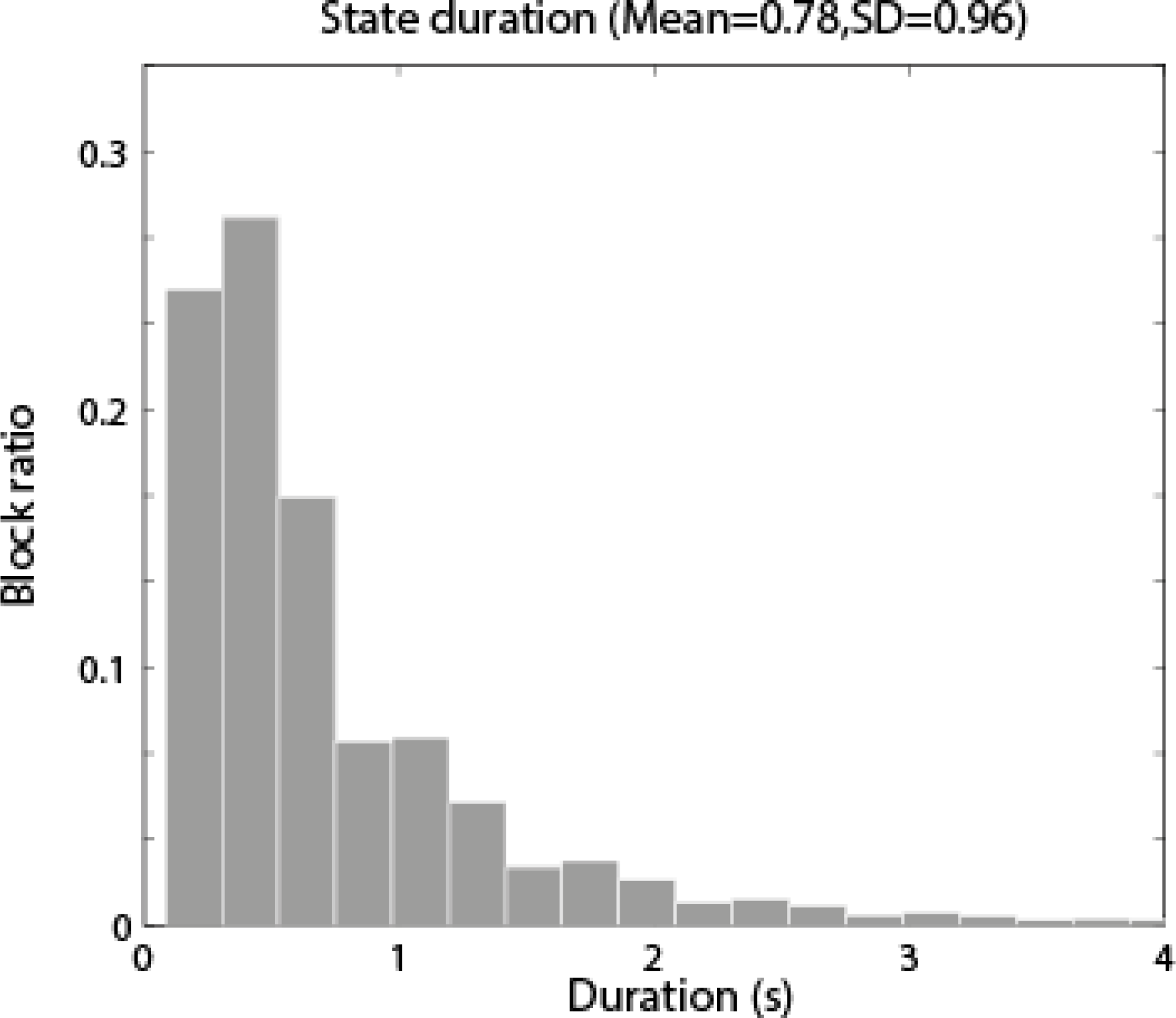
Block duration of states (Fewer than 1.2% last longer than 4 seconds). The mean is 0.78±0.96 seconds.

**Supplemental Figure 5.**
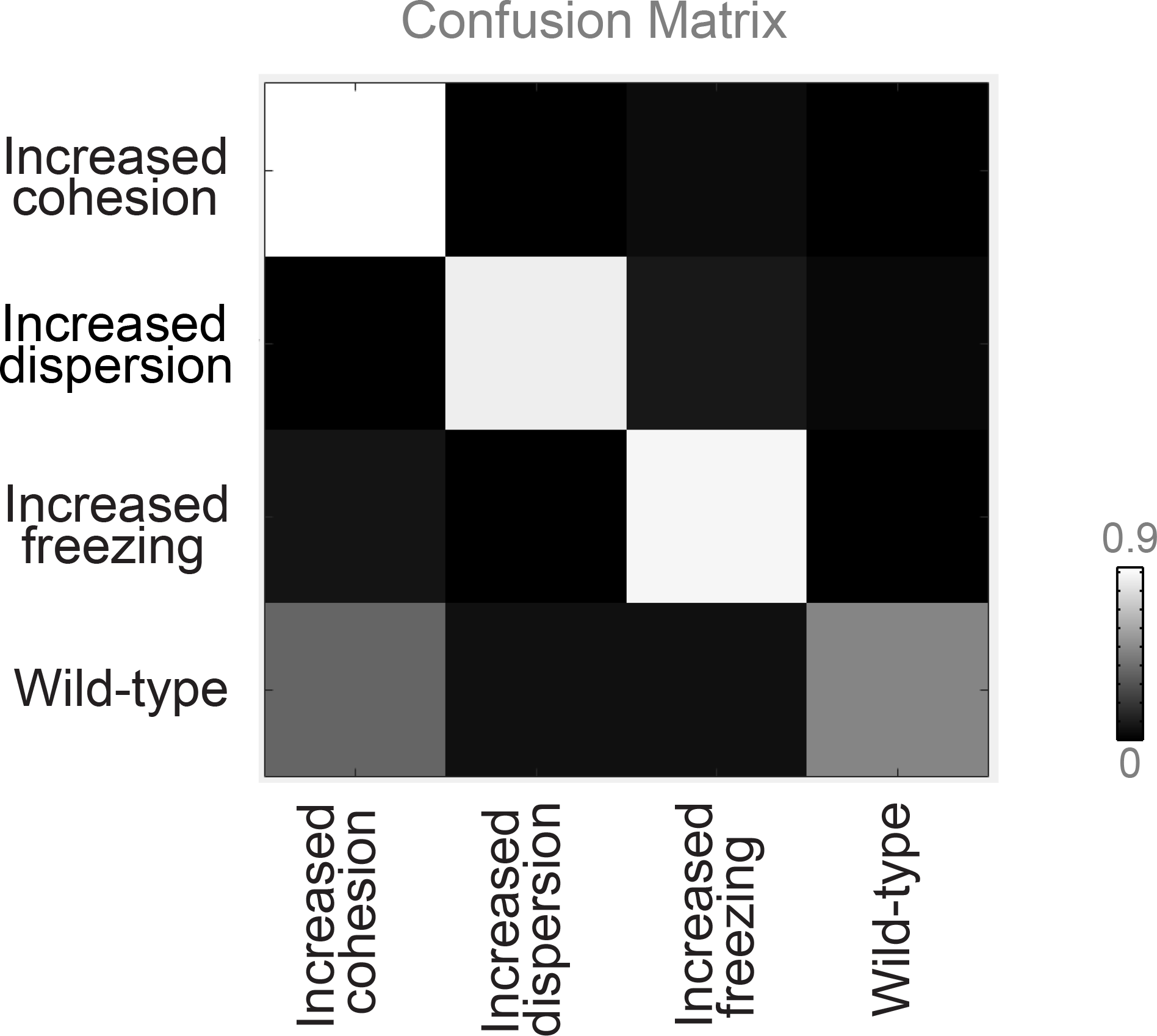
Cross-validation by multi-class Support Vector Machines confirms robust phenotypes revealed in highlighted mutants. Confusion matrix computed using features, including state usage ratios, transitions, *etc. immp2l*−/− (n=9 groups, Prediction Accuracy (PA) = 0.95; Leave-One-Out); *scn1lab*+/− (n=22 groups, PA= 0.88; Leave-One-Out); *adra1aa*−/− (n=8 groups, PA= 0.92; Leave-One-Out).

**Supplemental Figure 6.**
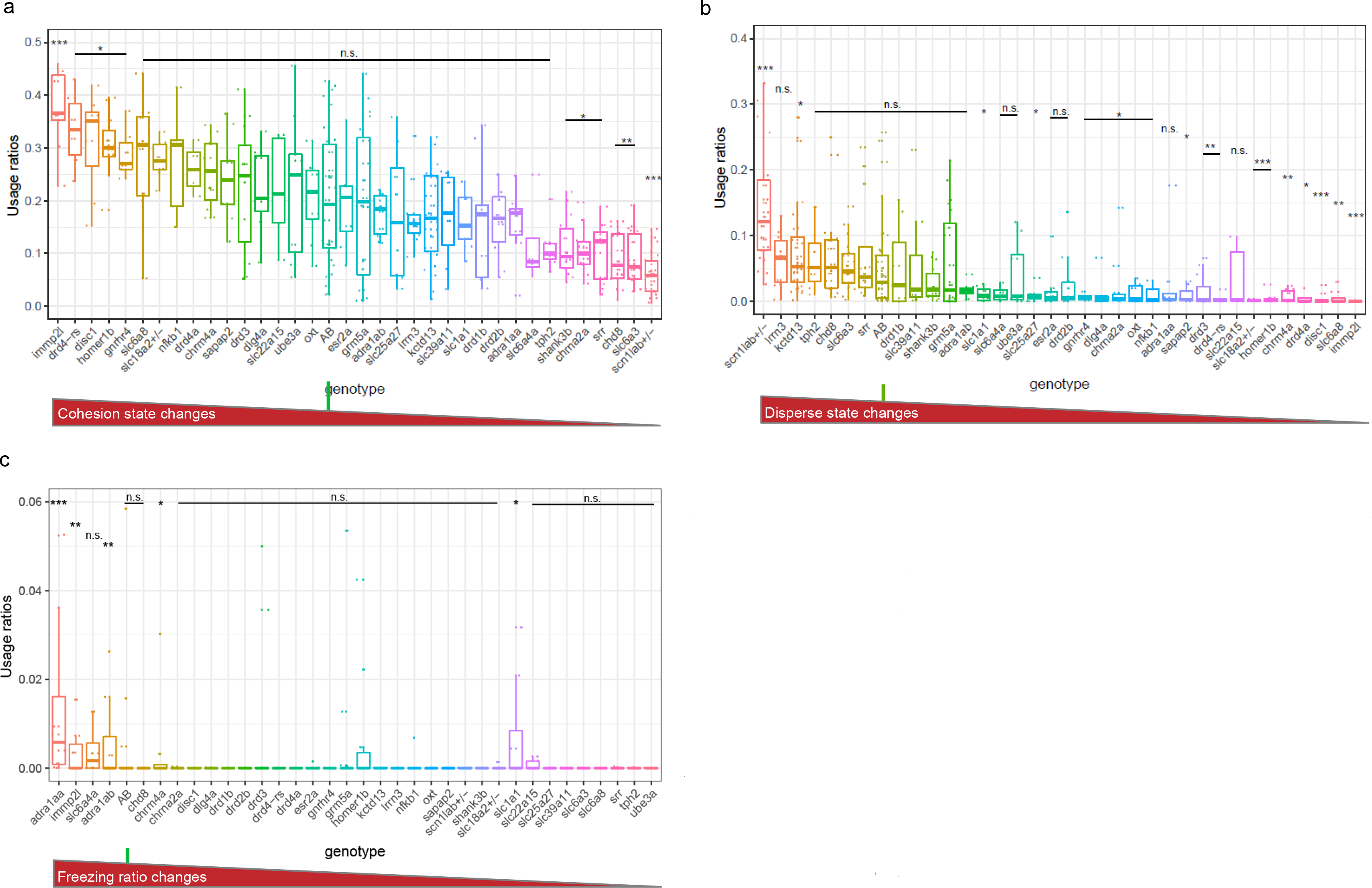
Ranked usage ratios of fish having each mutation, compared to: (a) *immp2l−/−‘s* tendency to cohesion; using the usage ratios of the top two tight cohesion sates which are significantly enhanced in *immp2l*−/− compared to wild-type (n=9 groups, p<0.001; Mann-Whitney U-test). By this comparison, significantly more cohesion is also demonstrated by *drd4-rs*−/− (n=6 groups; p<0.05; Mann-Whitney U-test), *disc1*−/− (n=9 groups; p<0.05; Mann-Whitney U-test), *homer1b*−/− (n=11 groups; p<0.05; Mann-Whitney U-test), and *gnrhr4*−/− (n=9 groups; p<0.05; Mann-Whitney U-test). (b) *scn1lab+/−‘s* tendency to disperse, using the usage ratios of the top two dispersed states significantly enhanced in *scn1lab*+/− compared to wild-type (n=22 groups; p<0.001; Mann-Whitney U-test). By this comparison significantly more dispersion also is shown by *kctd13*−/− (n=28 groups; p<0.05; Mann-Whitney U-test). (c) *adra1aa−/−‘s* tendency to freeze; the ratio is calculated from the duration of simultaneous freezing of 3 fish in the last 10 minutes of the recording (n=8 groups; p<0.001; Mann-Whitney U-test). This tendency also is shown by, *immp2l*−/− (n=9 groups, p<0.01; Mann-Whitney U-test) and *adra1ab*−/− (n=10 groups, p<0.01; Mann-Whitney U-test).

**Supplemental Figure 7.**
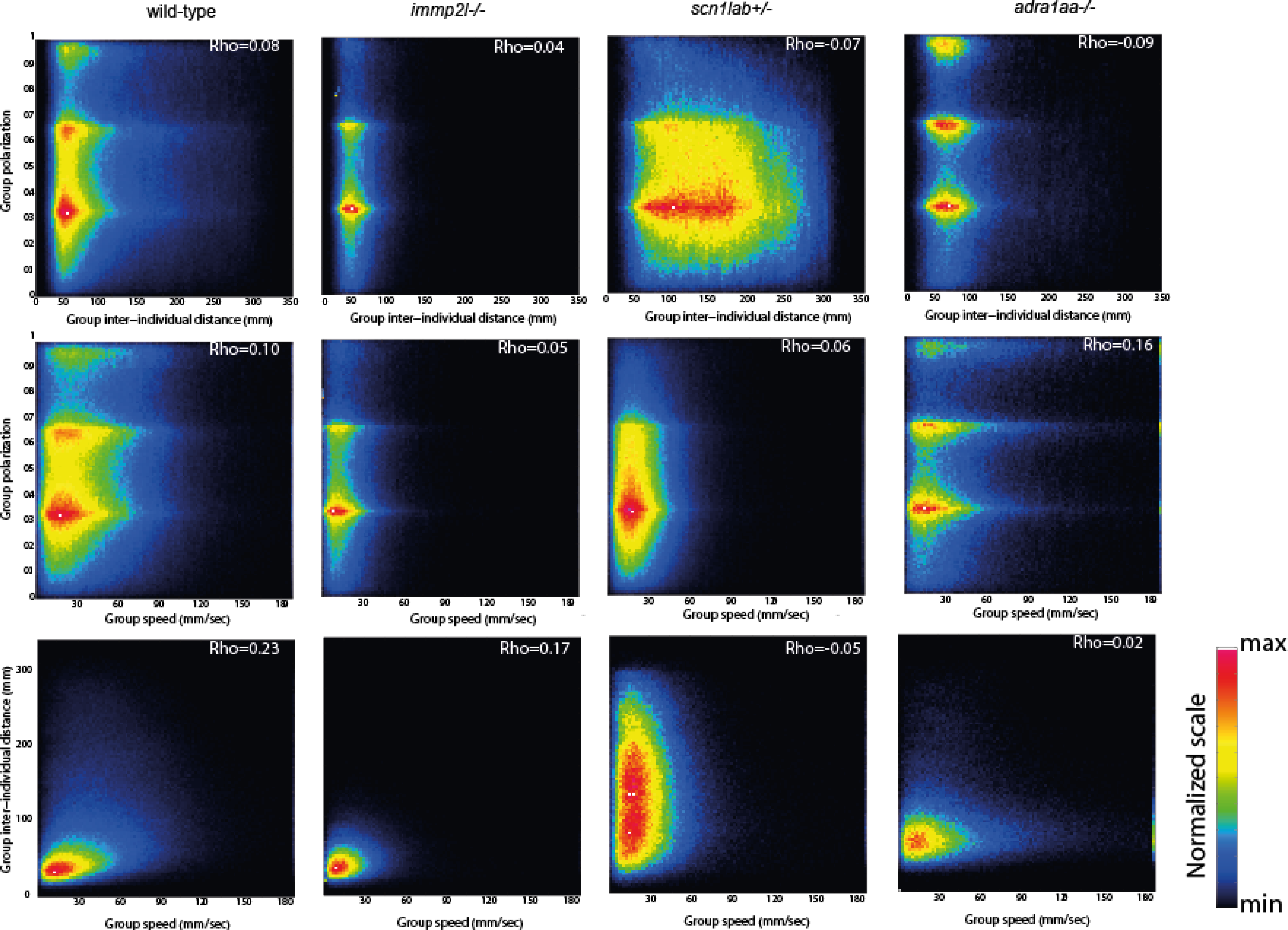
Lack of relationship between group polarization and inter-individual distance (top panels) and speed (middle panels) or between inter-individual distance and speed (bottom panels), for the mutations discussed. The colors indicate the number of frames in each bin (100 bins in each dimension).

**Supplemental Figure 8.**
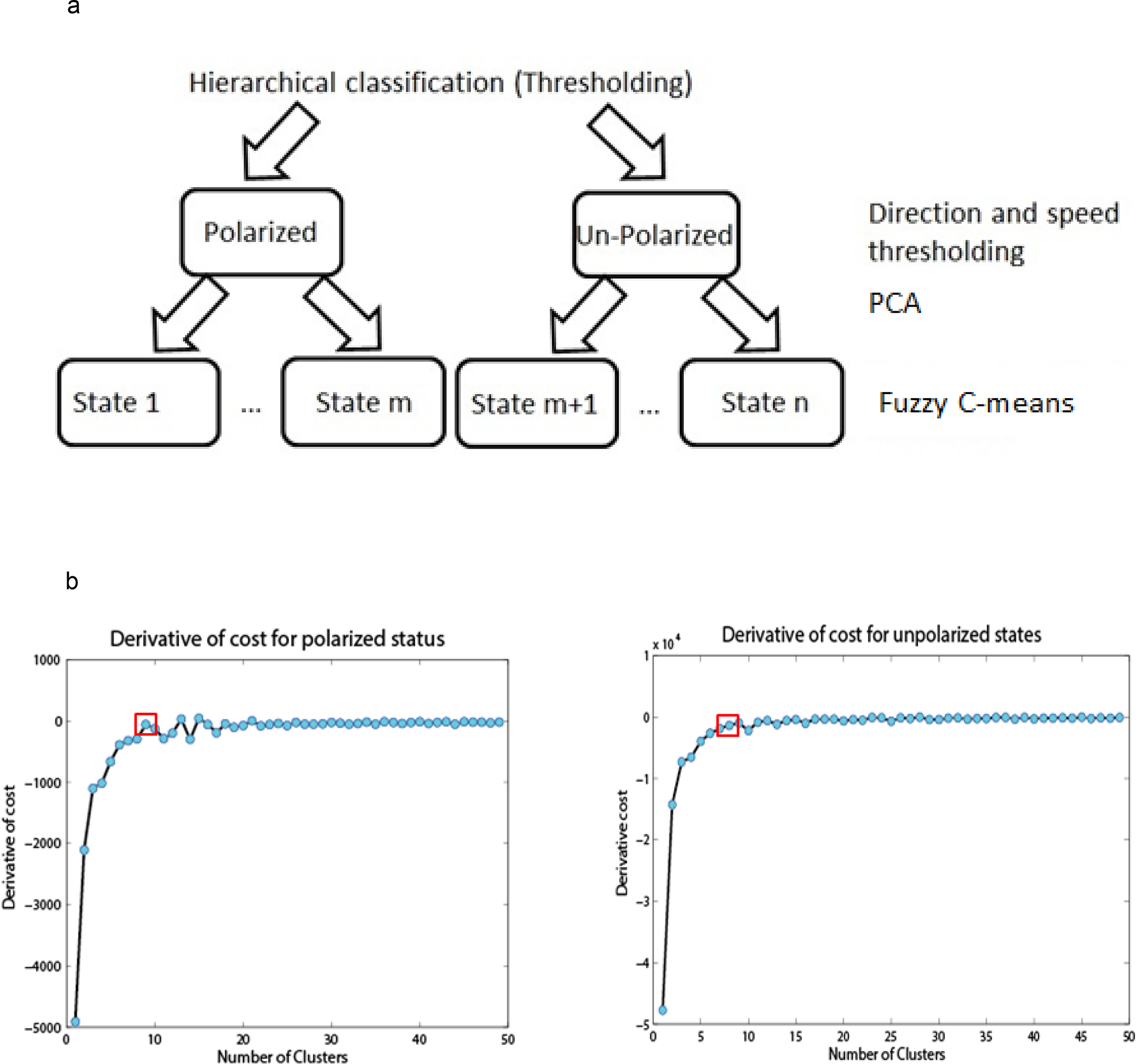
(a) Hierarchical clustering flow chart of collective behavior unsupervised learning (b) The derivative of cost of the clustering for porlarized and unporlarized states, respectively. The red square boxed mark the cluster number we chose.

**Supplemental Figure 9.**
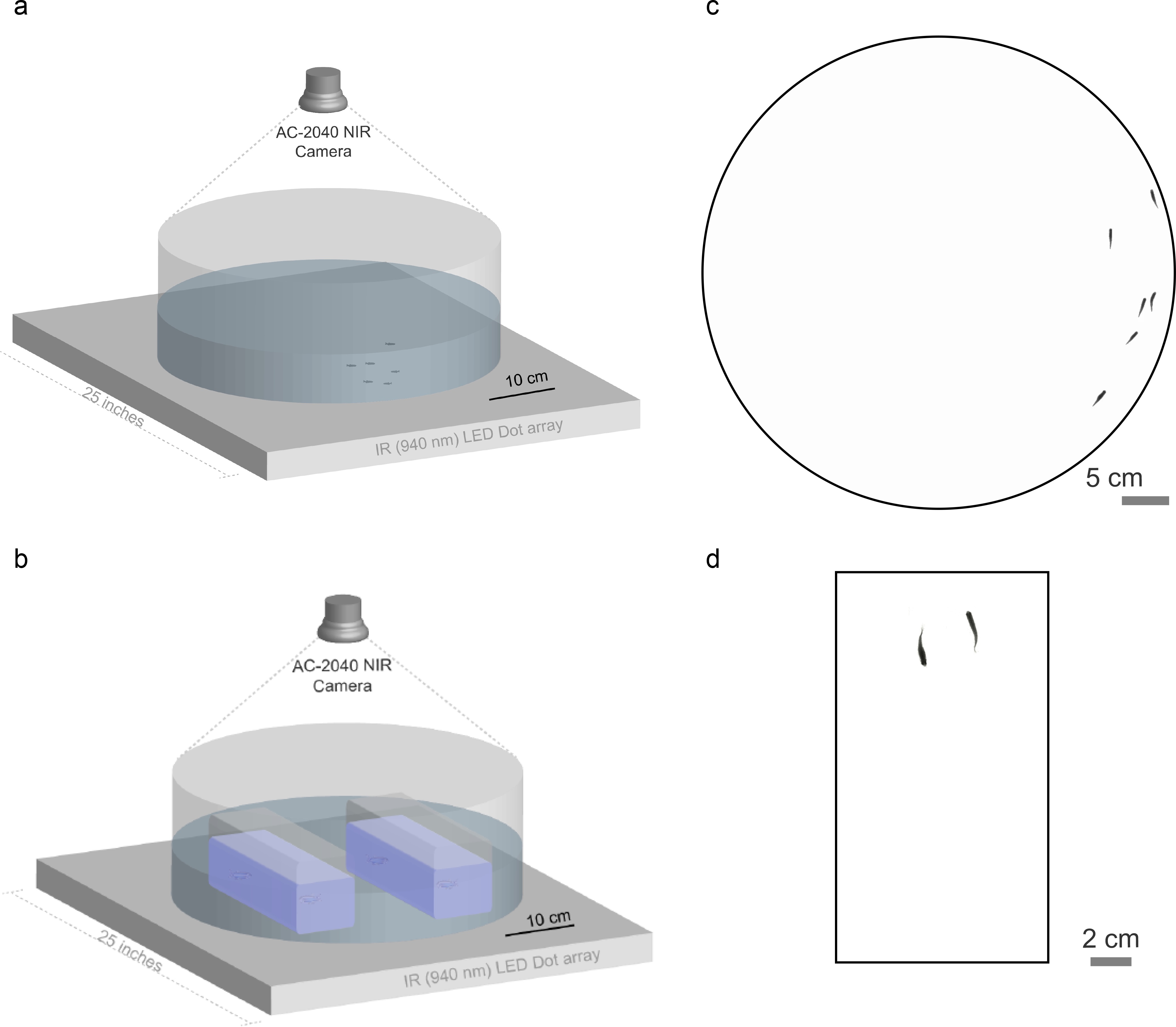
Behavioral assay setup (a) Collective behavior assays and (b) courtship behavior assays were recorded from Top View, with bottom IR (940 nm) illumination. Examples of background removed images are shown for (c) collective behavior assays and (d) courtship behavior assays.

**Supplemental Figure 10.**
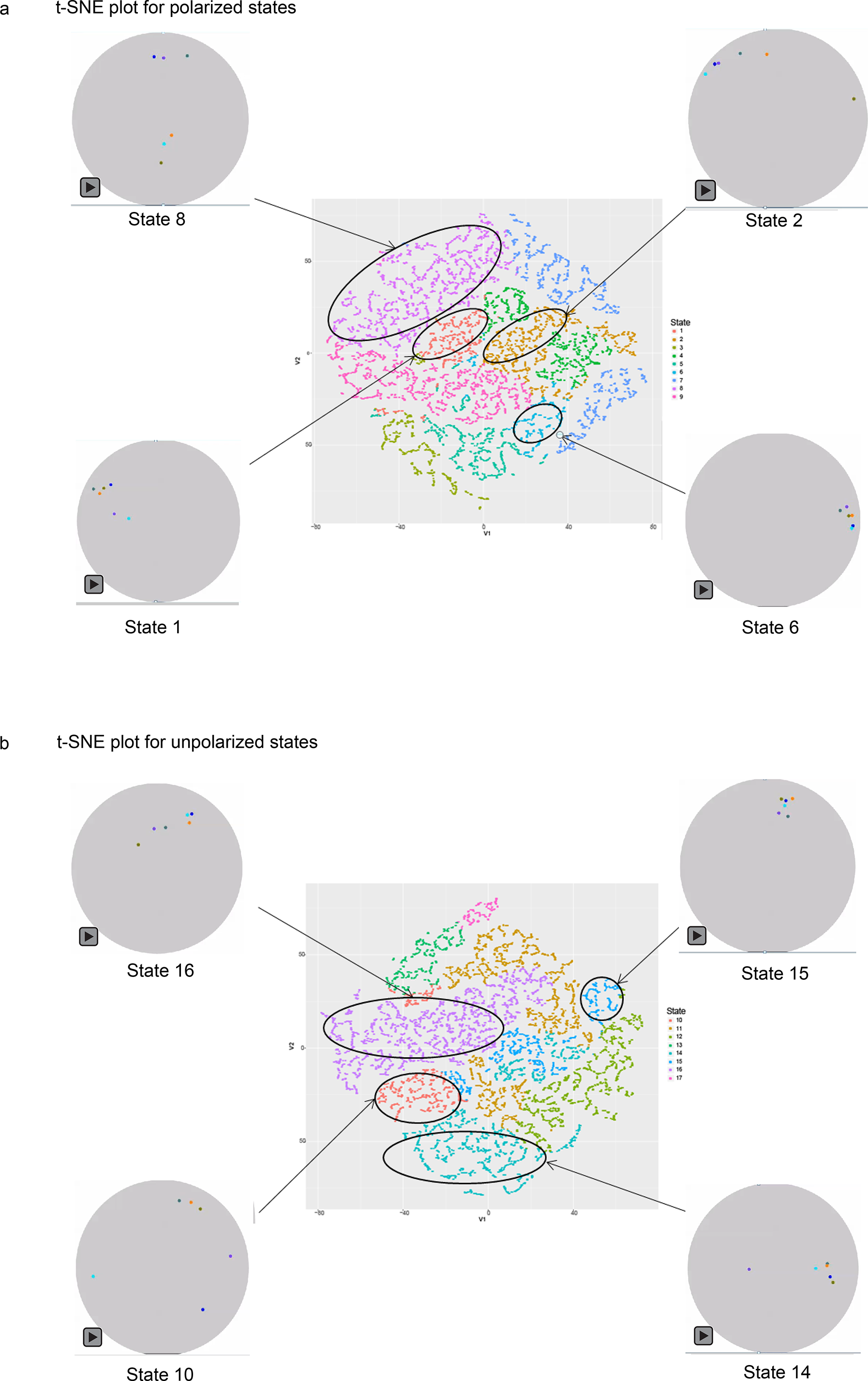
t-SNE plots for (a) 9 polarized collective behavior states and (b) 8 unpolarized collective behavior states. Different colors in the plot represent different state clouds. Each plot shows the clustering of 5000 epochs (1 epoch=5 frames of recorded images) from polarized and unpolarized states, respectively. Videos show the collective behavior pattern of each corresponding state and can be accessed by clicking where shown. The colored dots in each video represent the centroid location of each individual fish.

**Supplemental Table 1.**
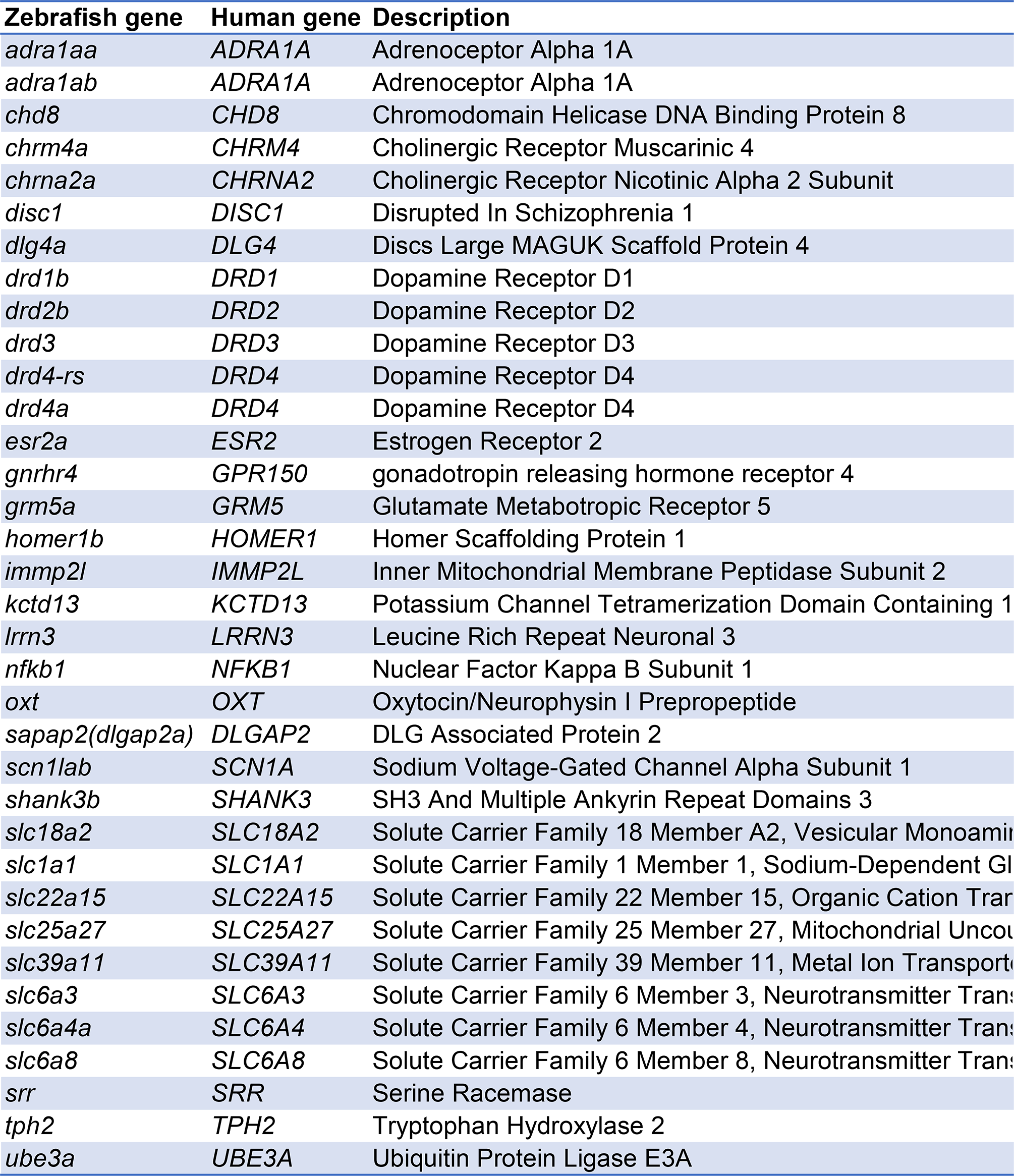
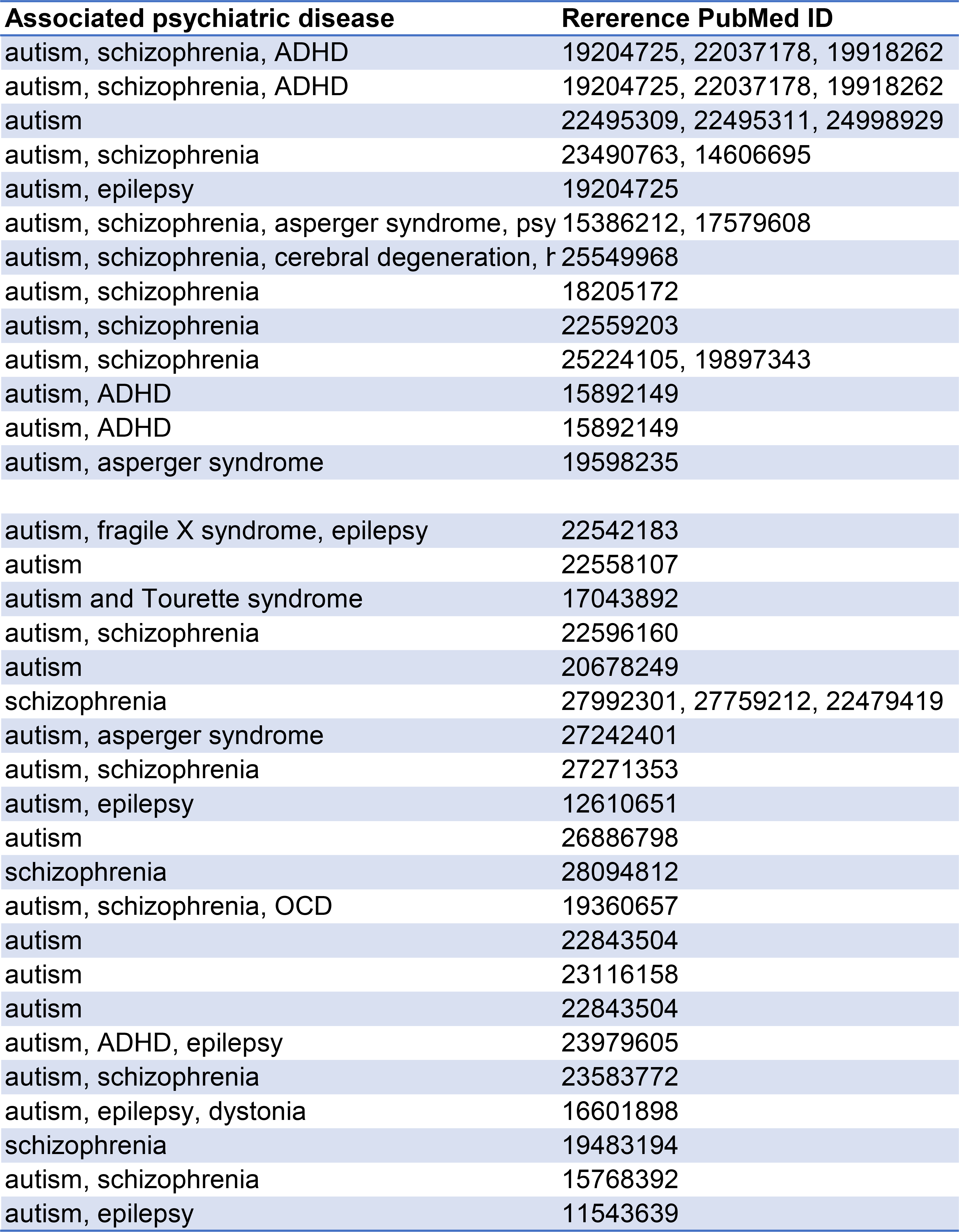

**Supplemental Table 2.**
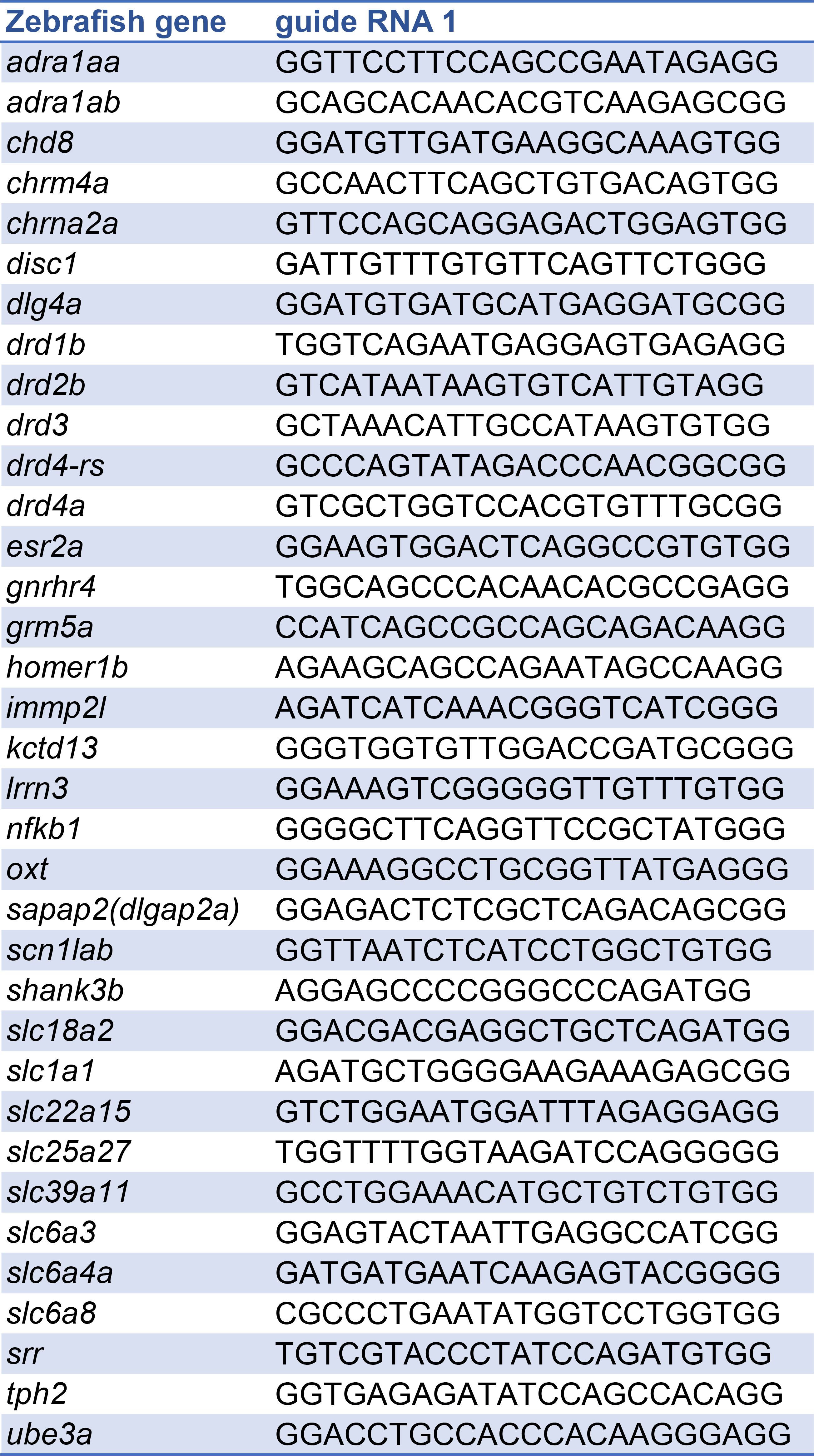
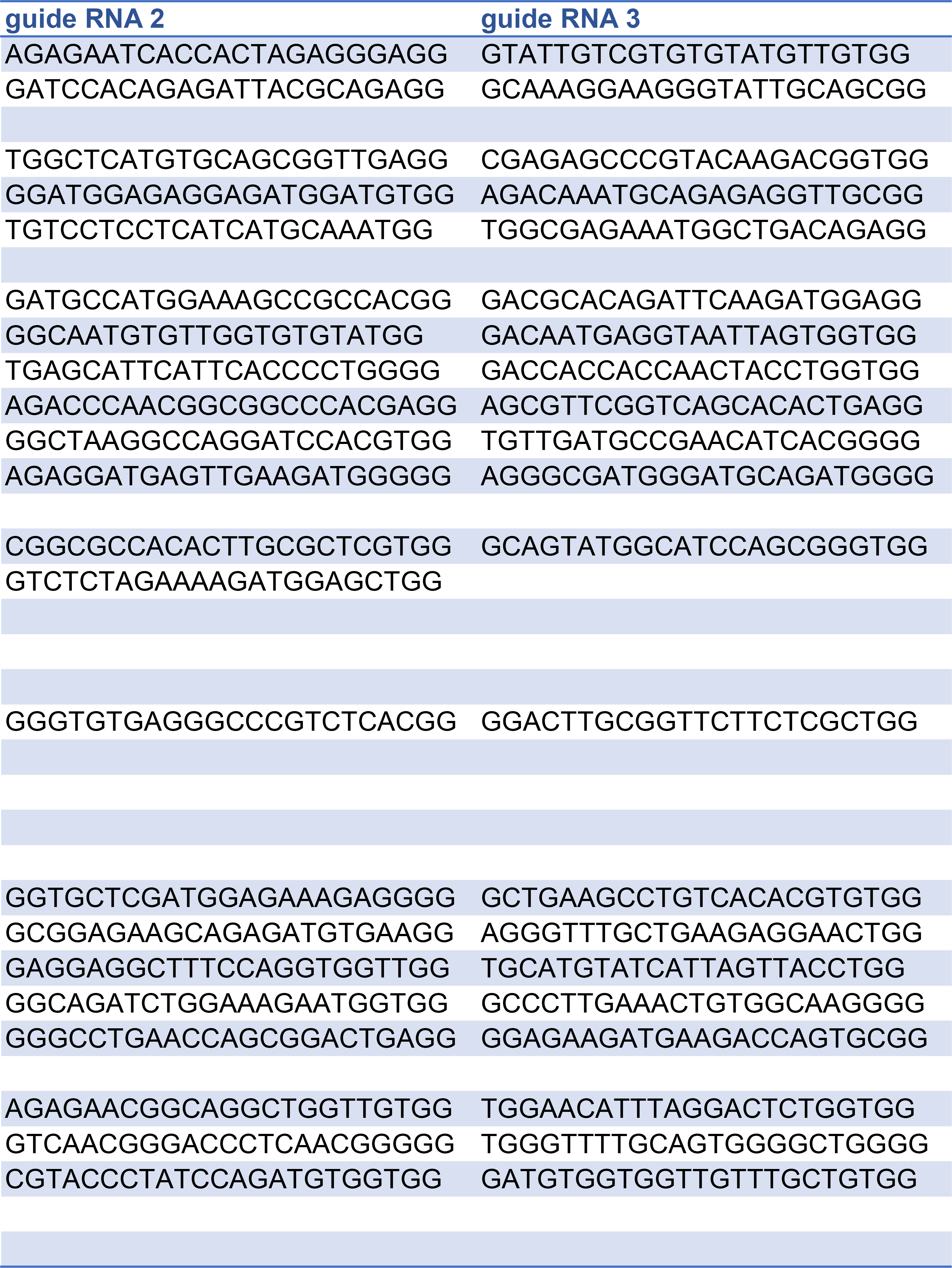
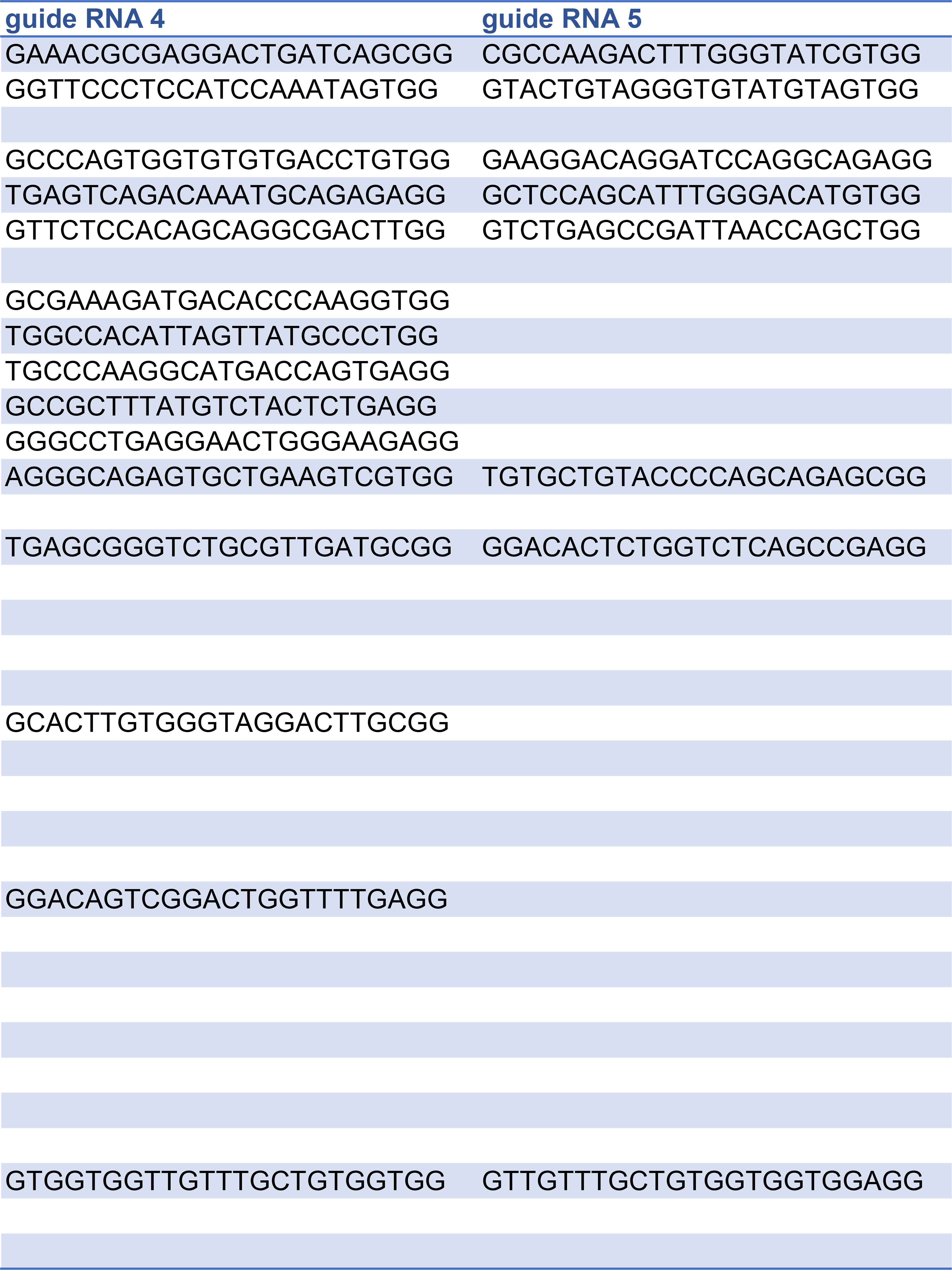
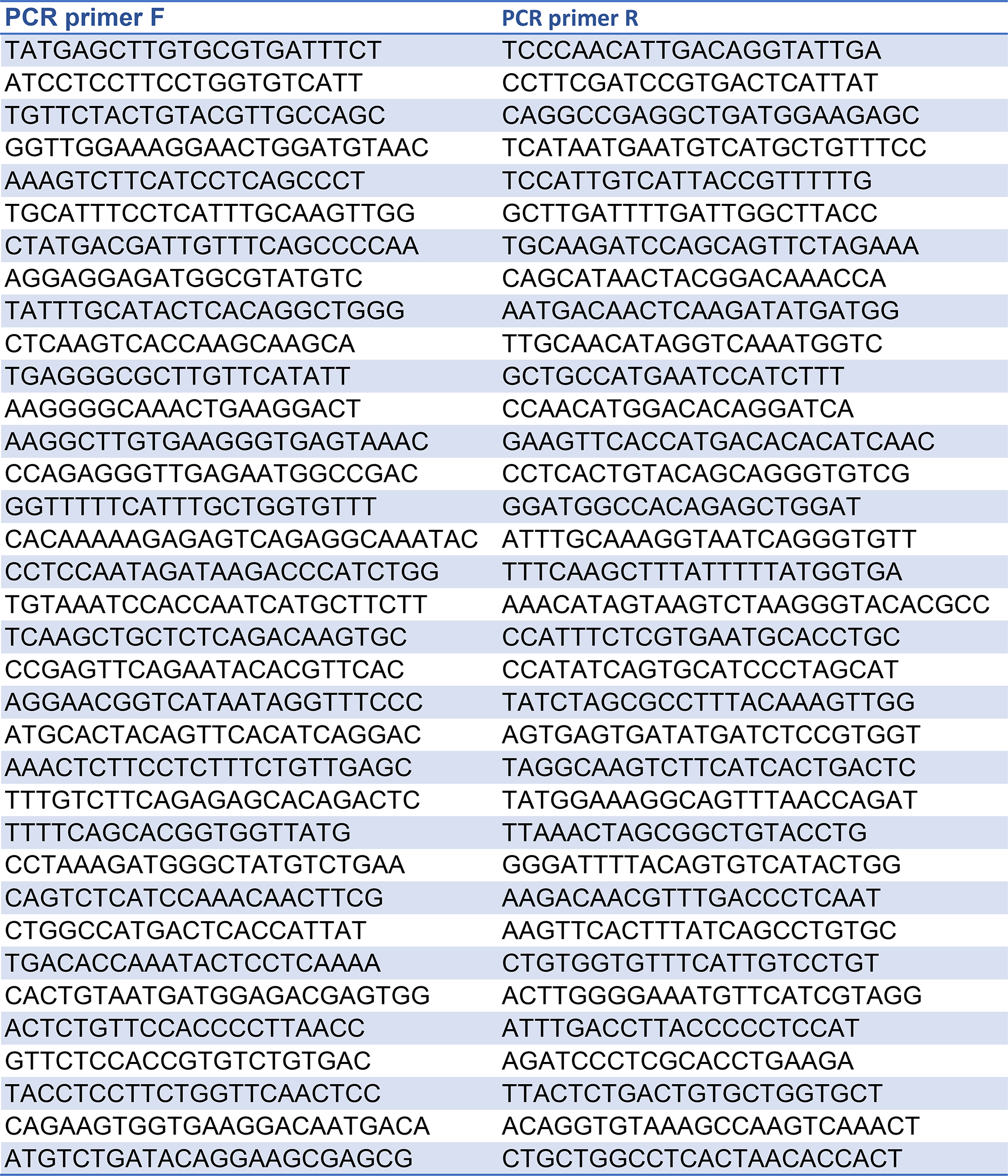

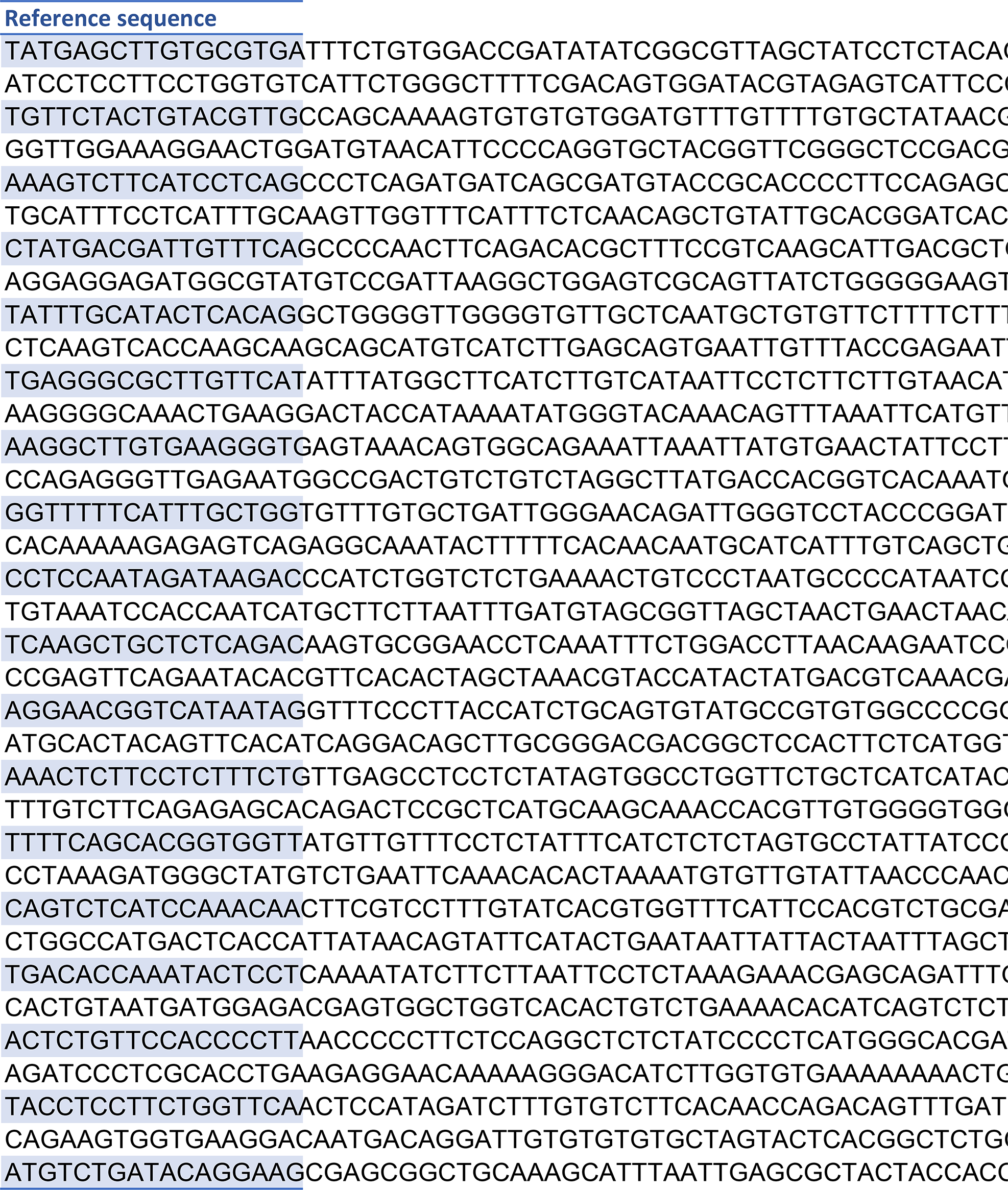

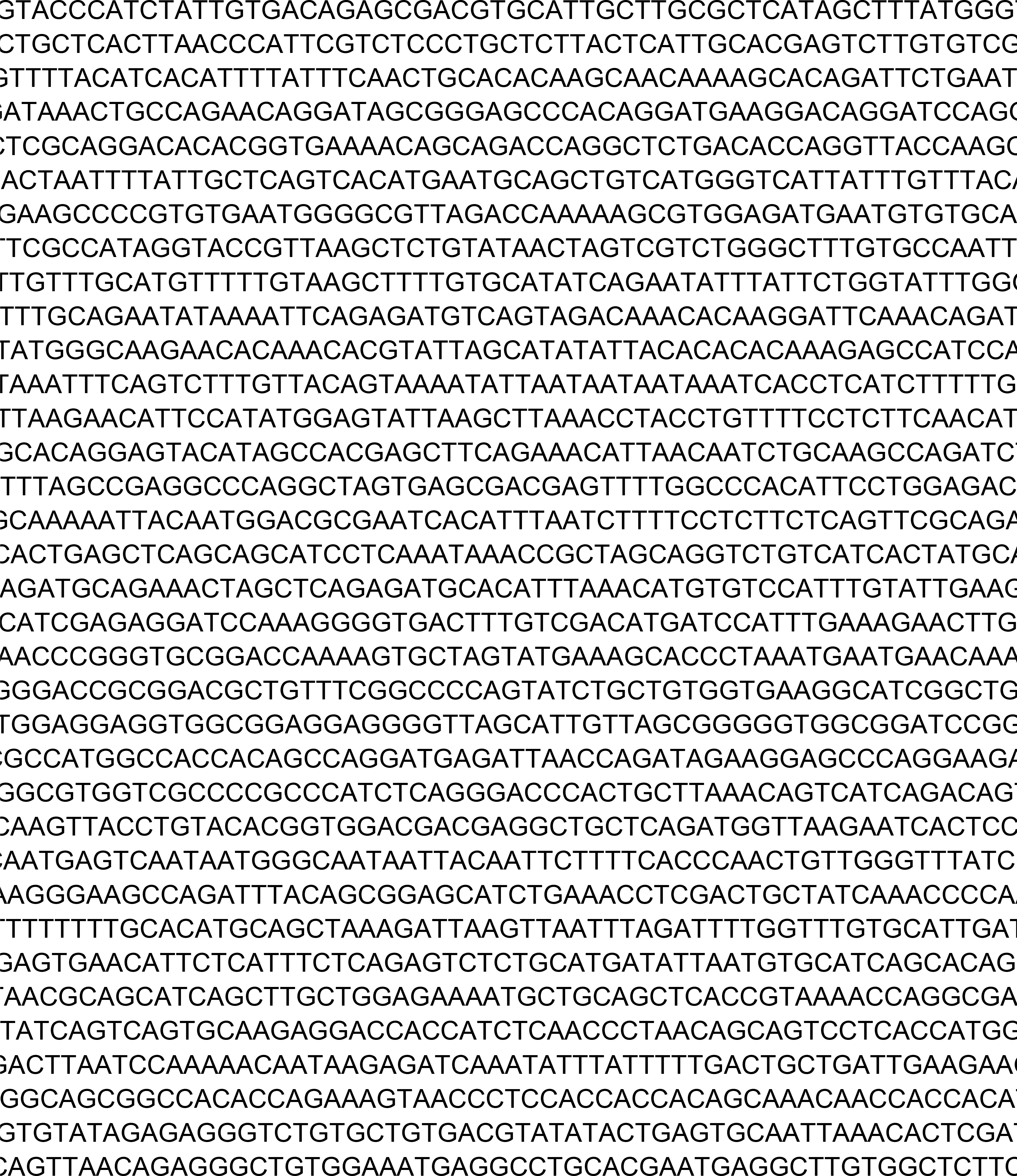

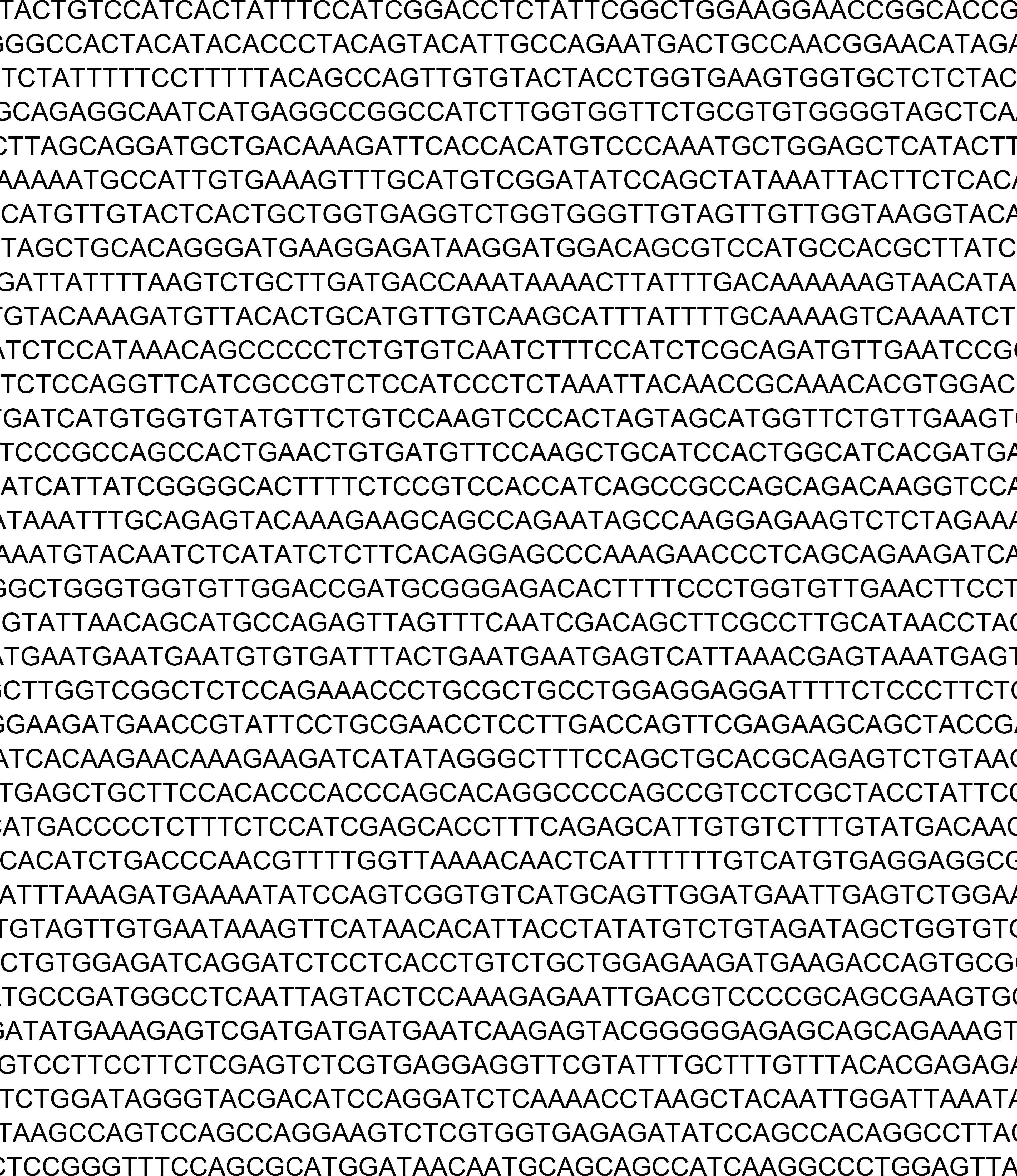

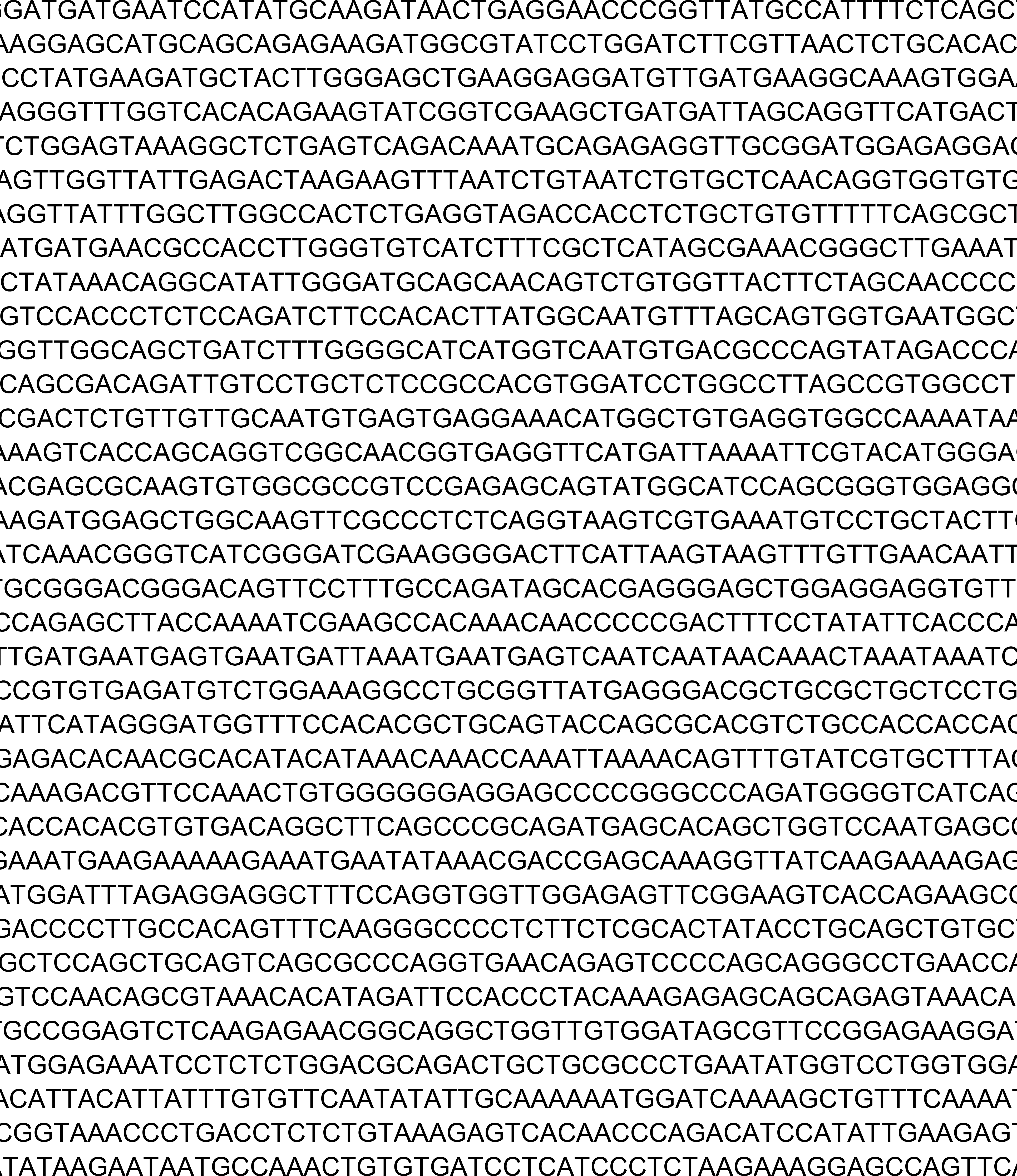

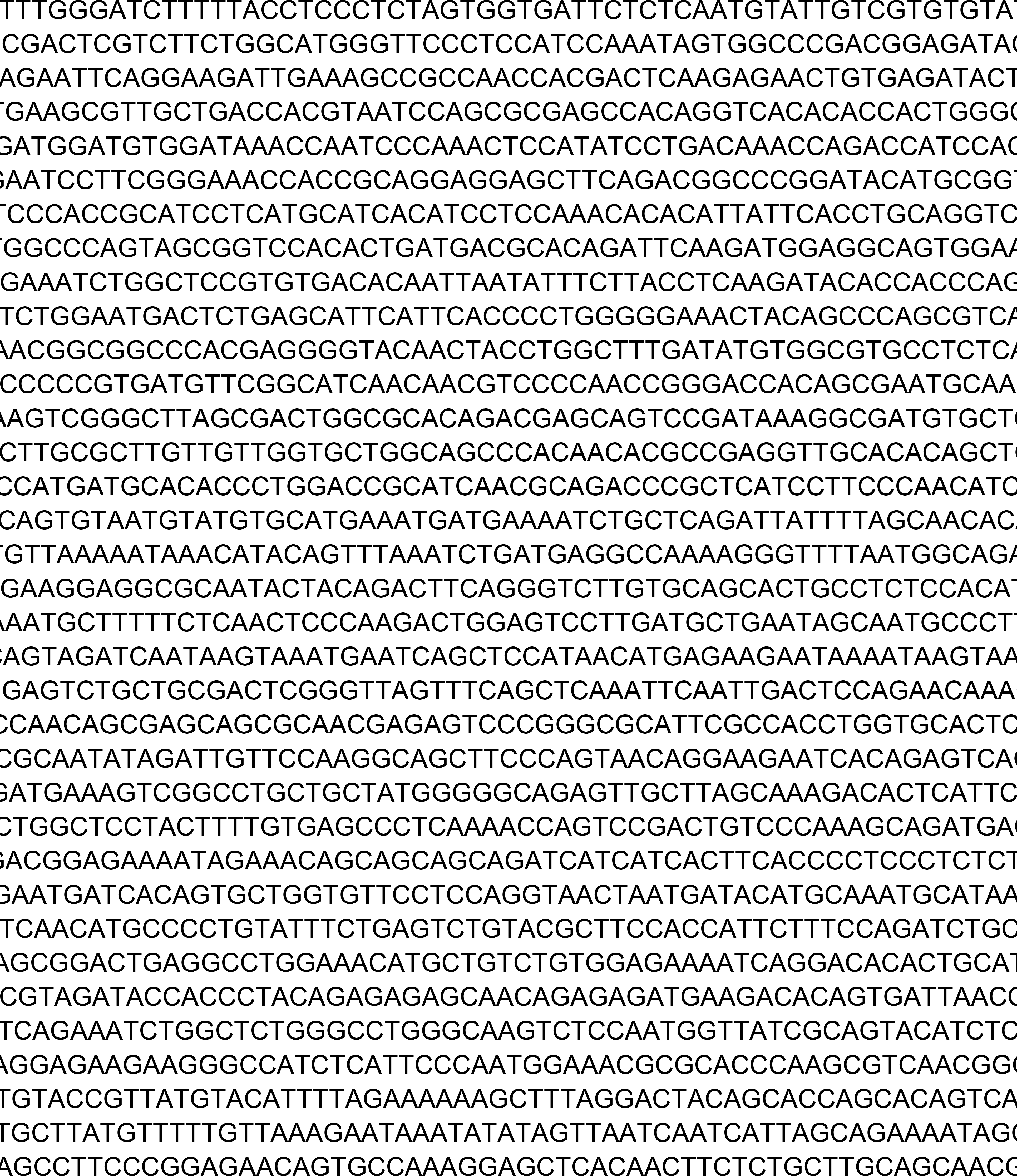

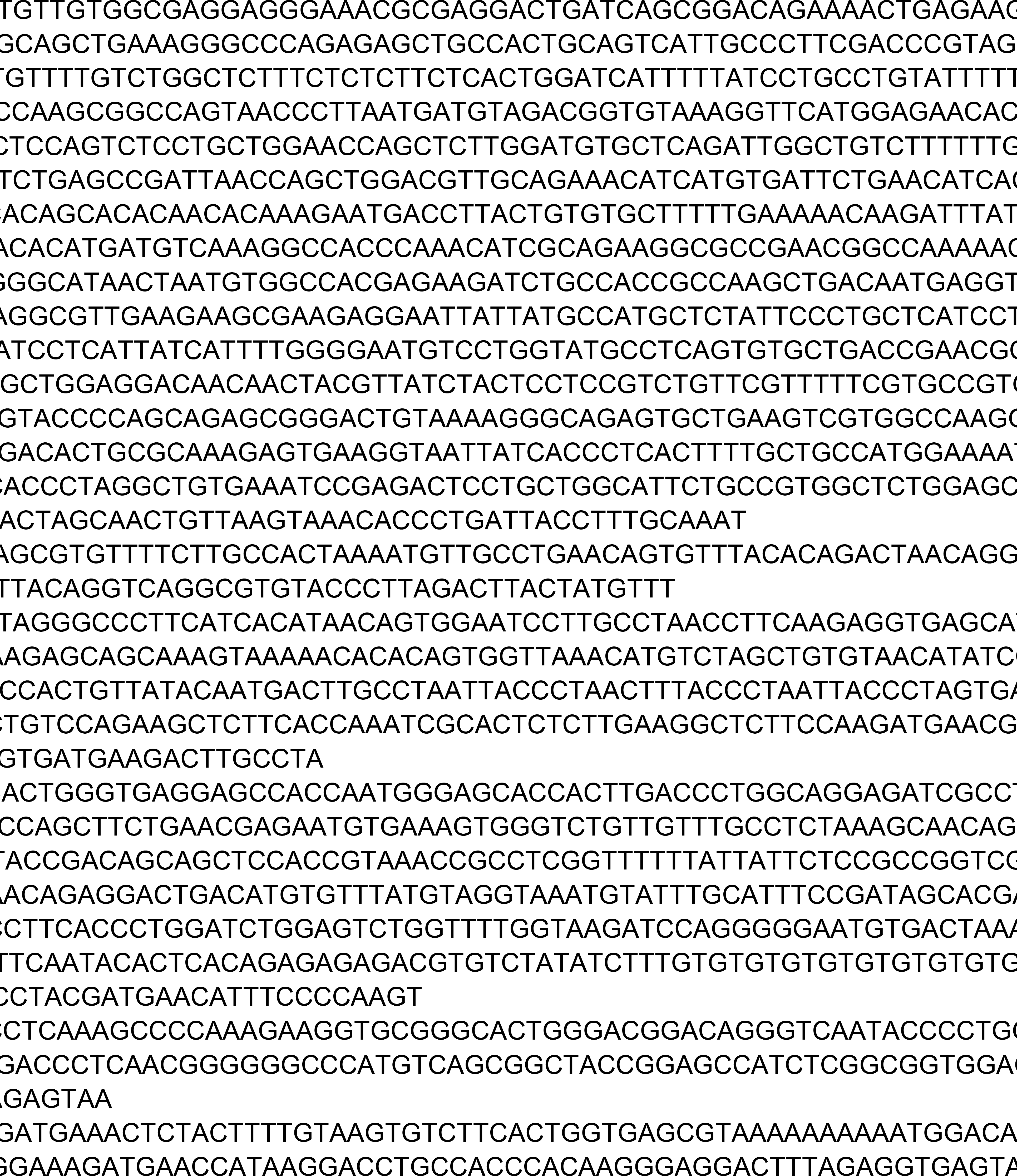

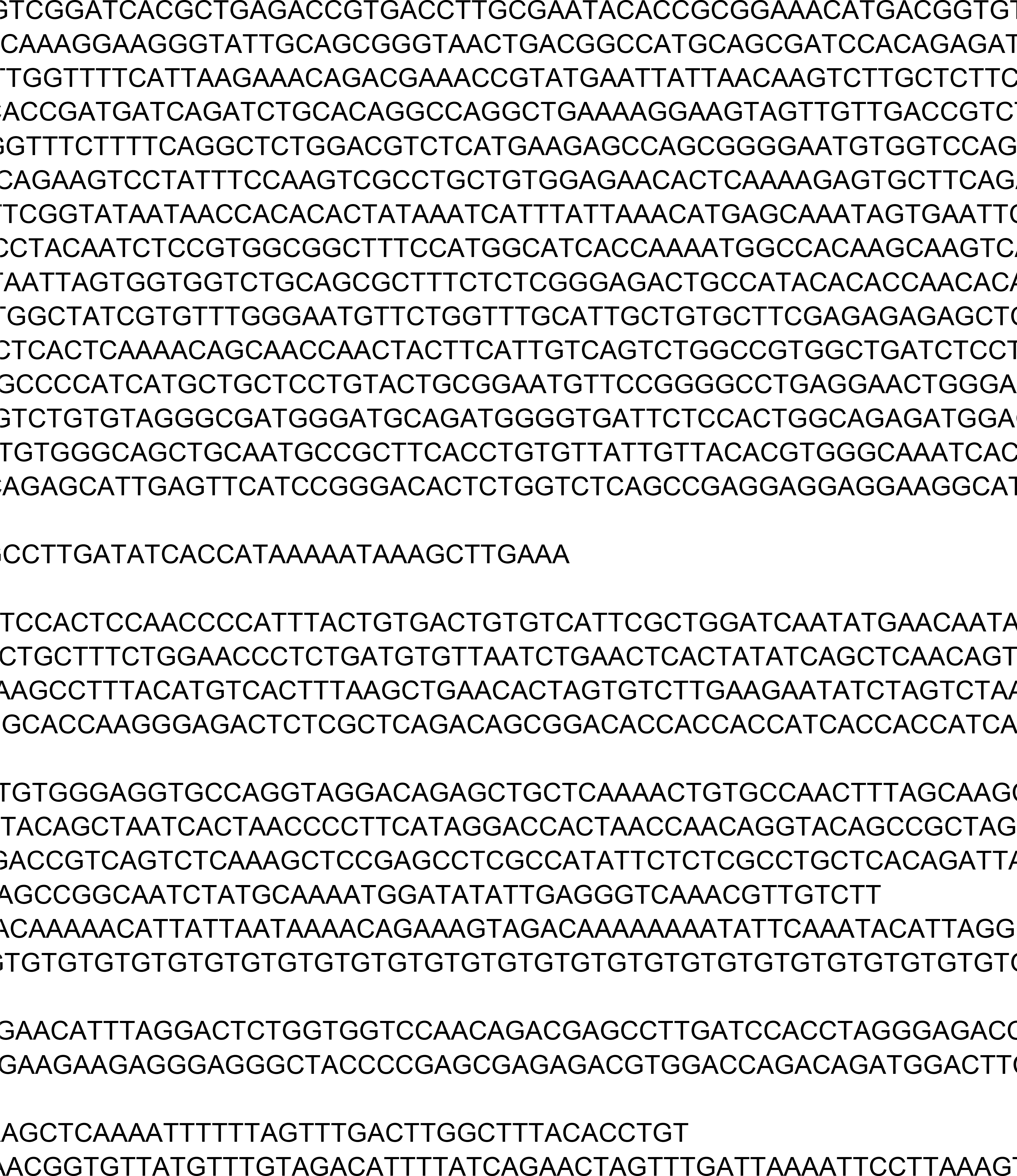

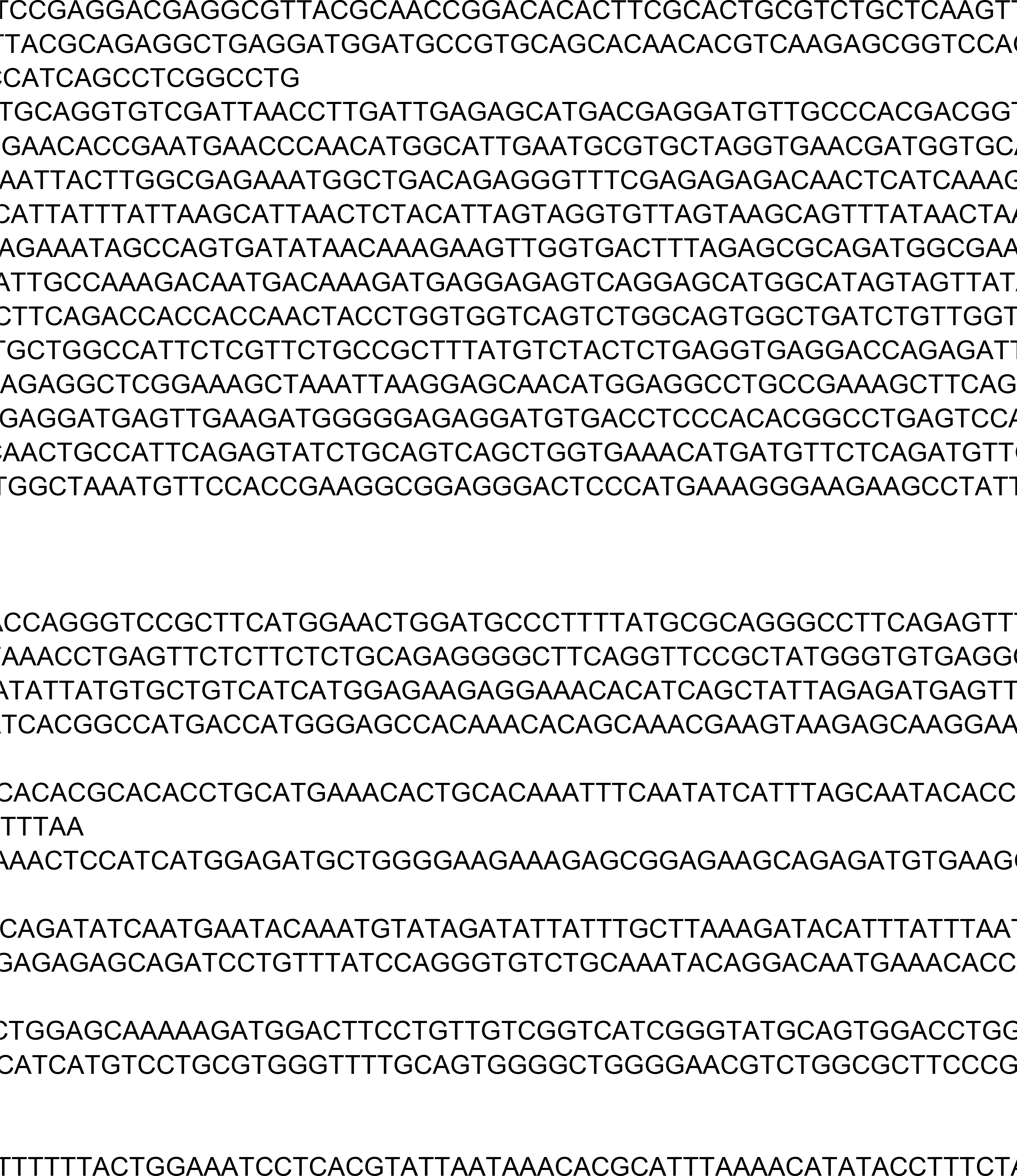

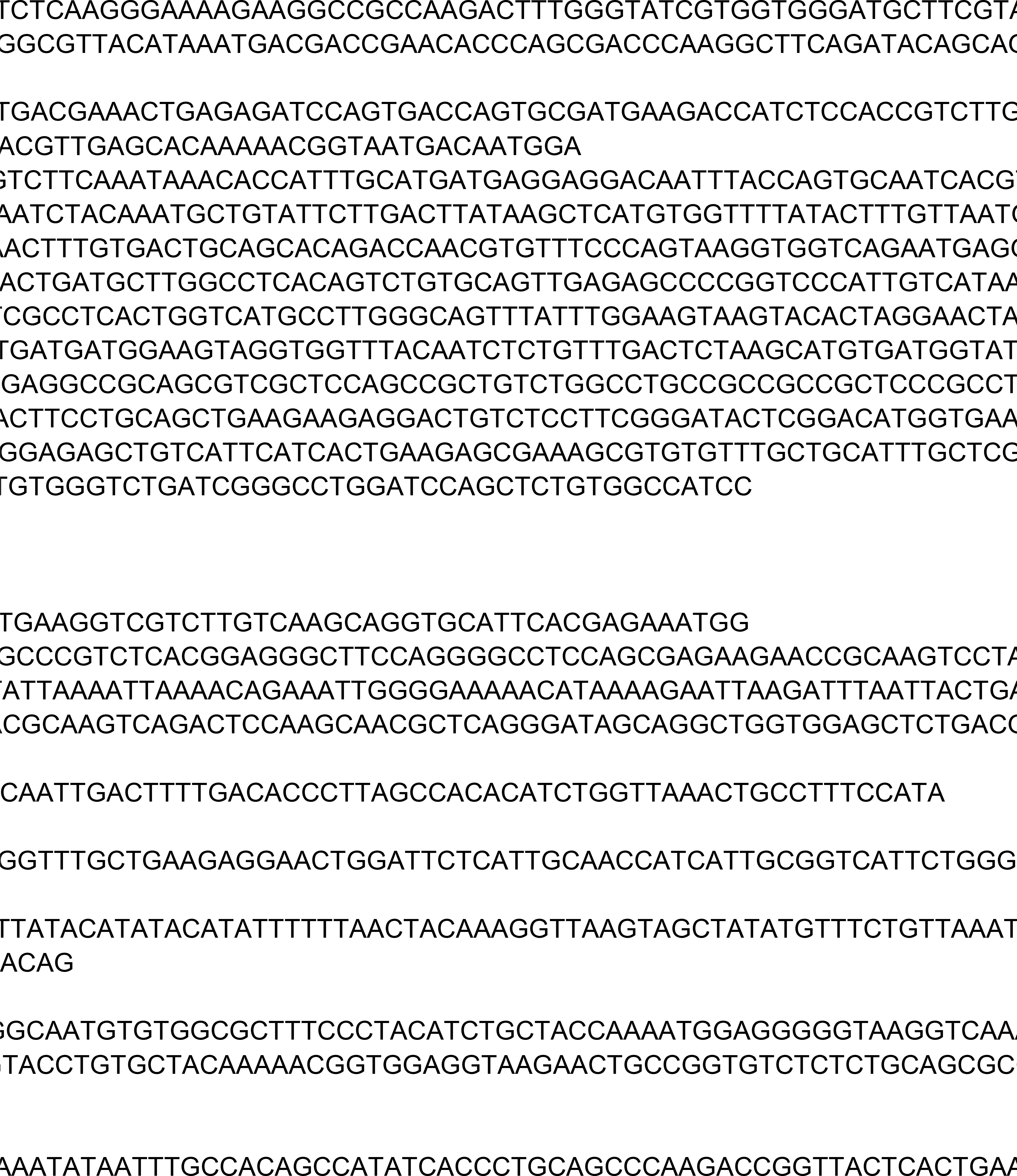

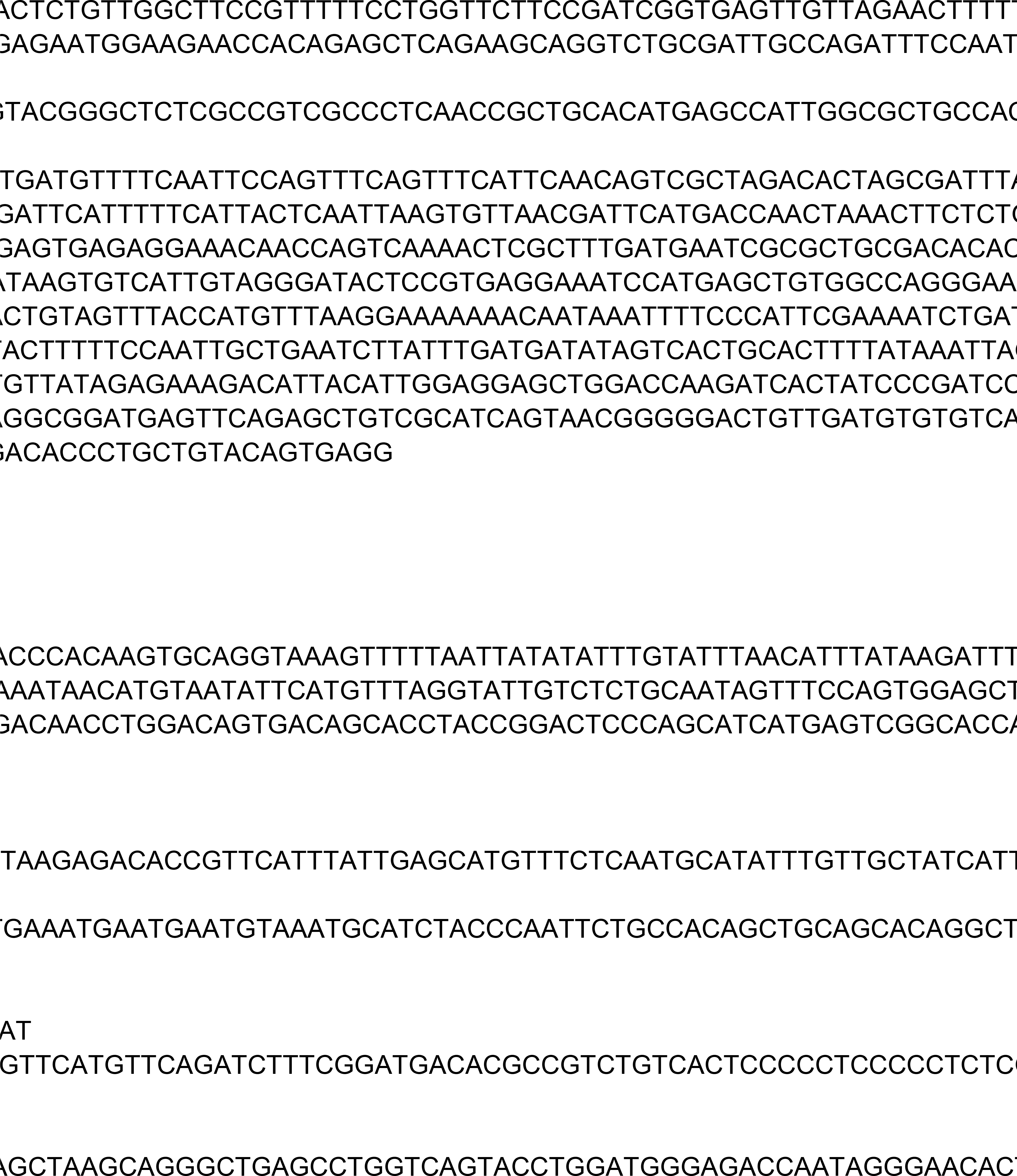

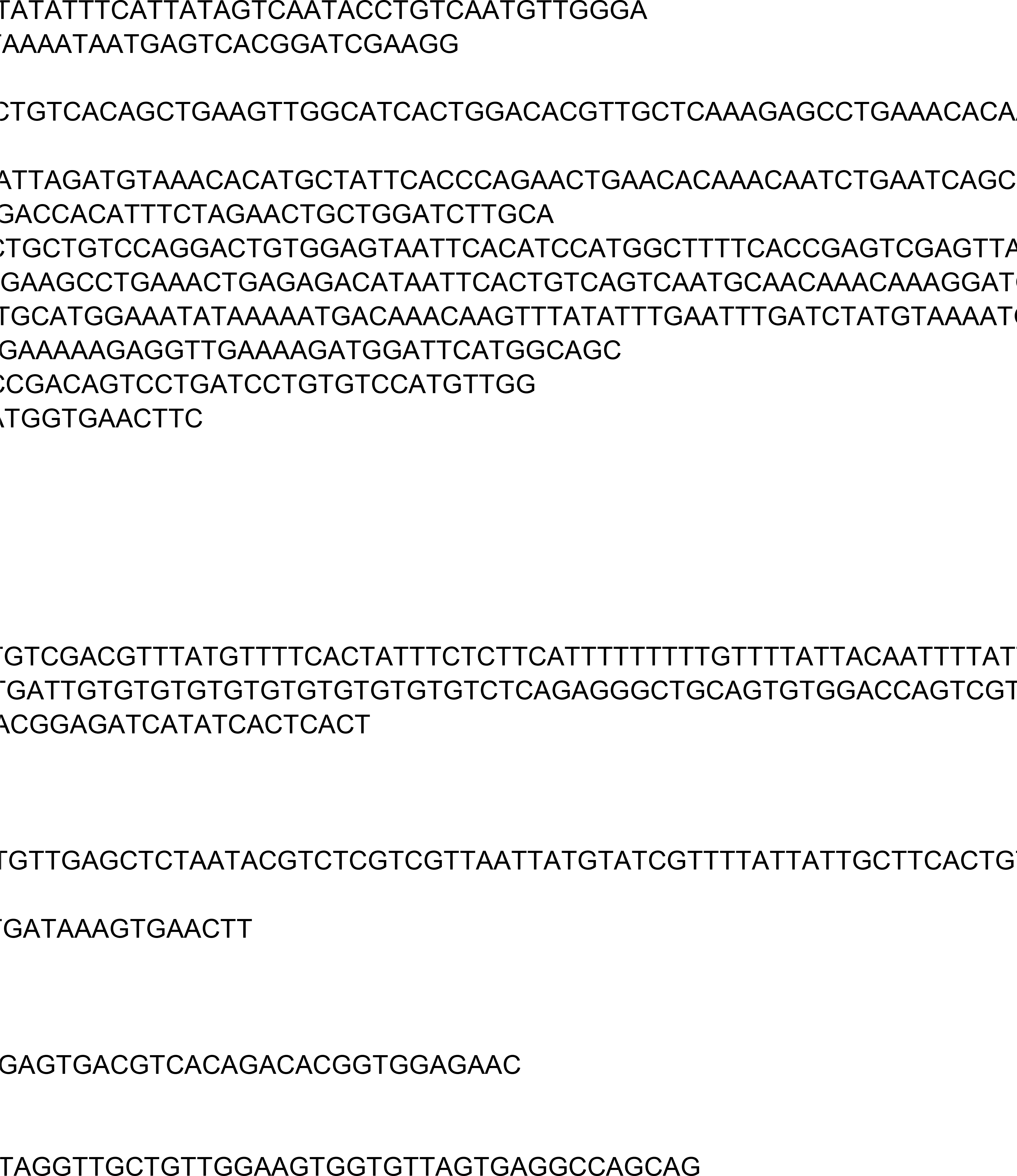

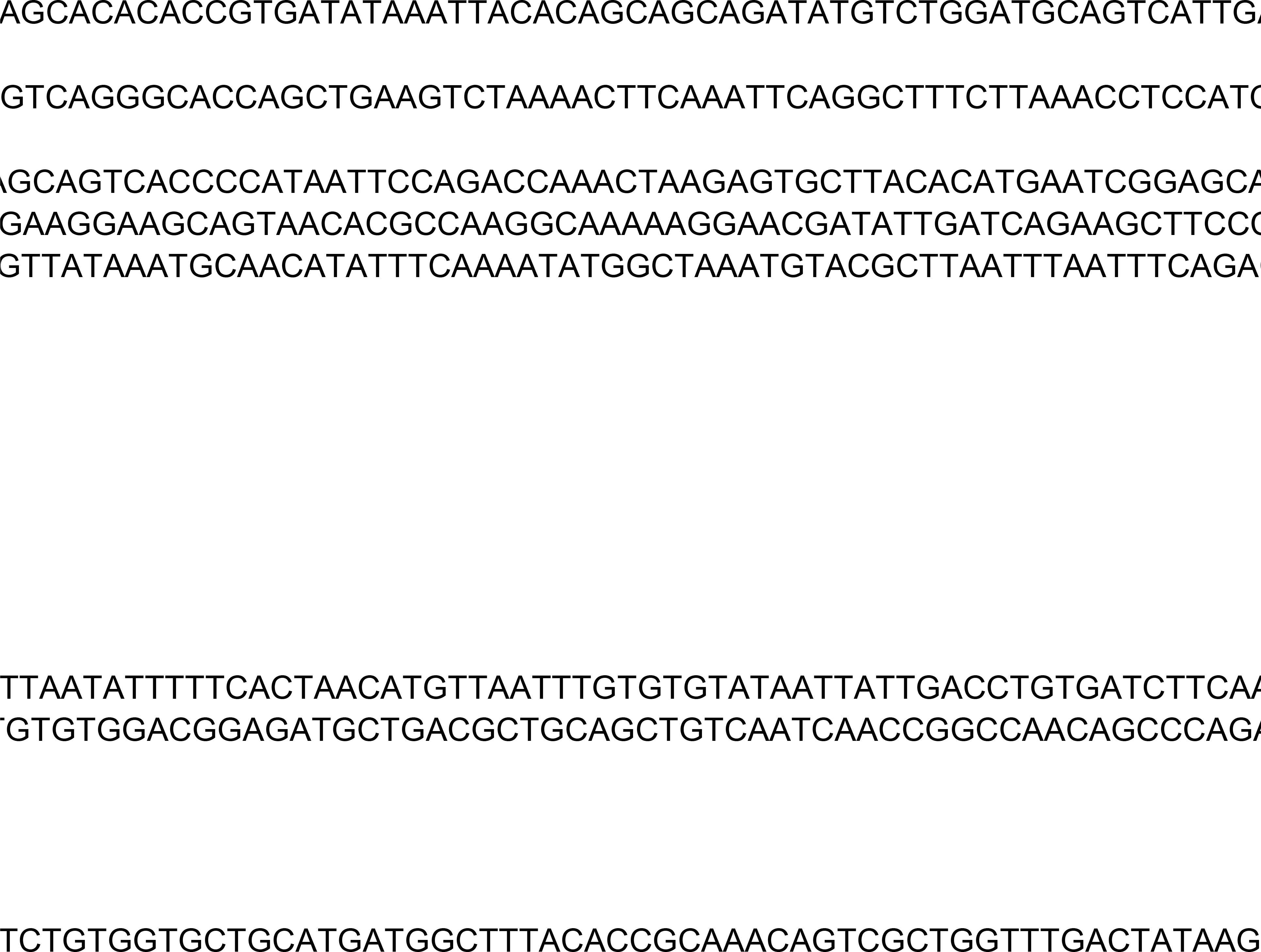

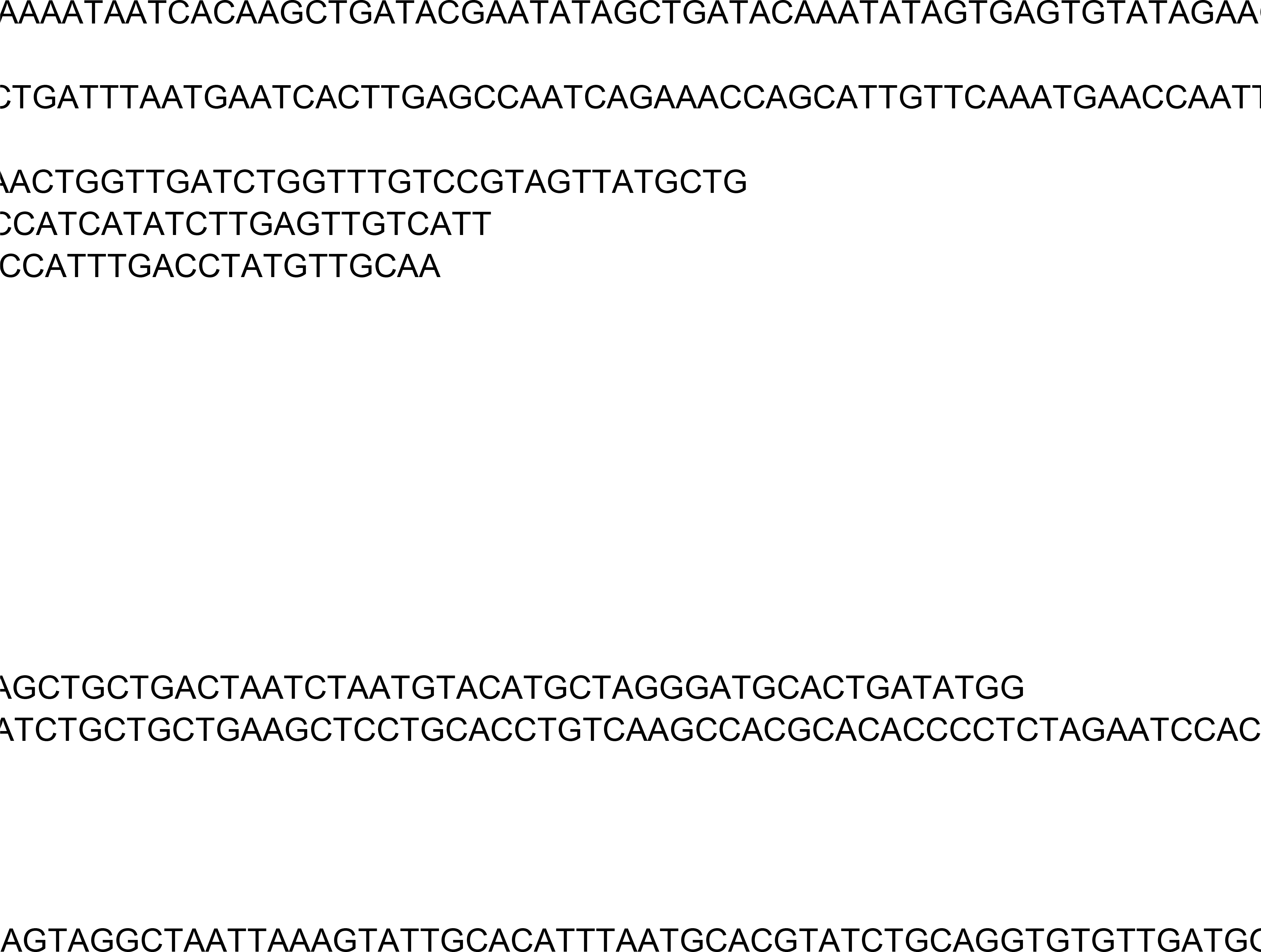

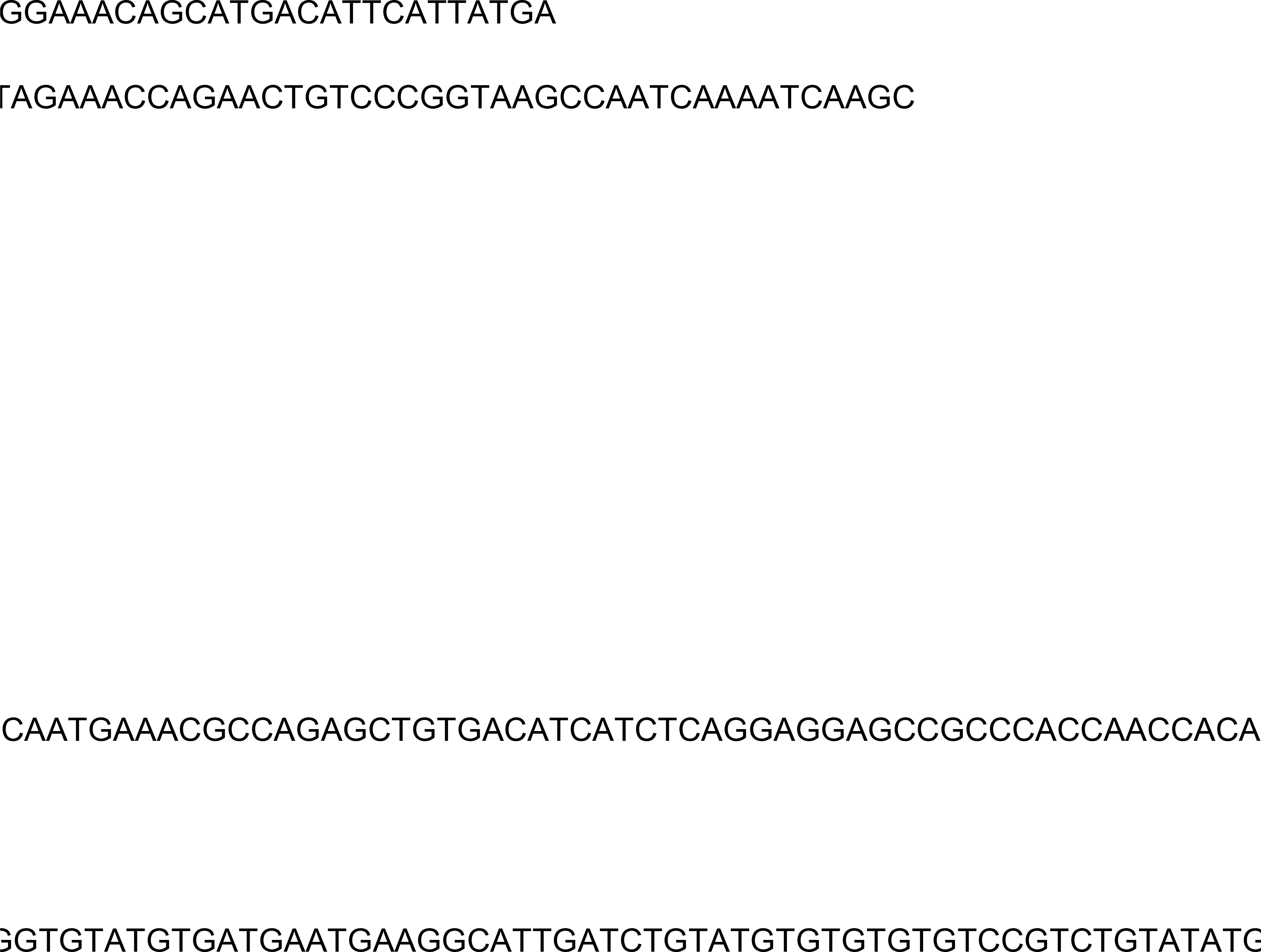

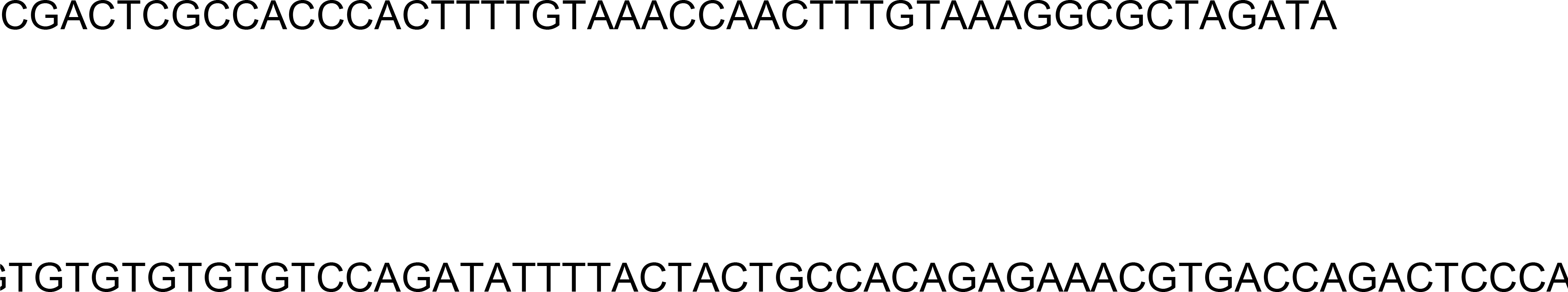

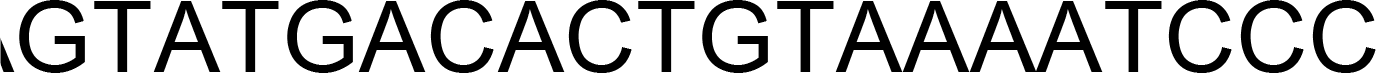

**Supplemental File 1.**
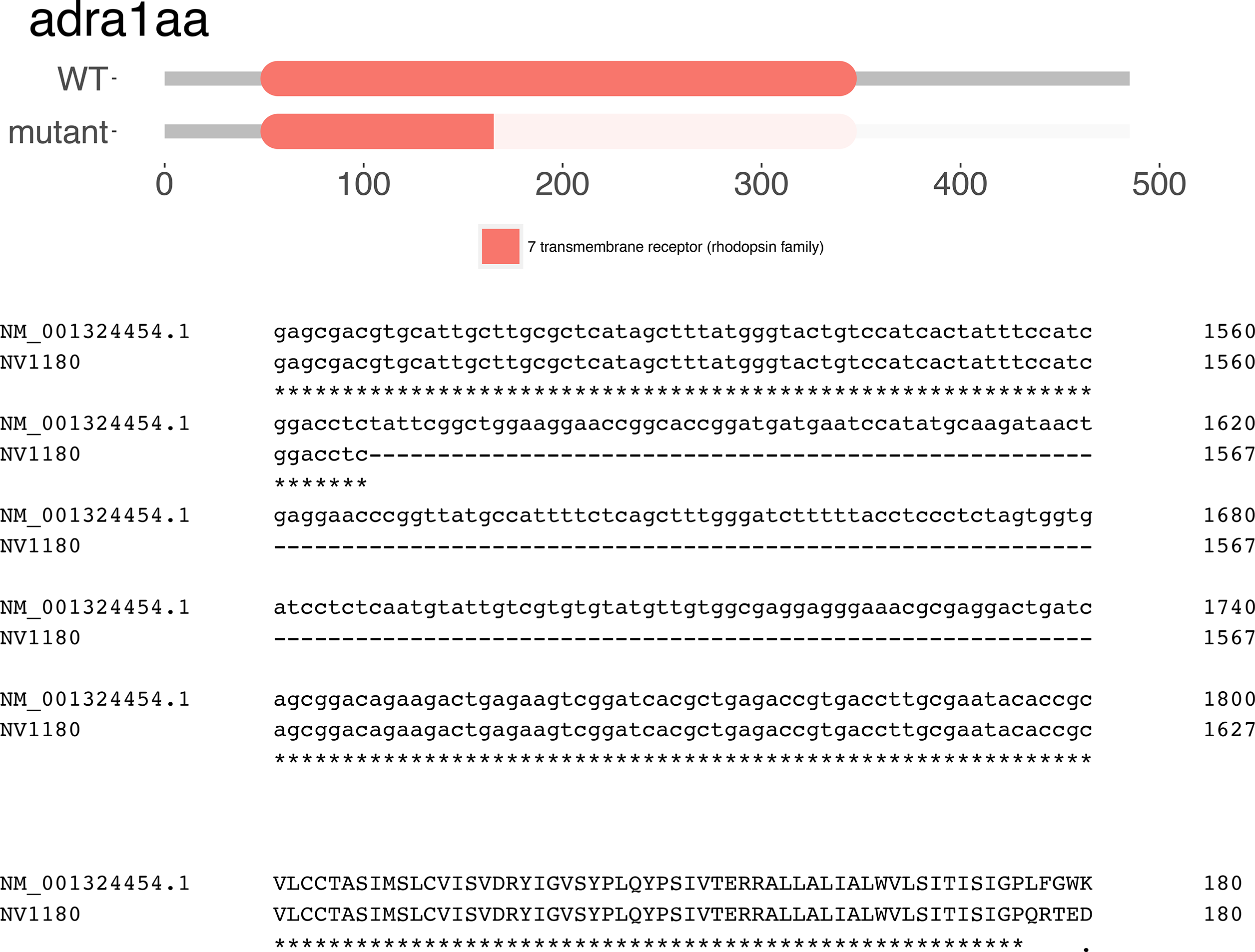

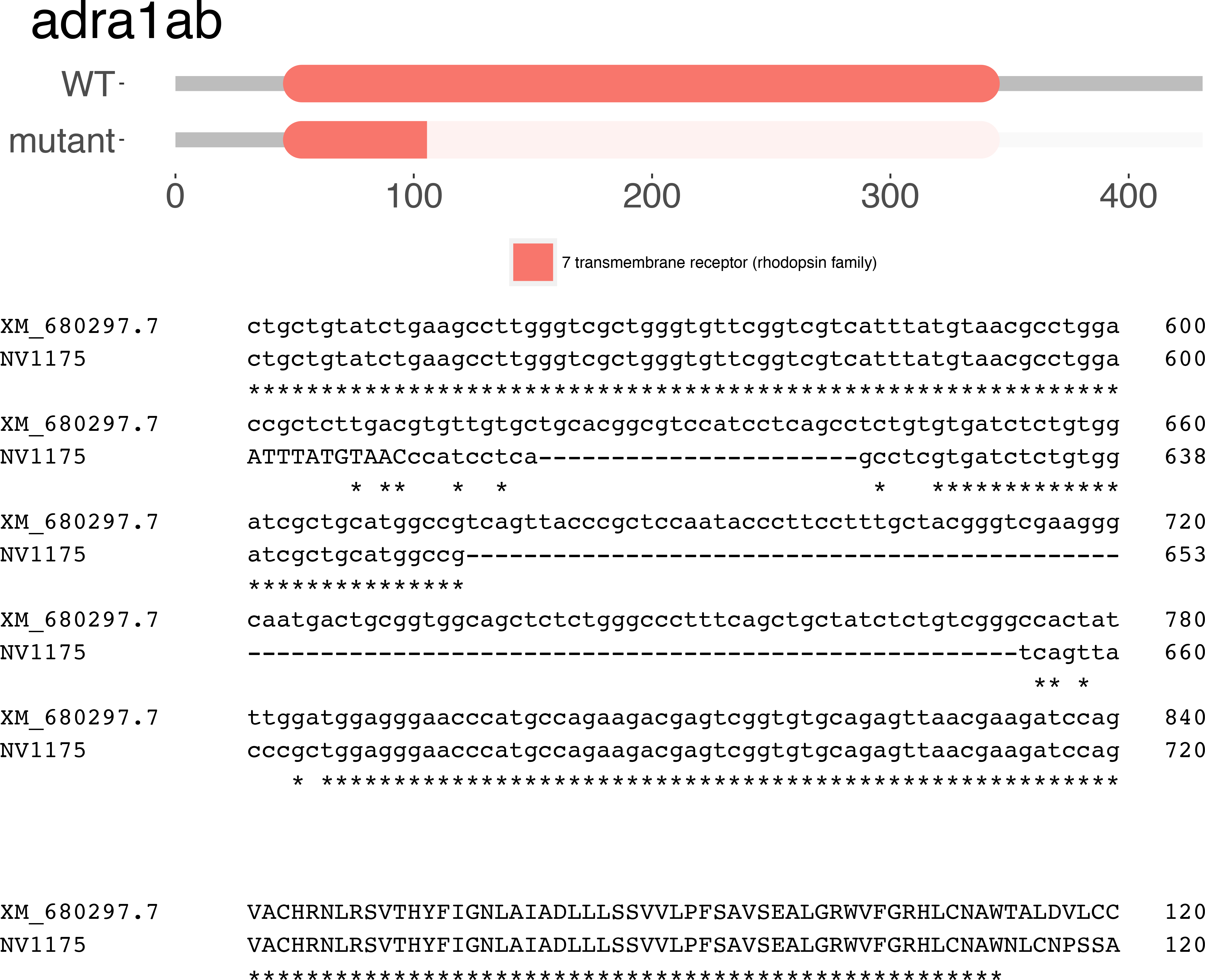

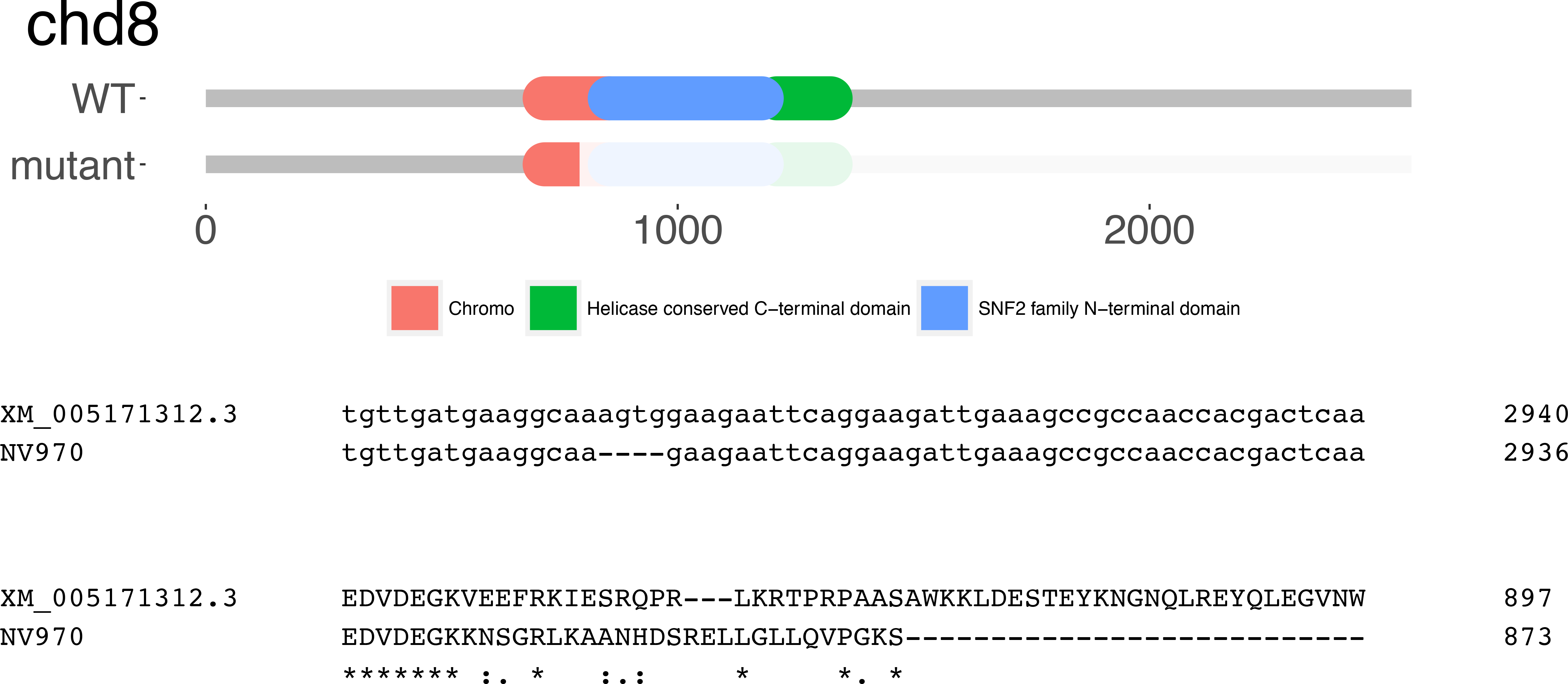

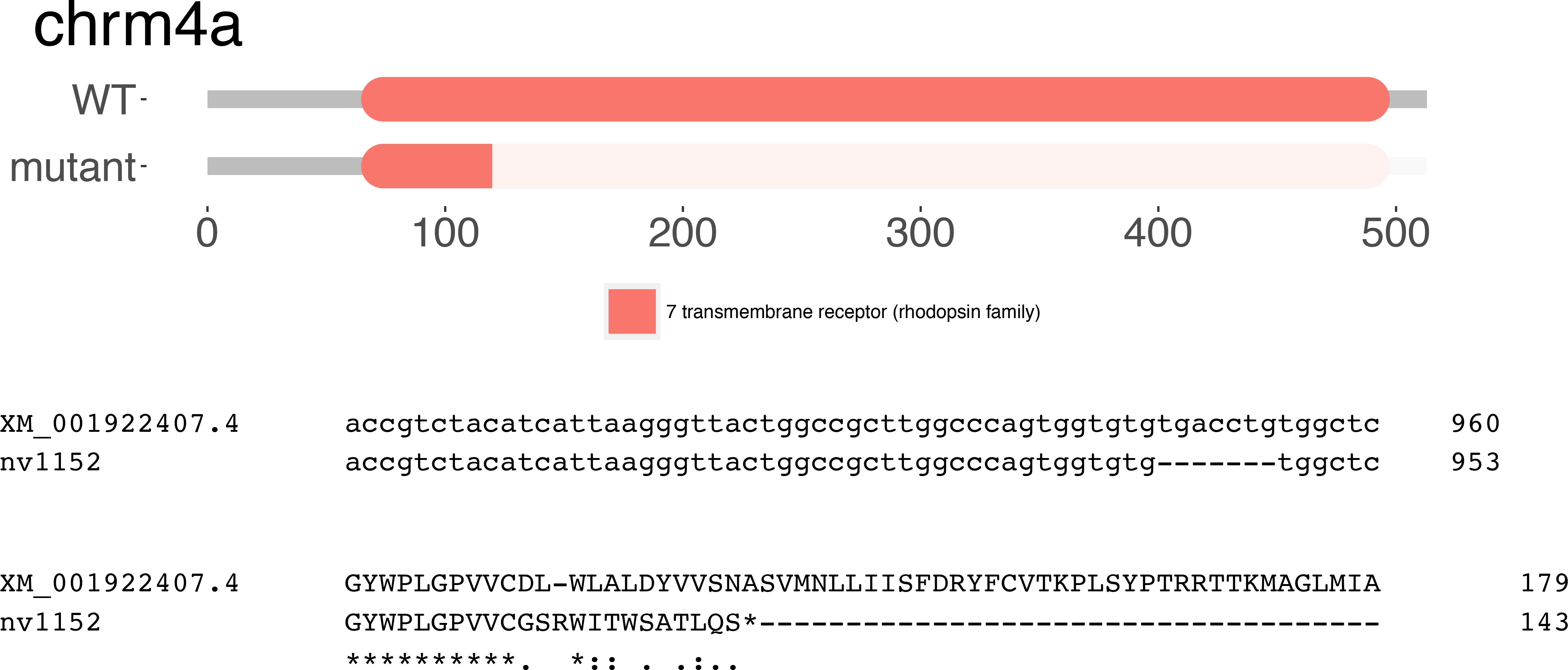

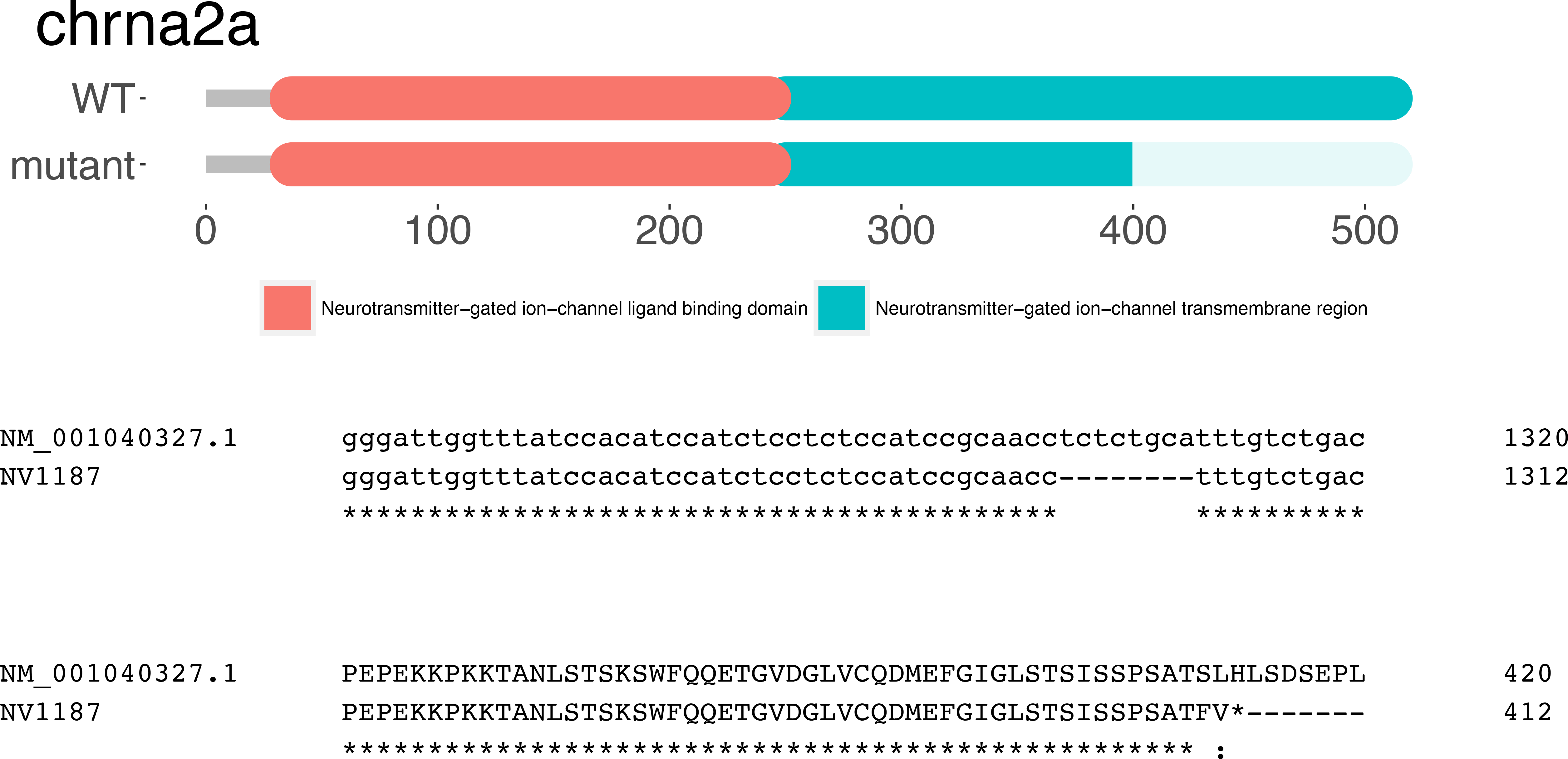

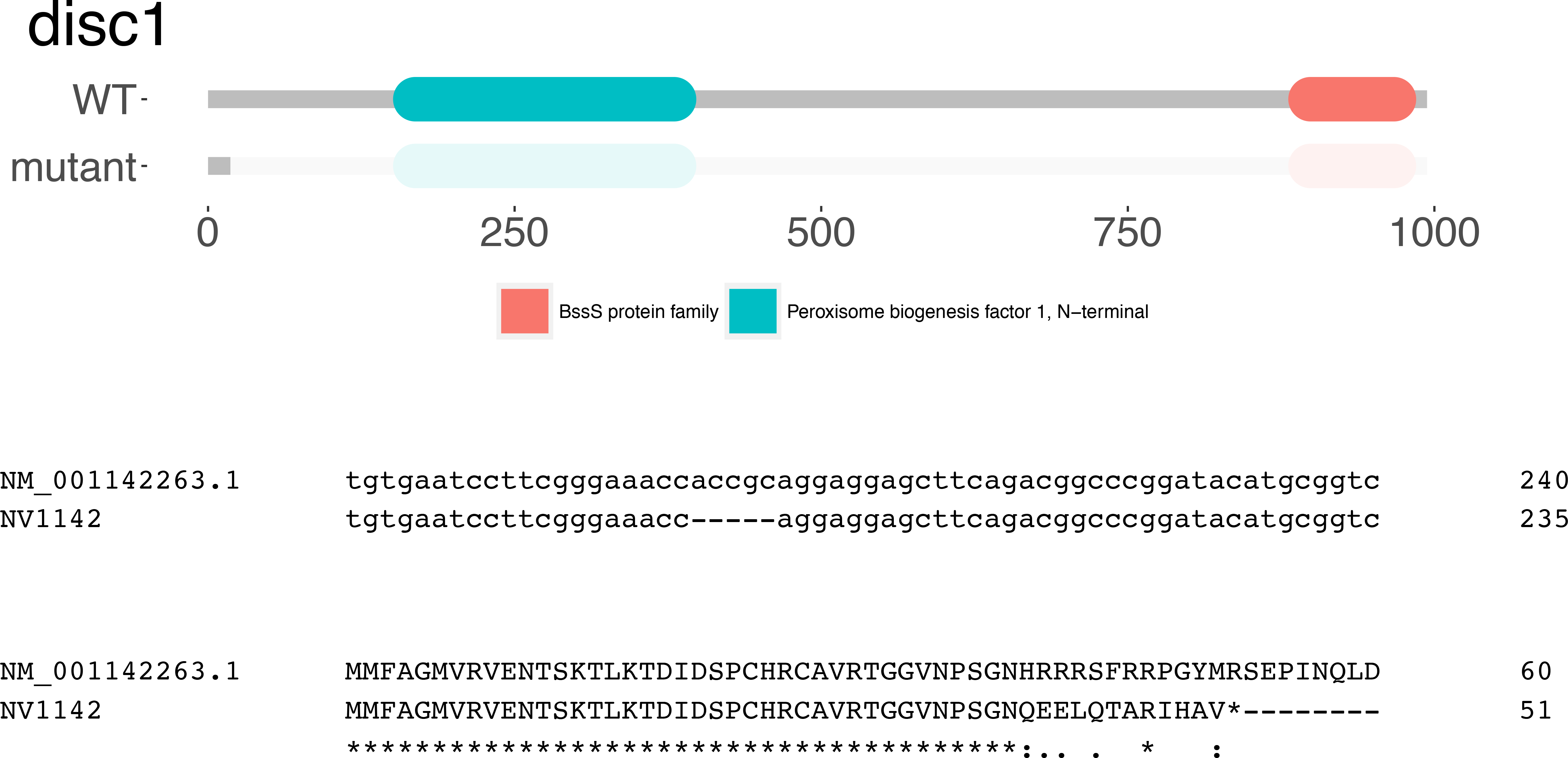

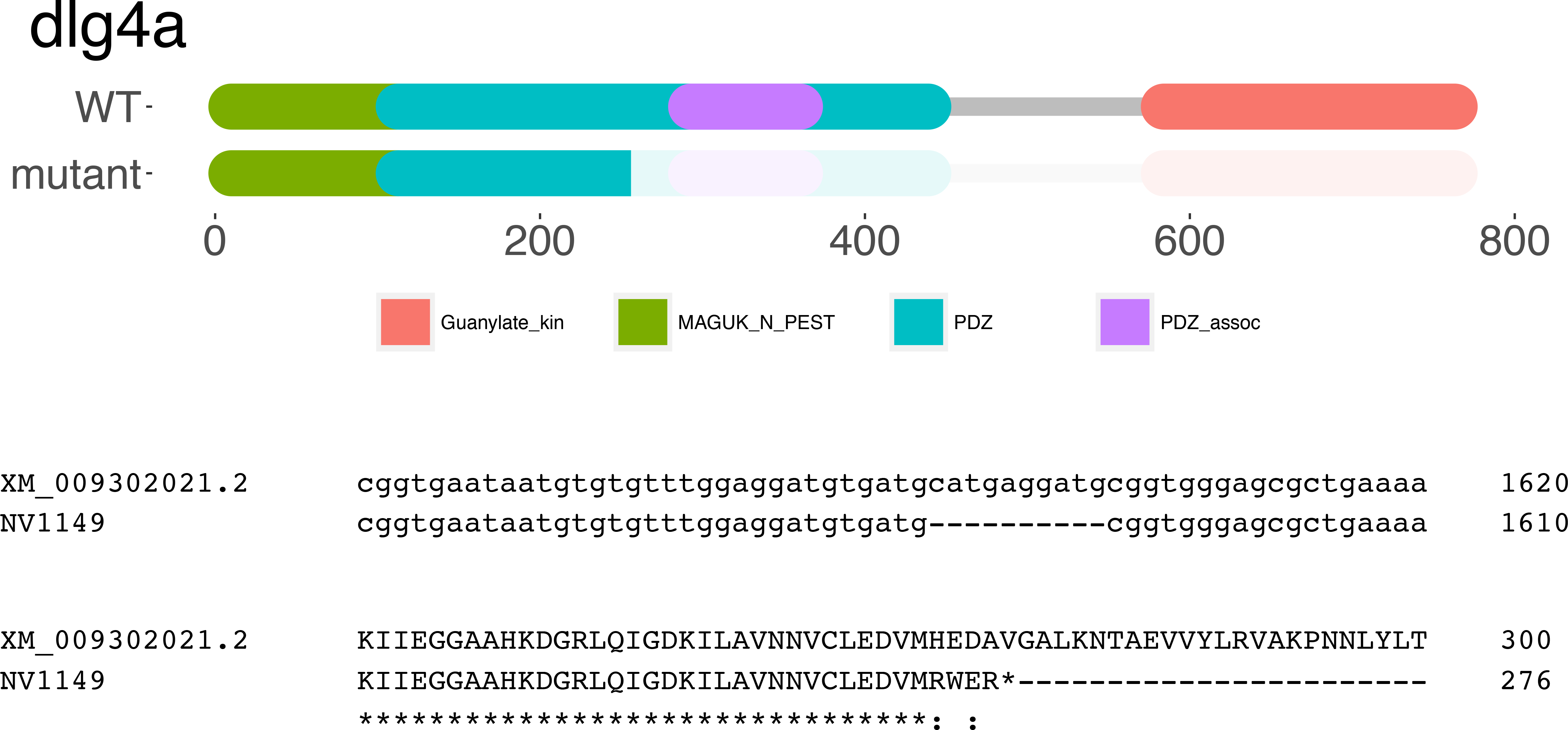

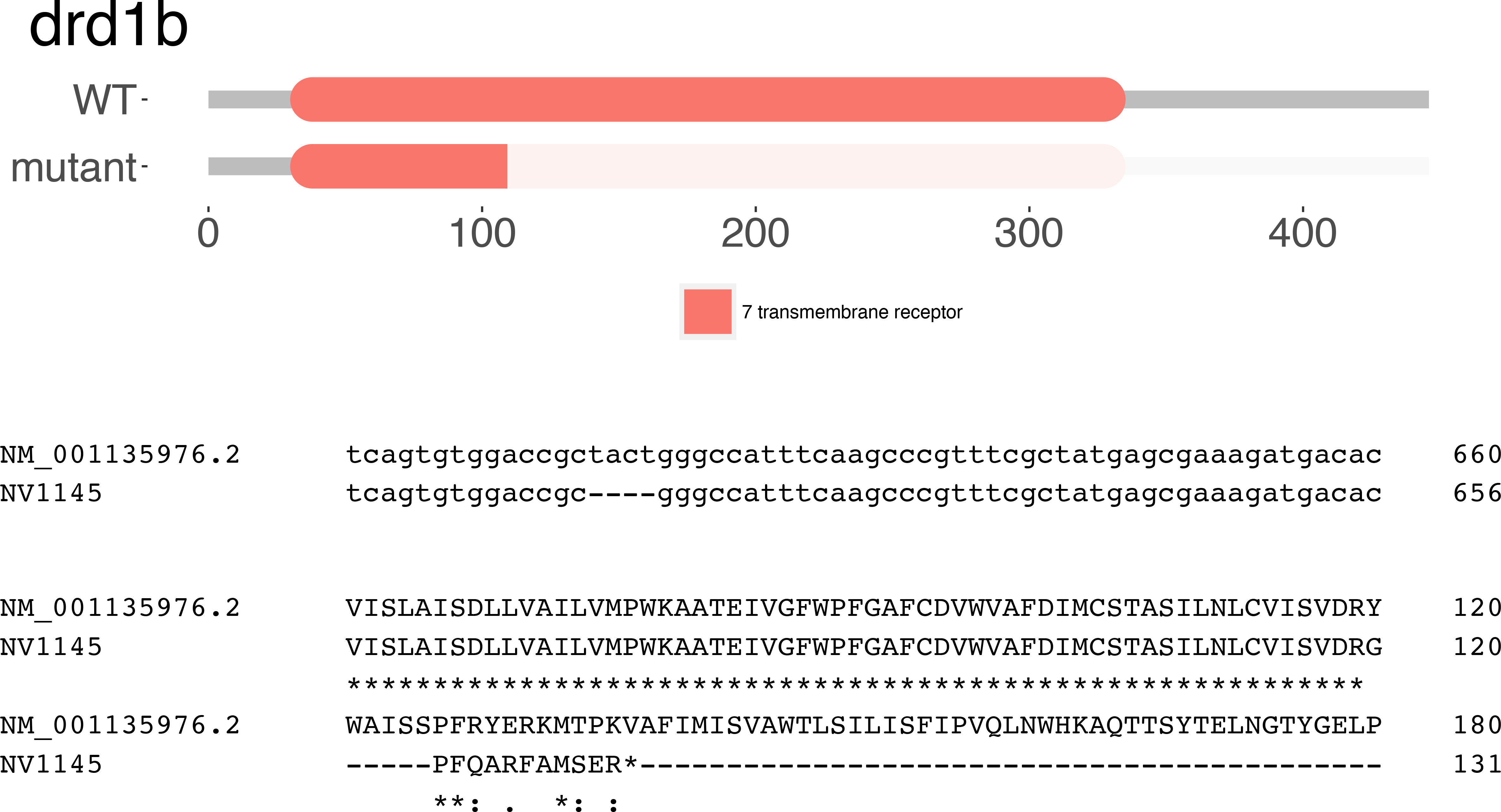

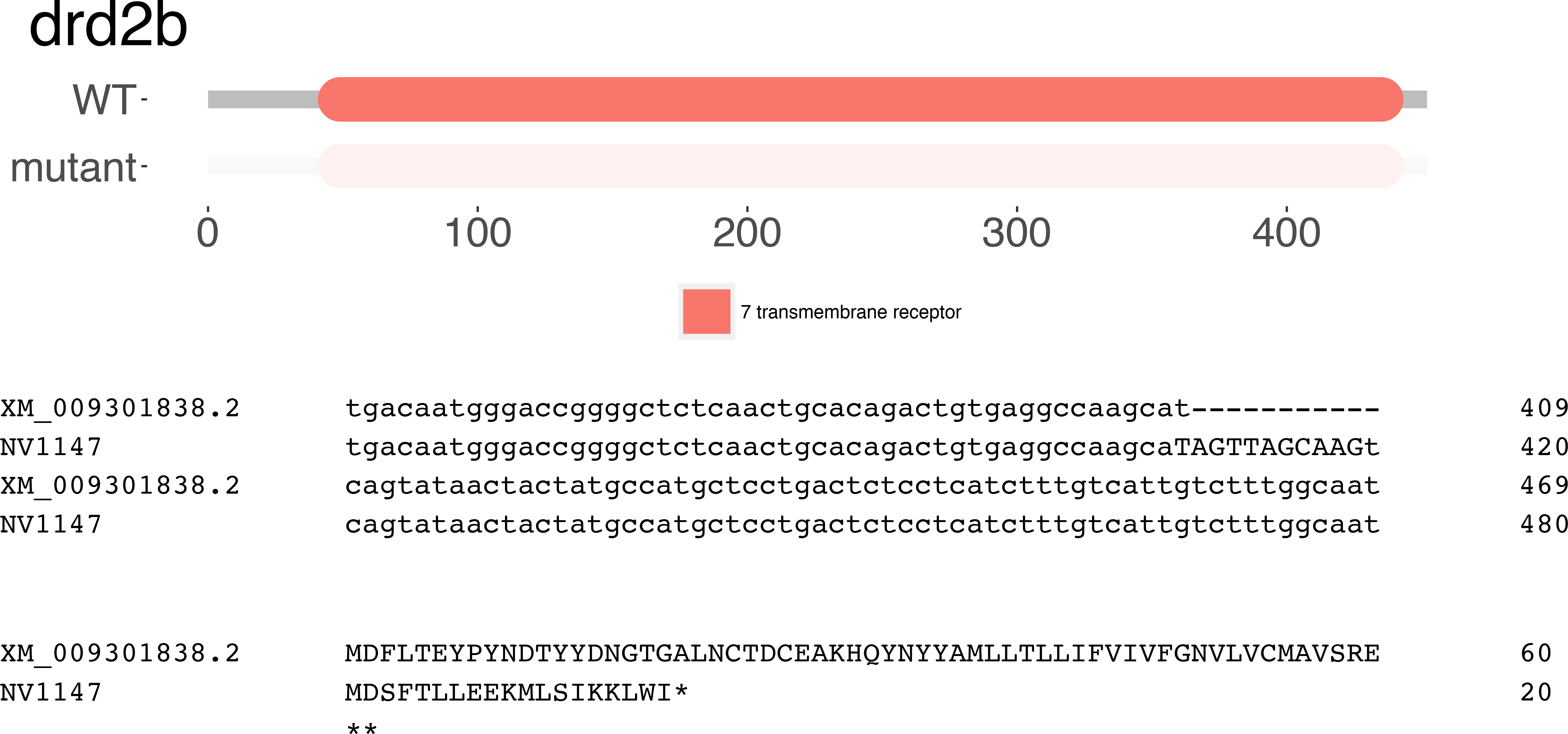

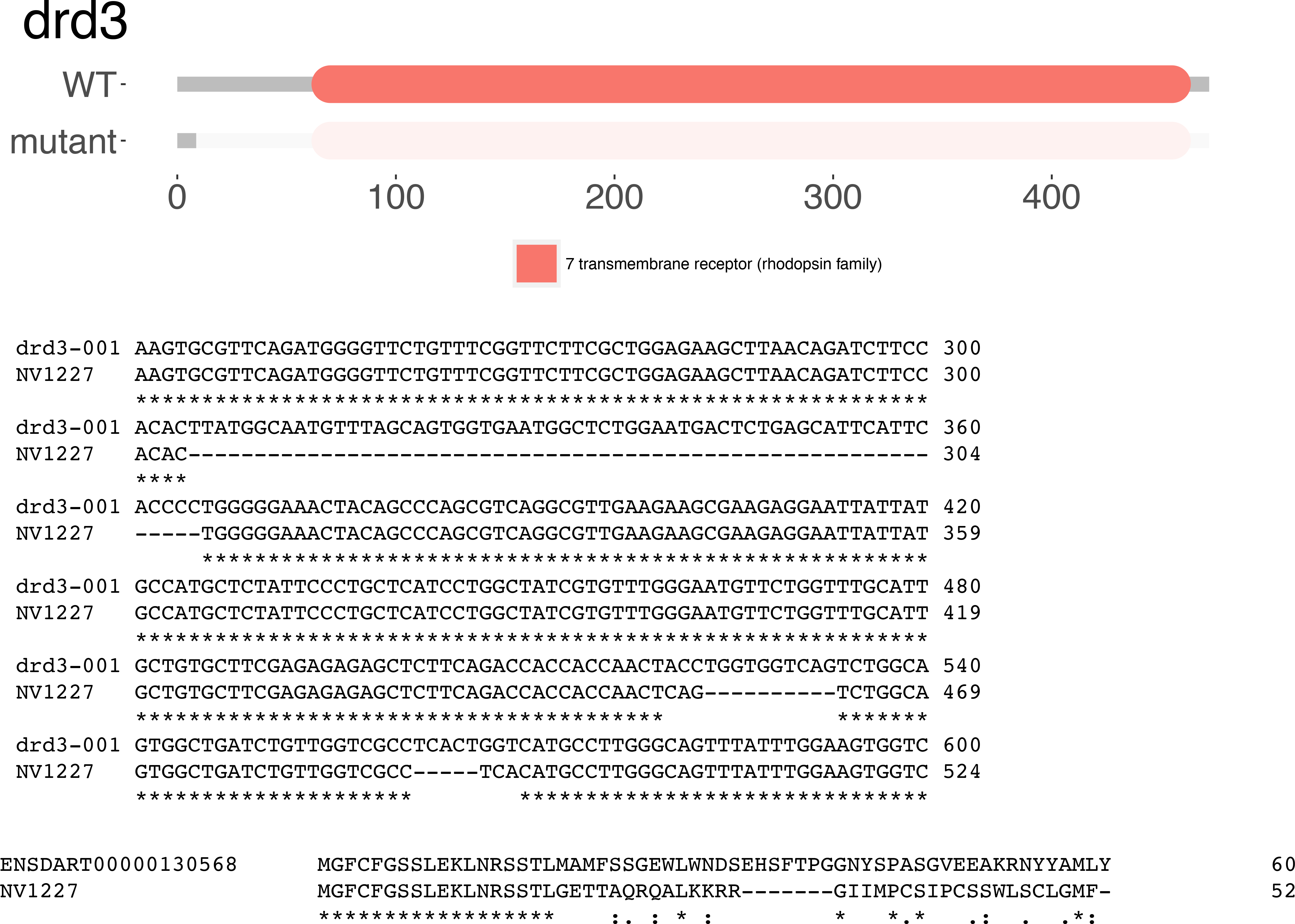

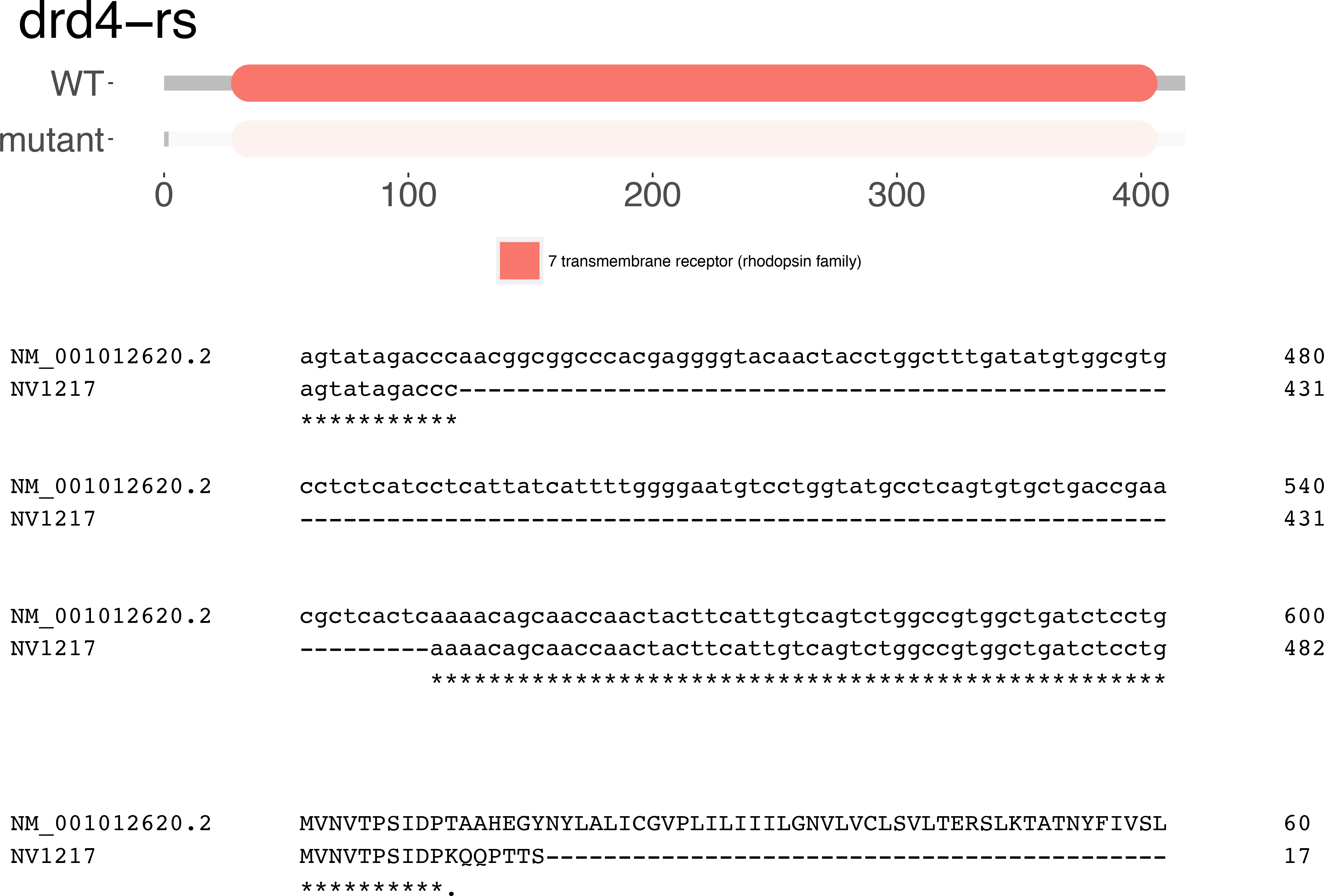

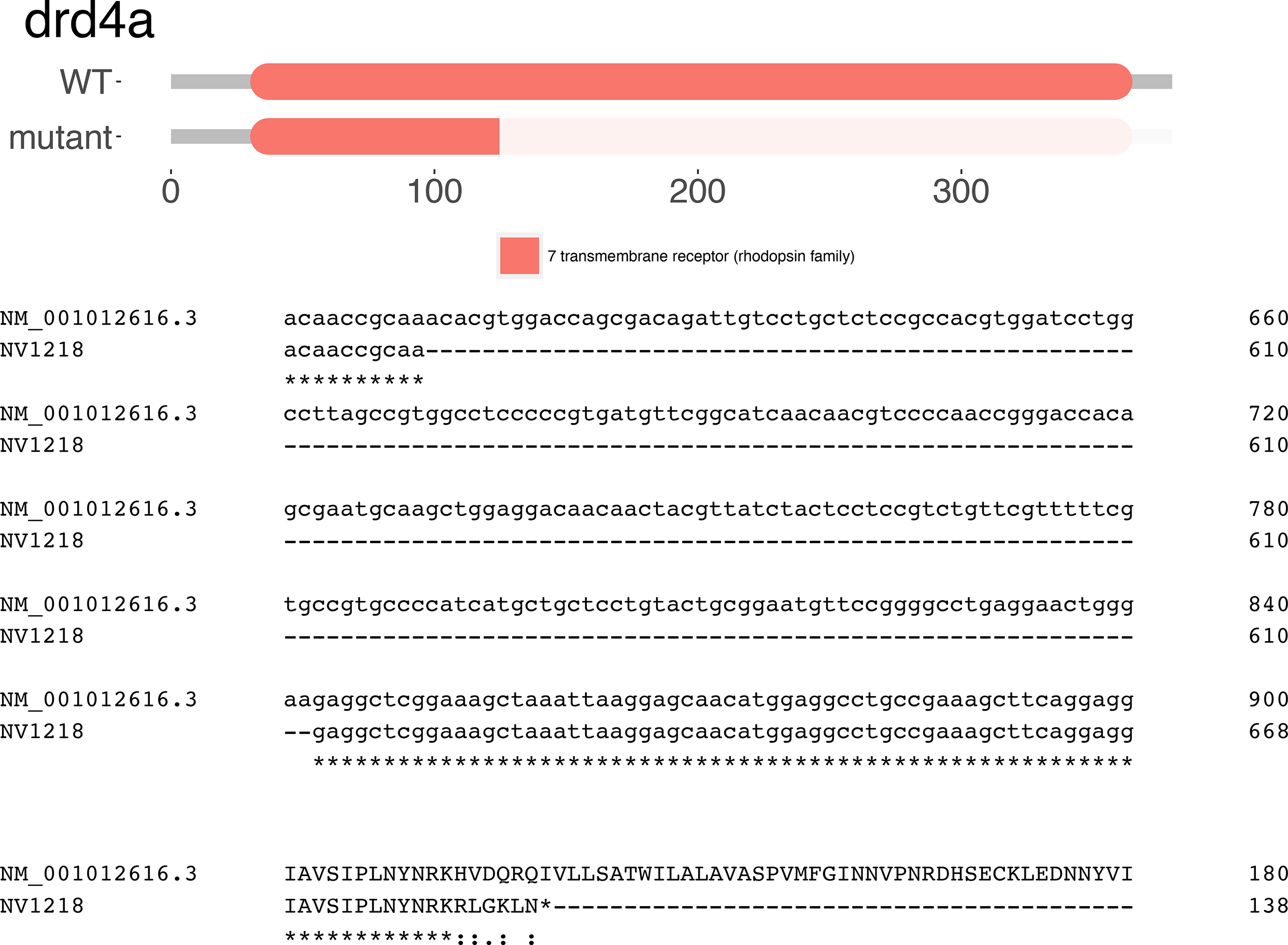

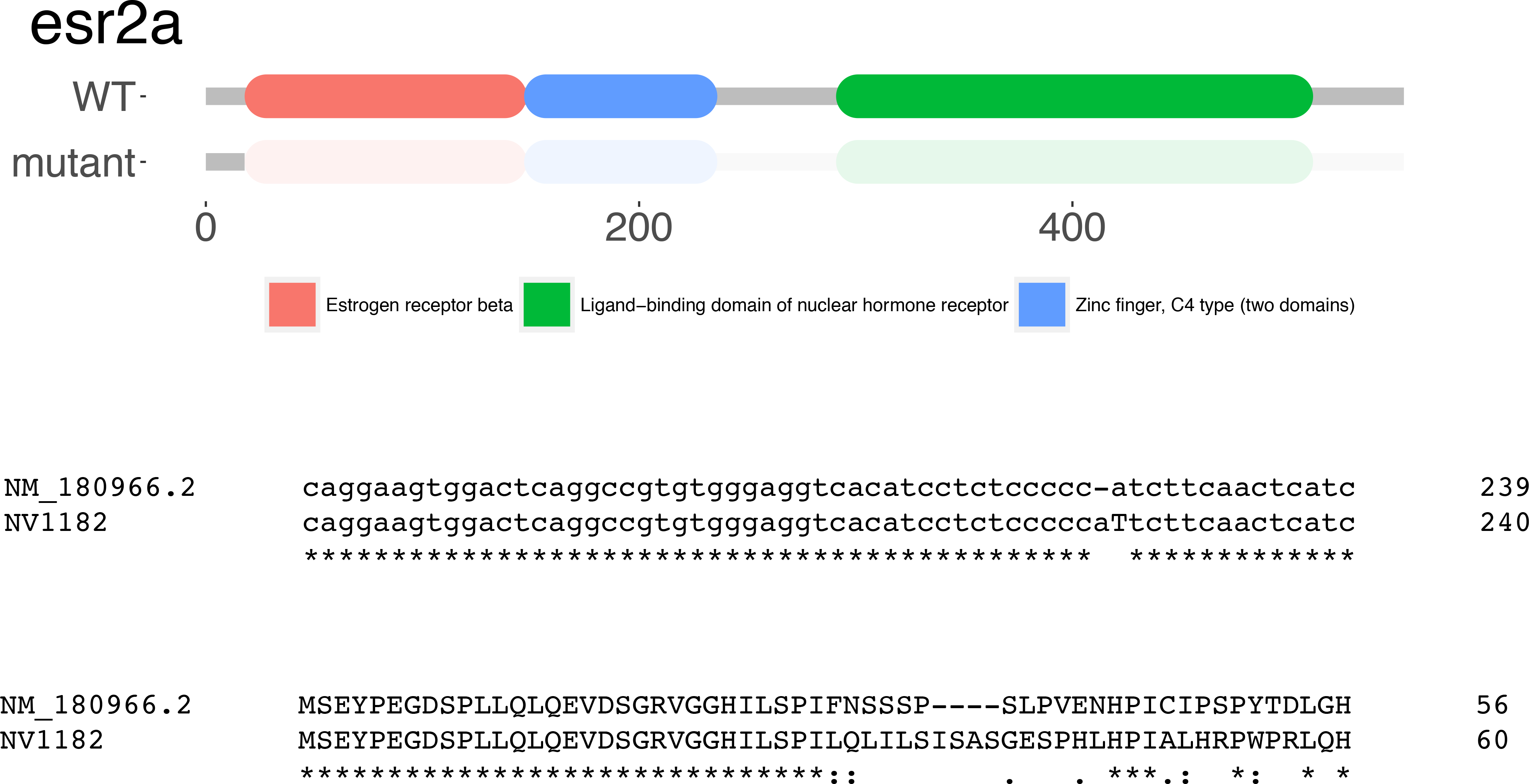

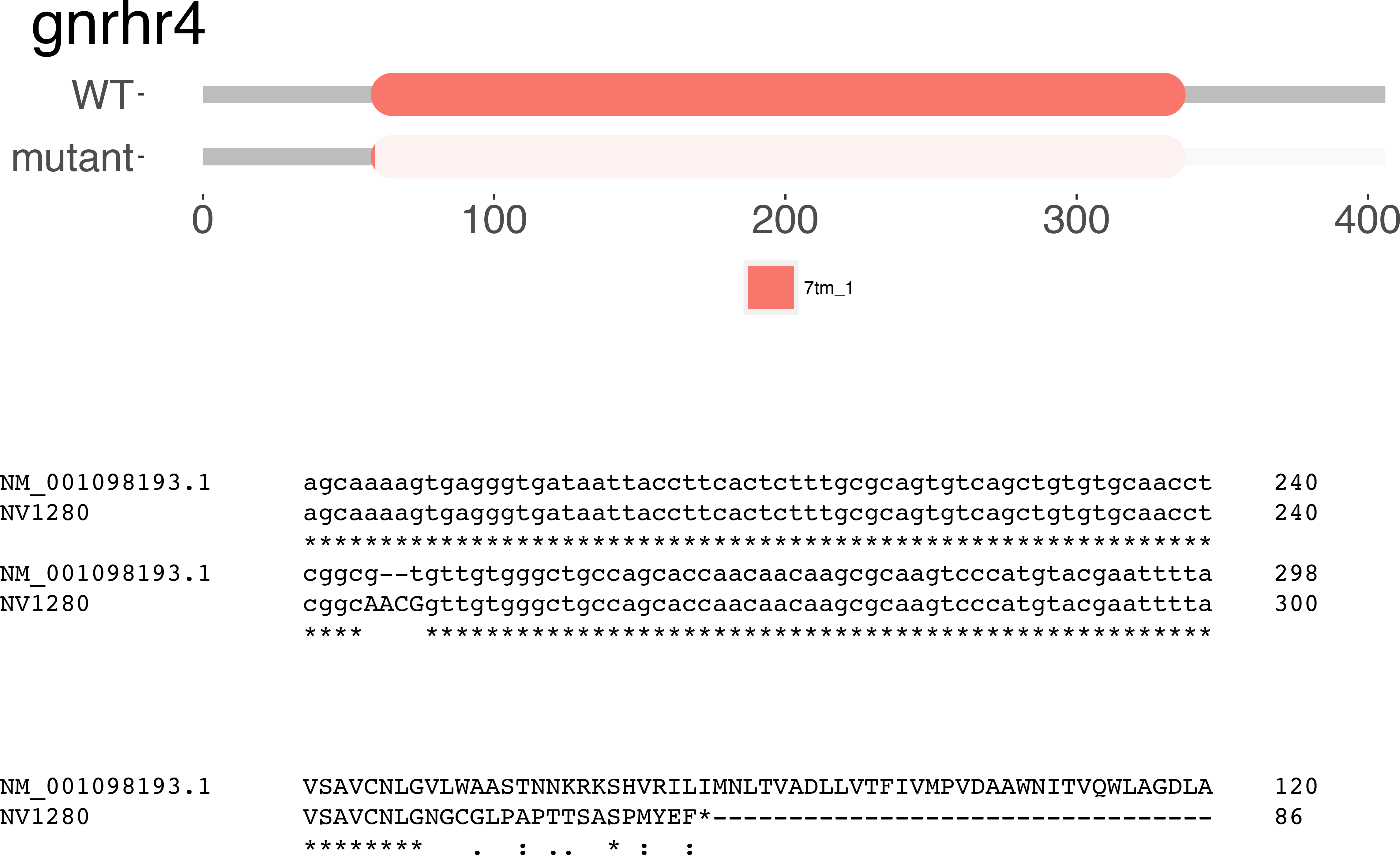

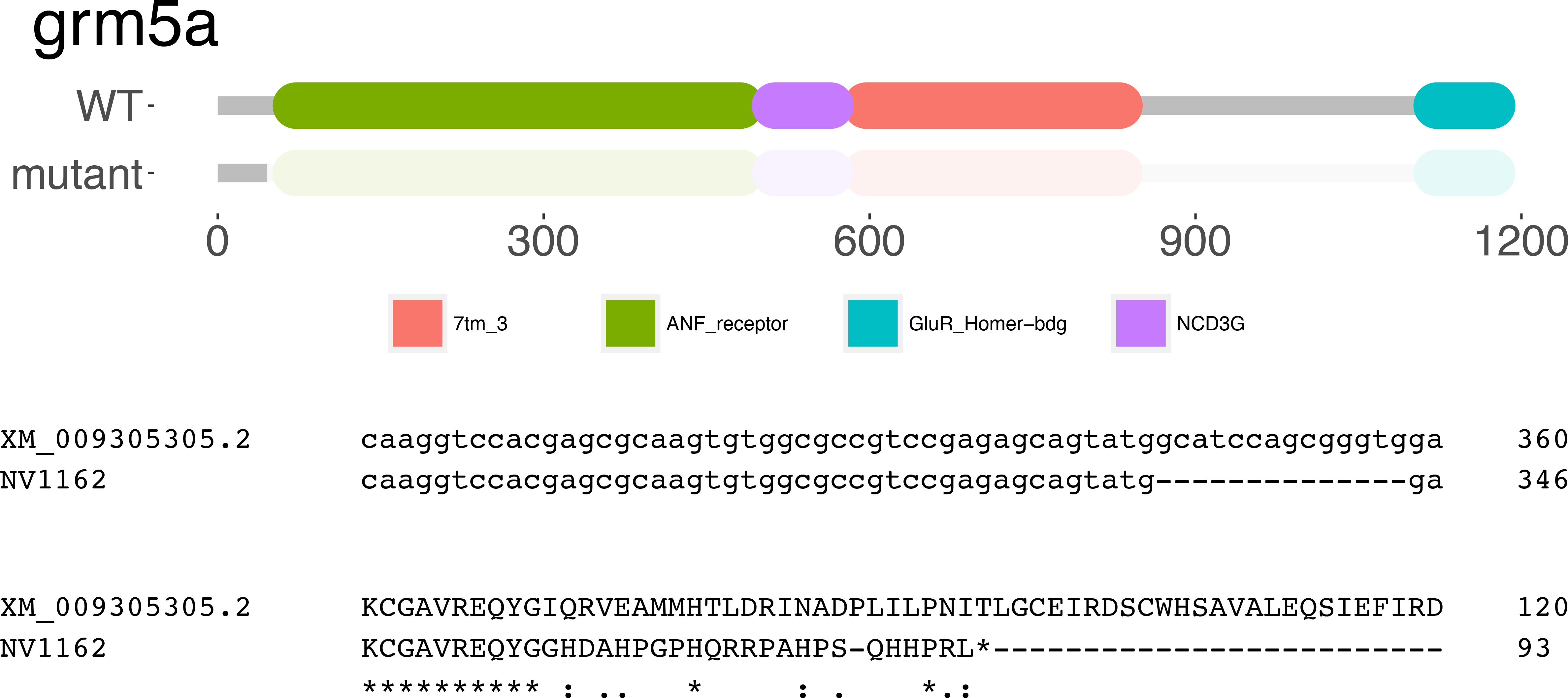

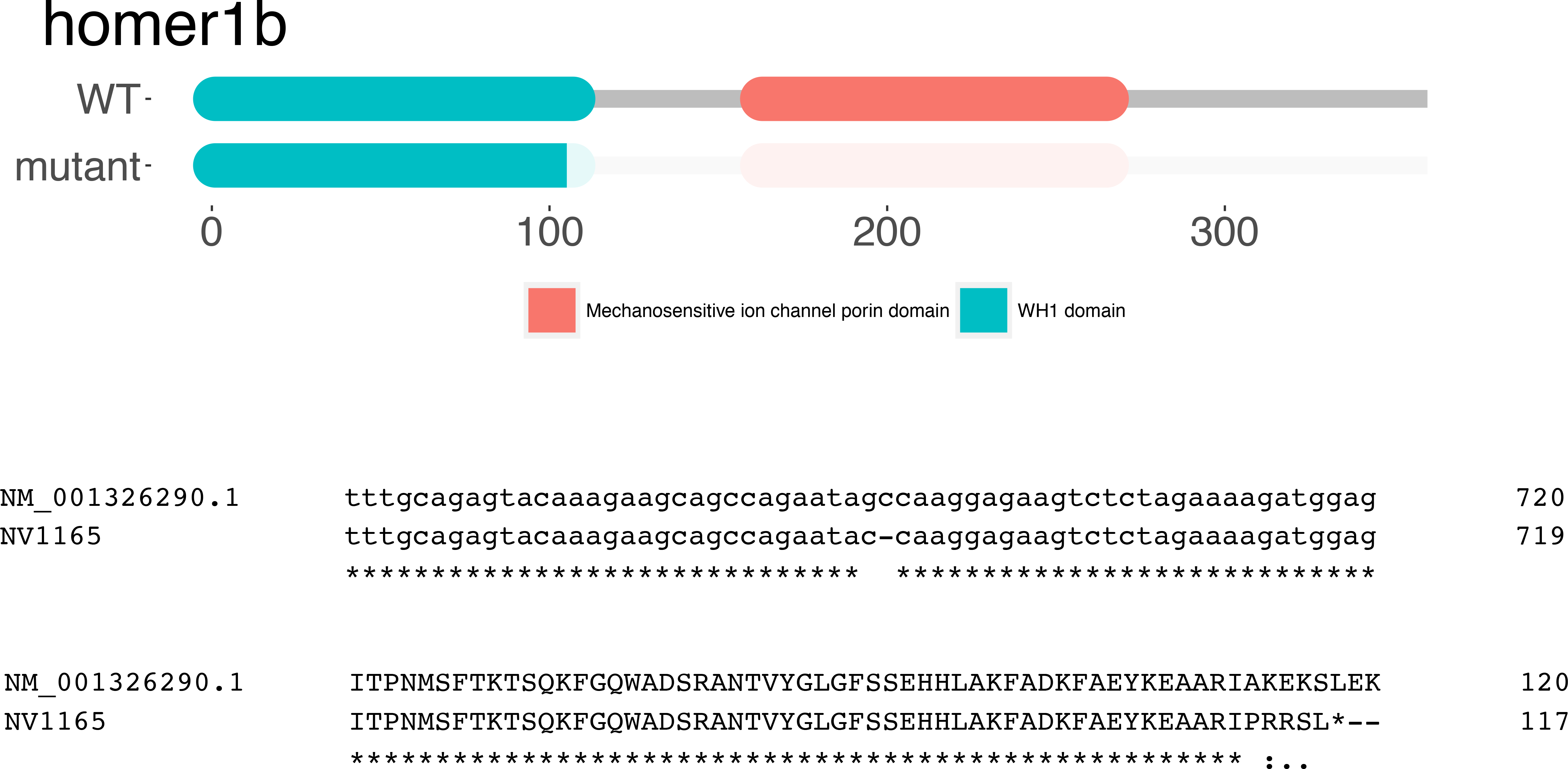

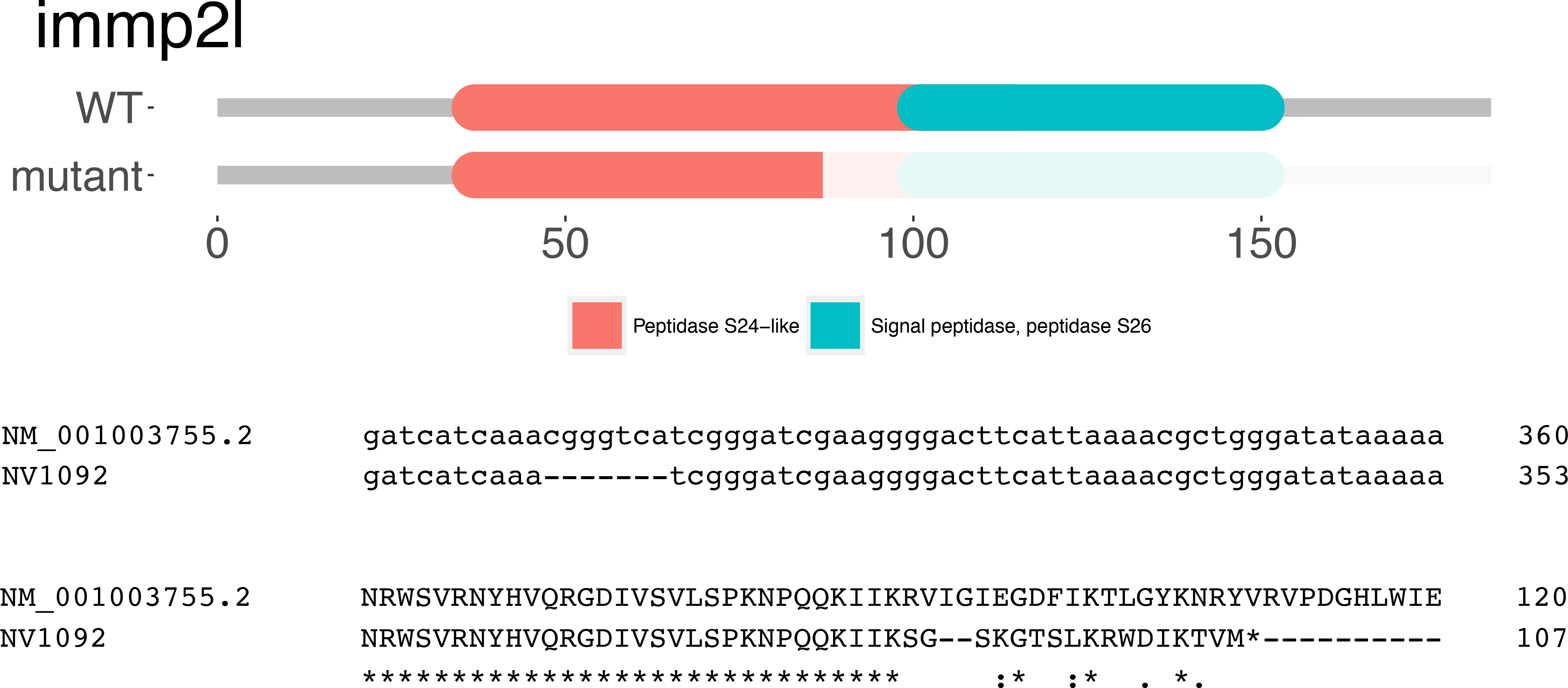

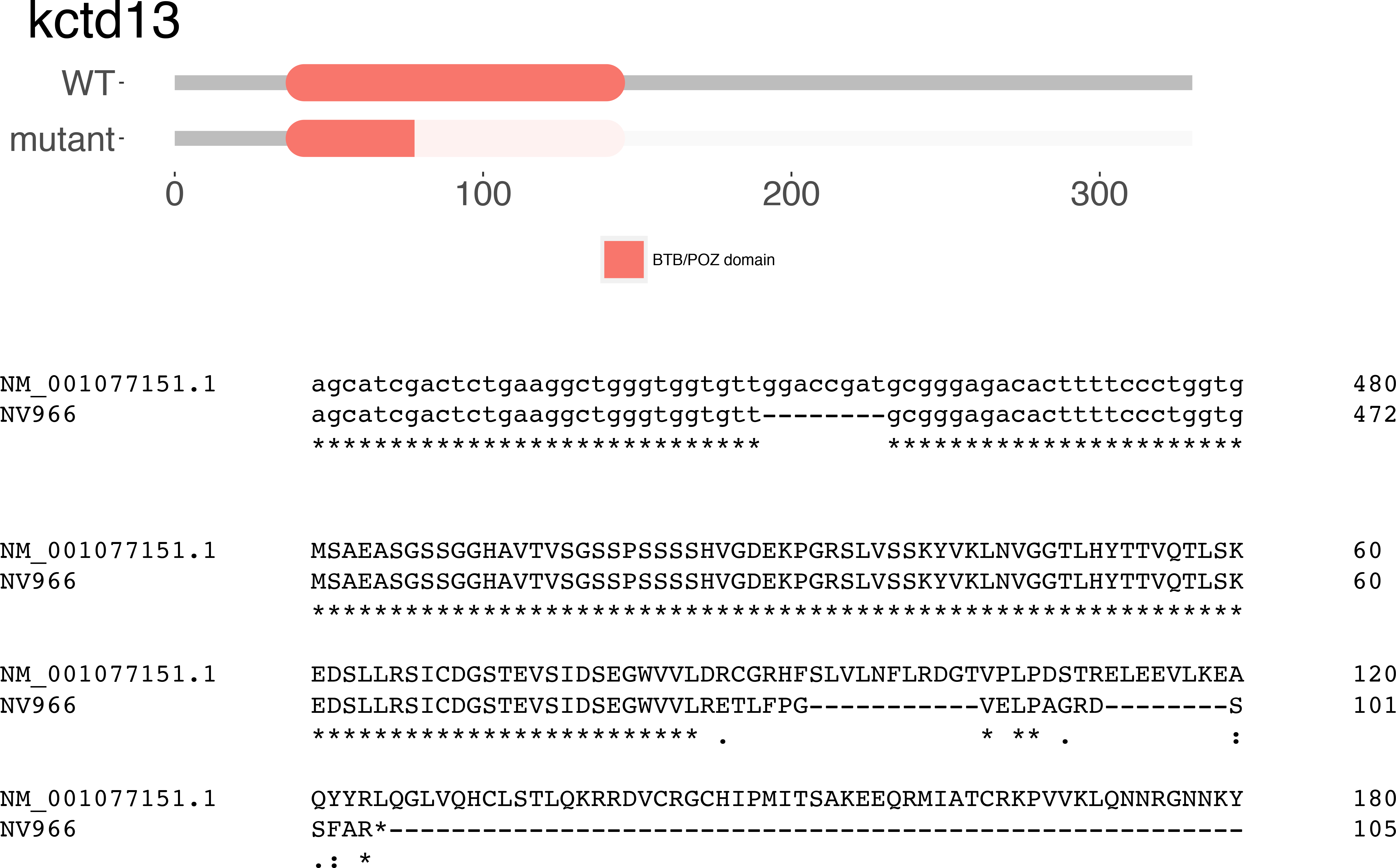

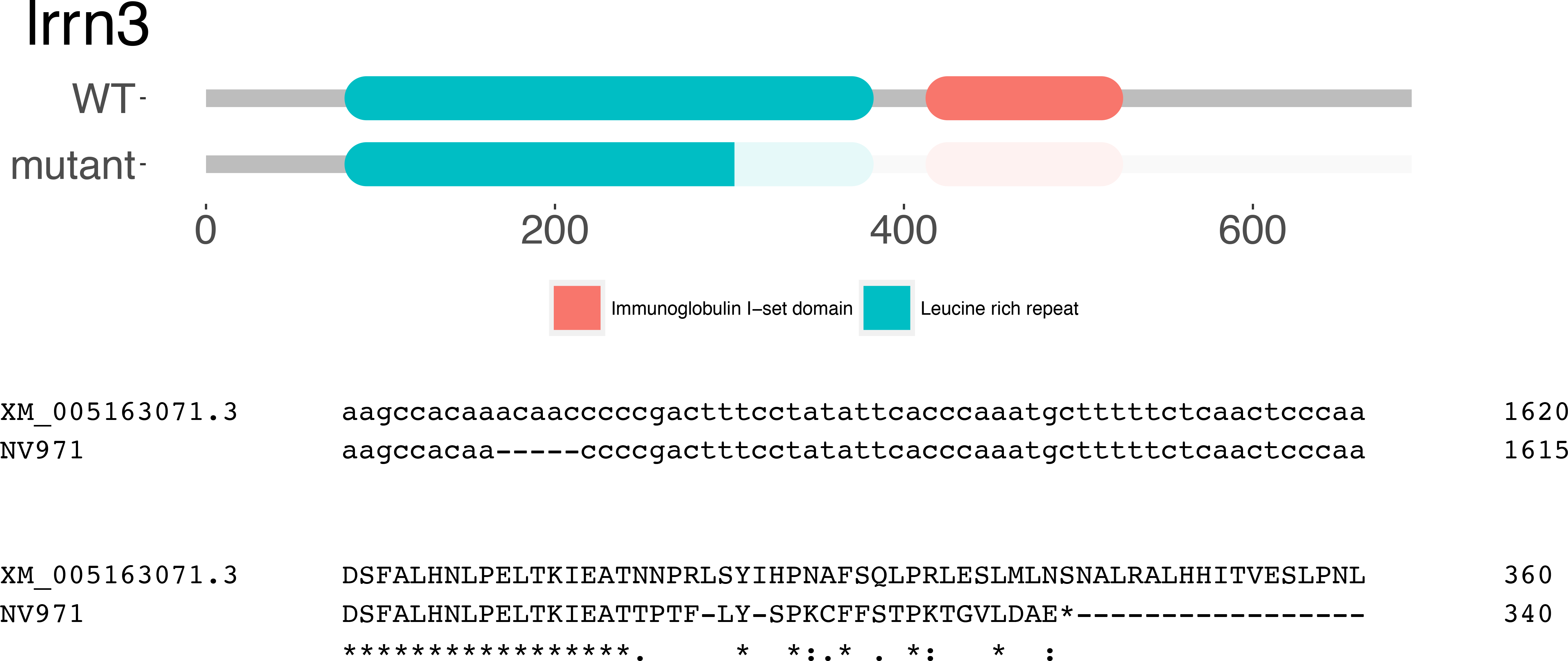

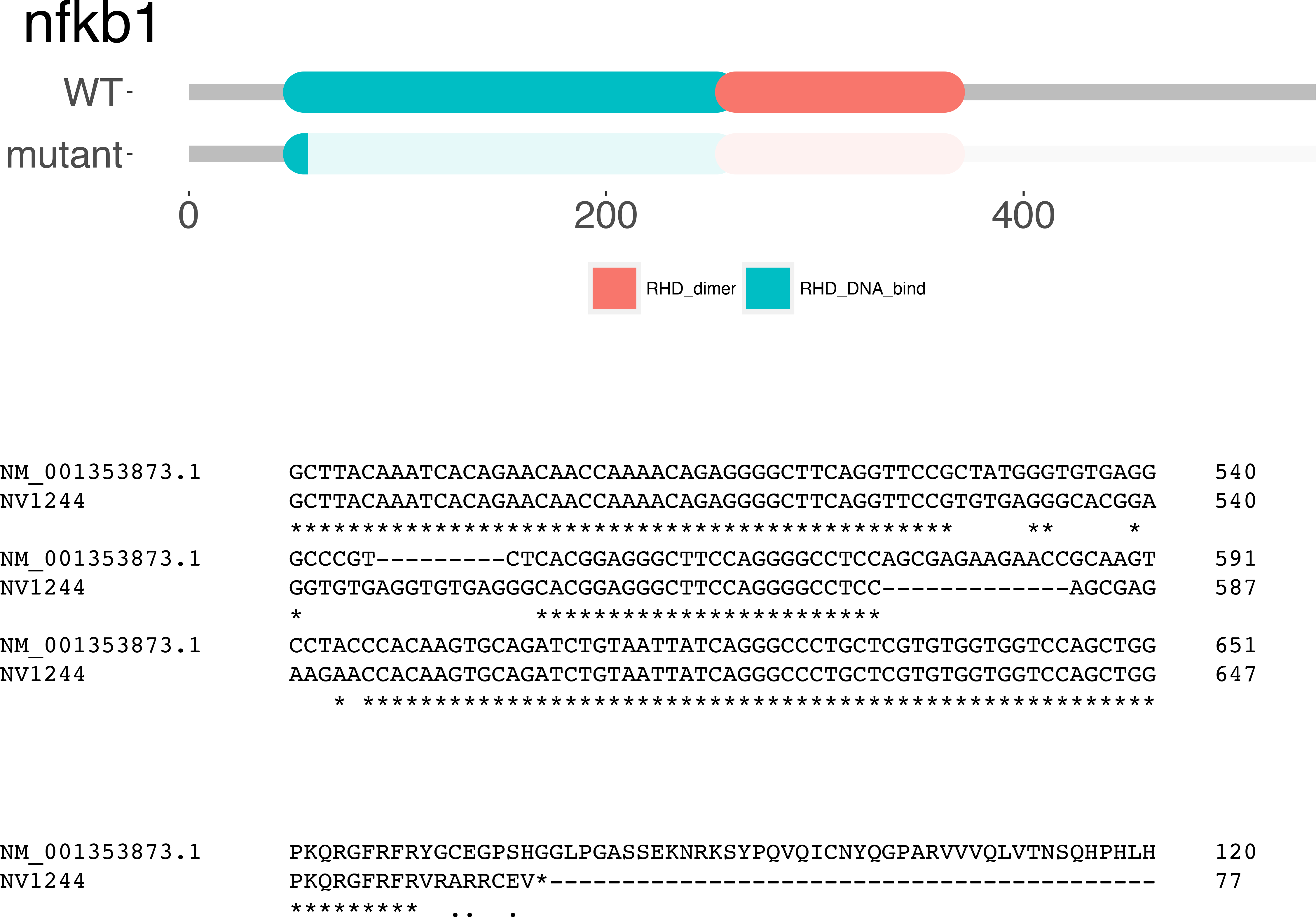

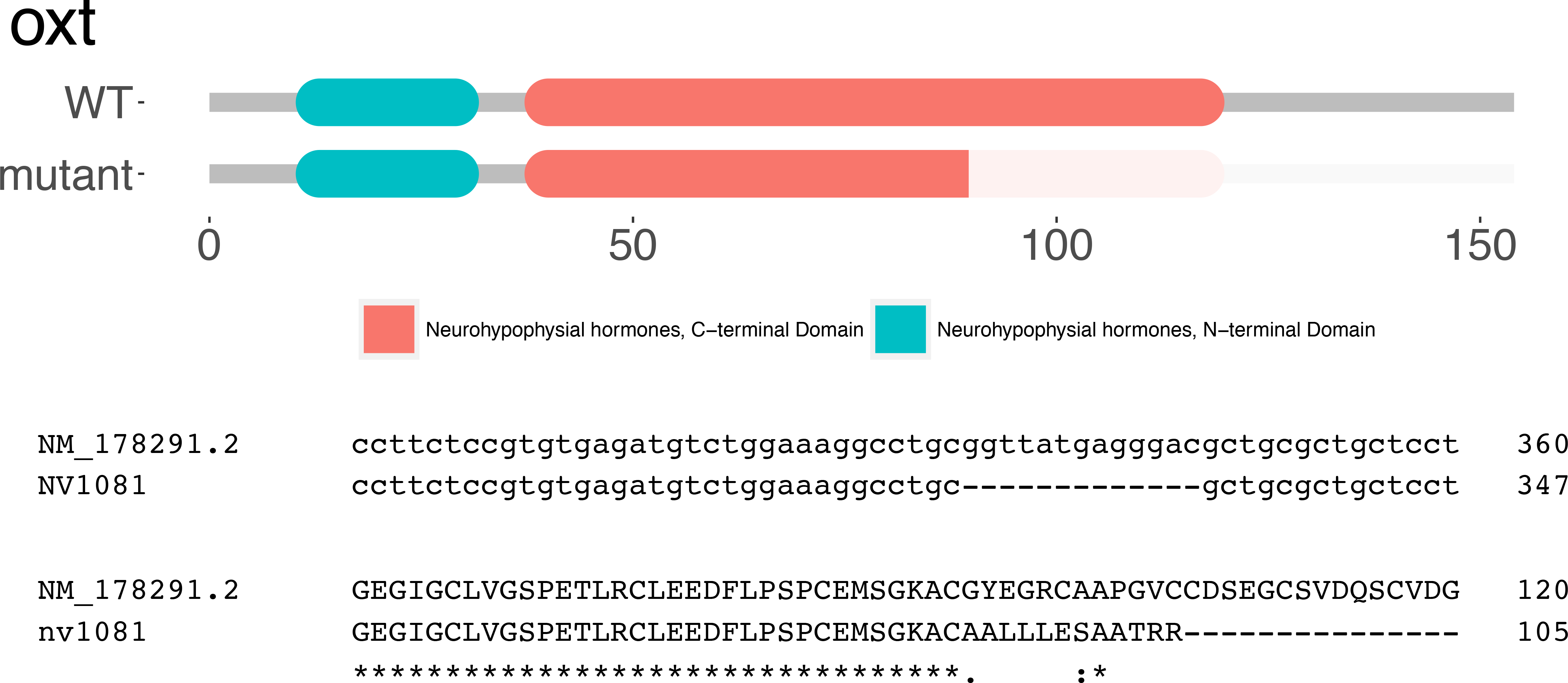

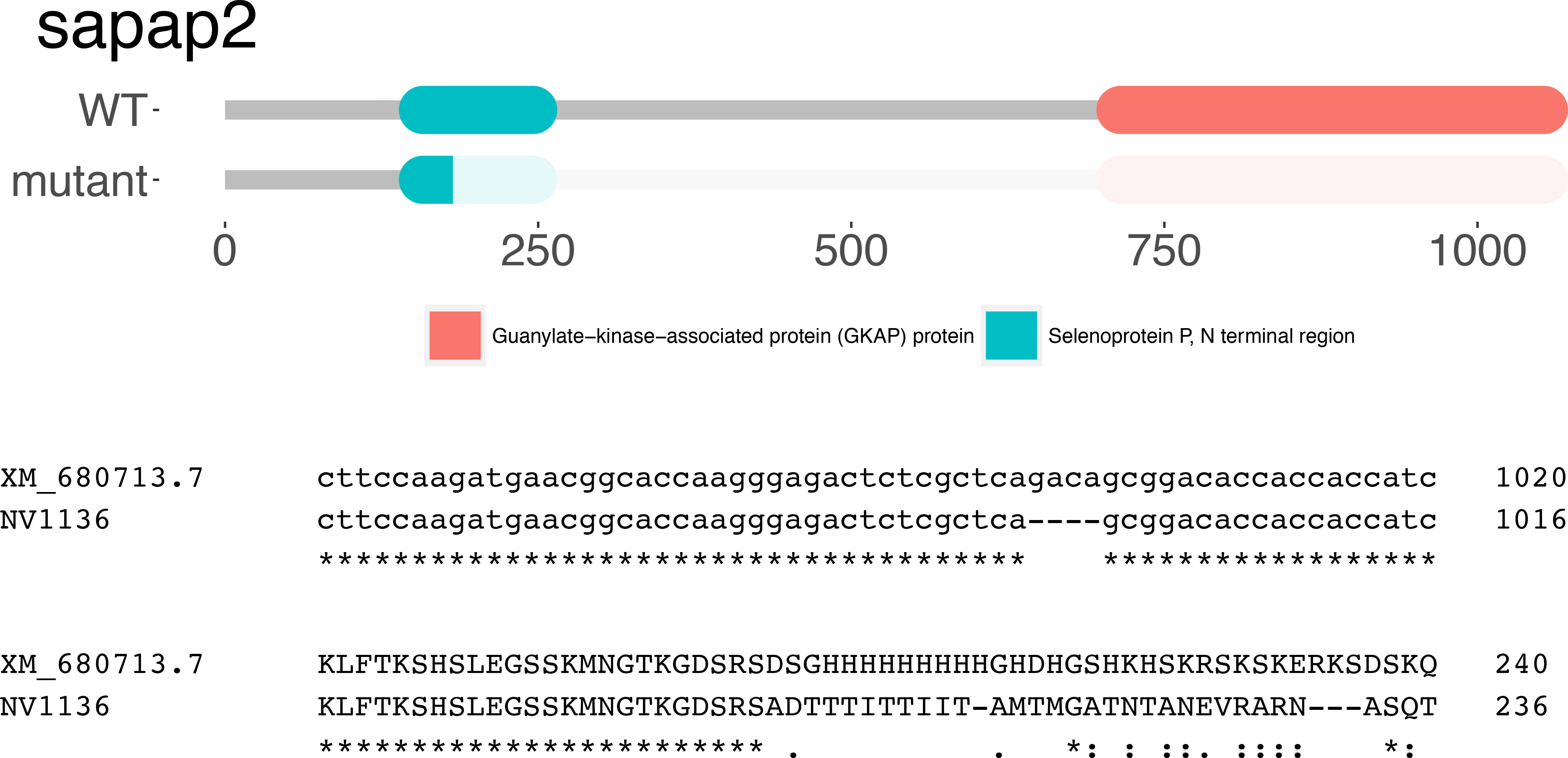

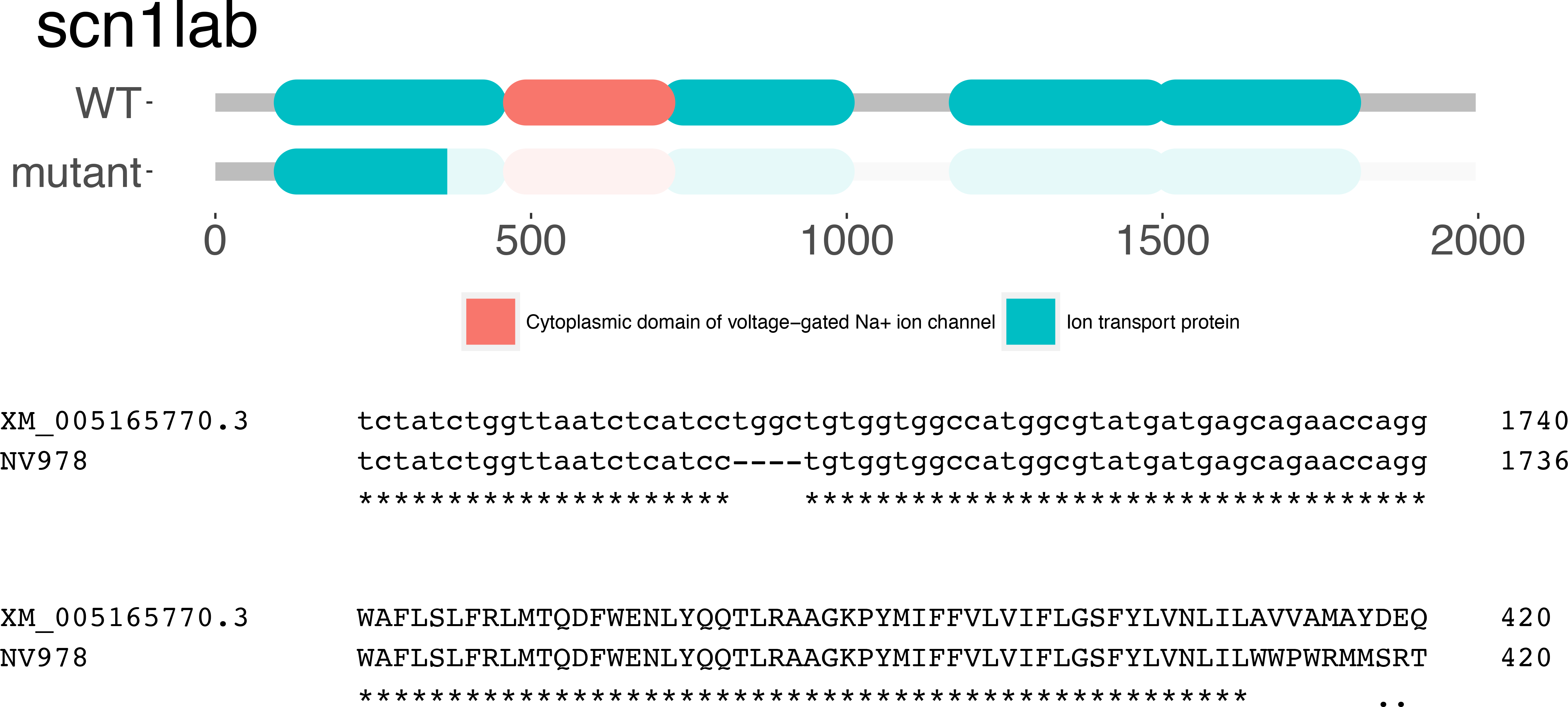

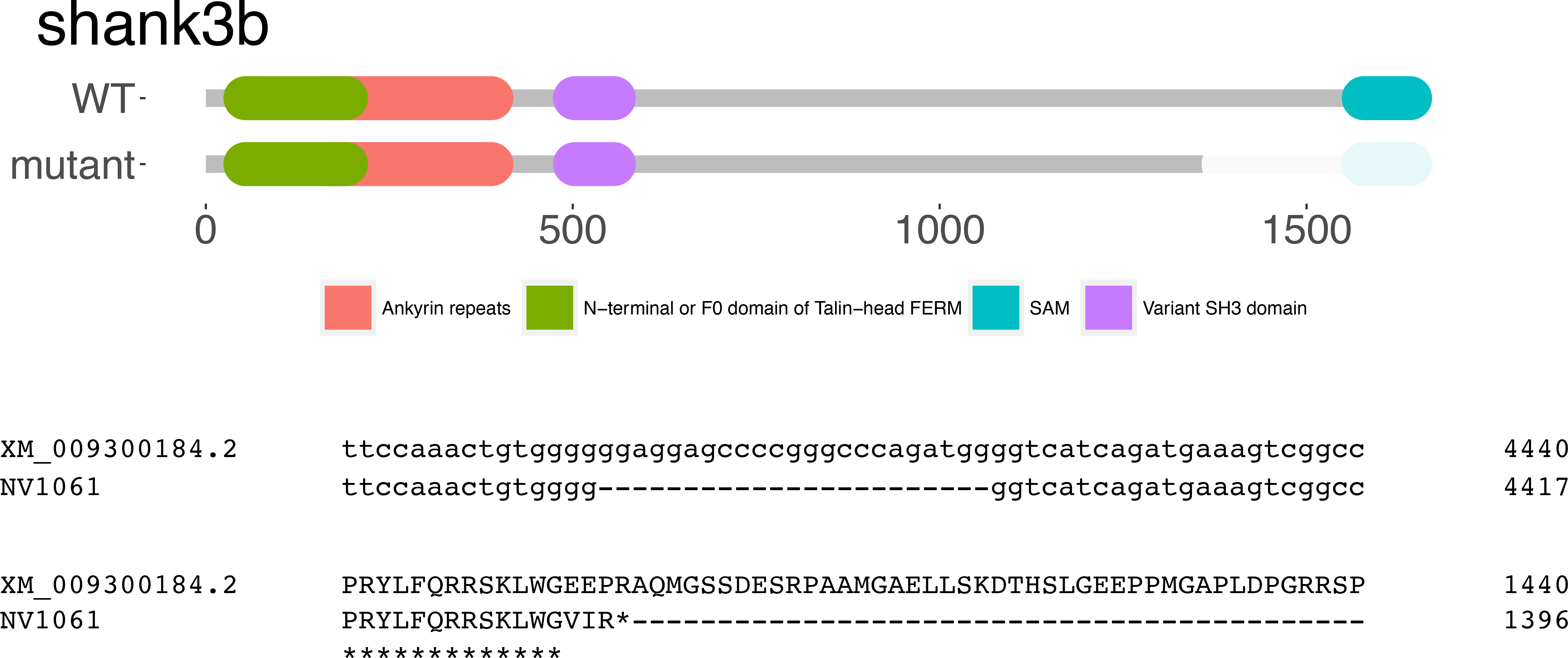

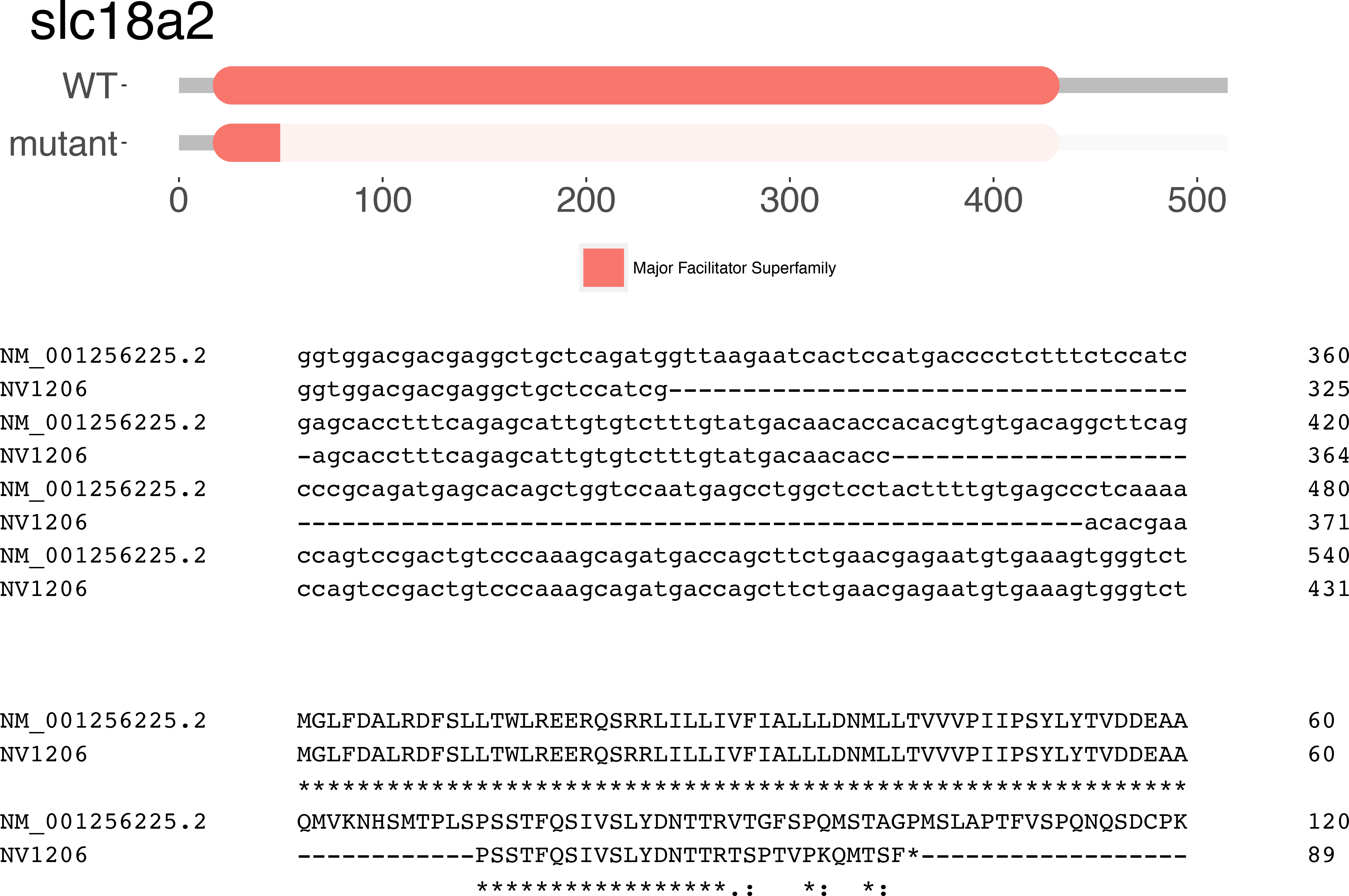

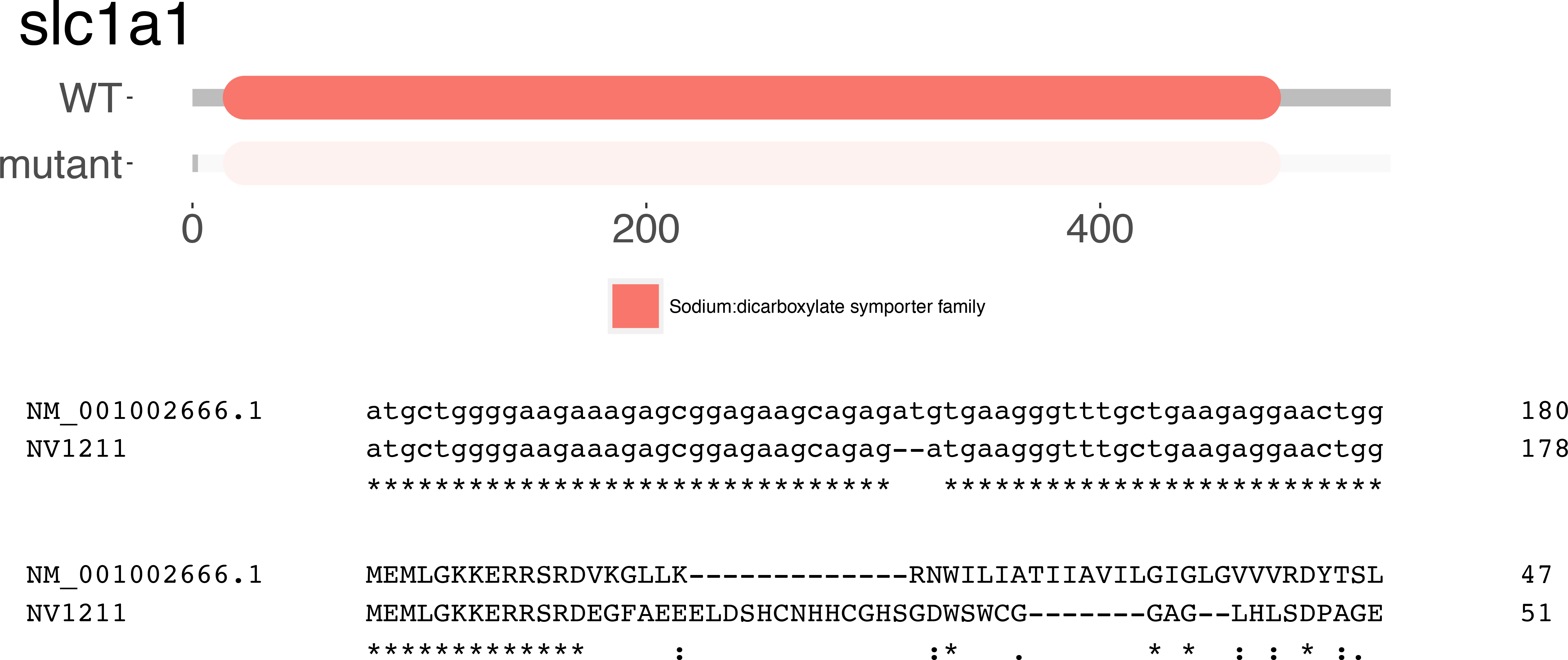

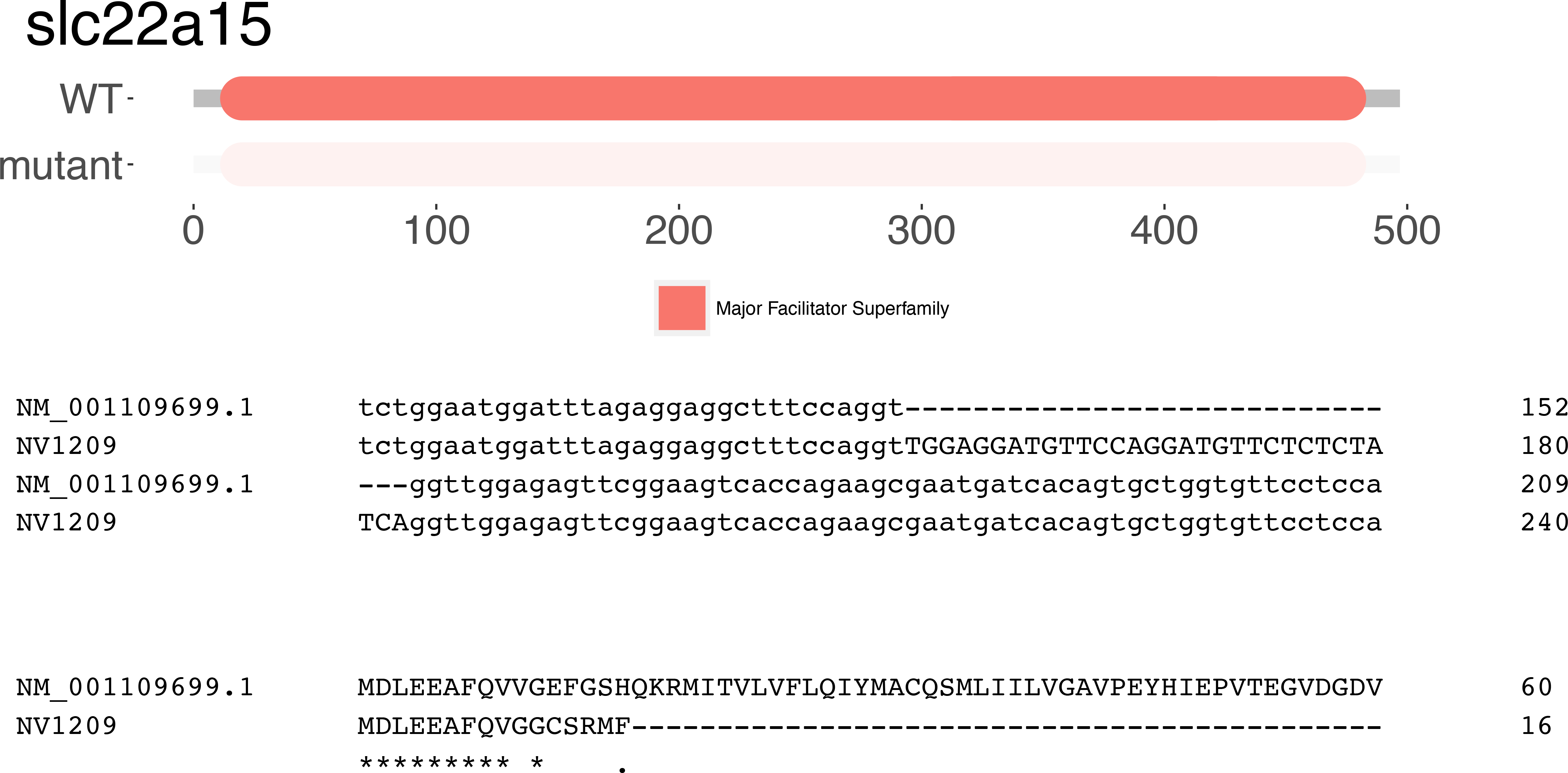

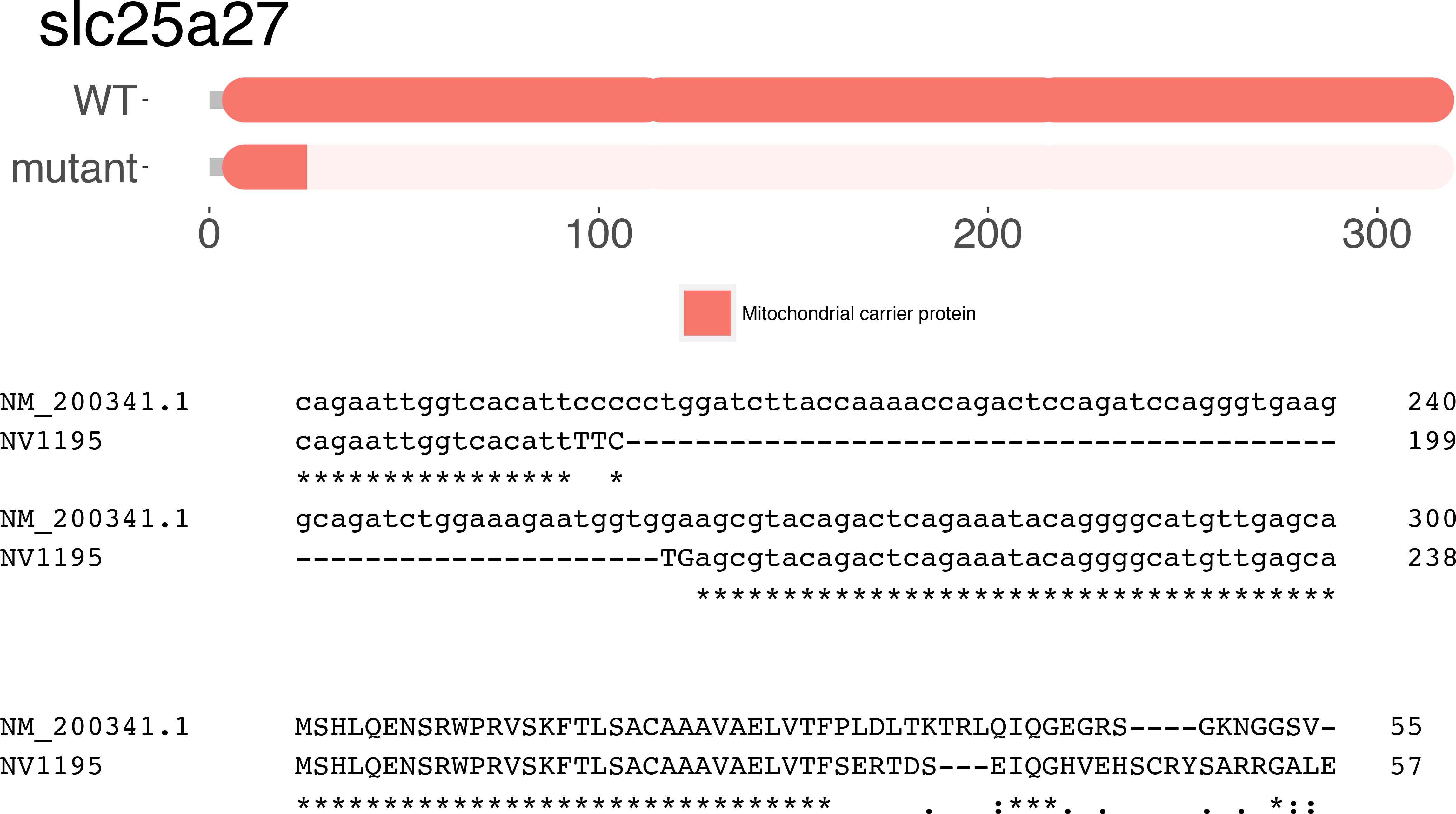

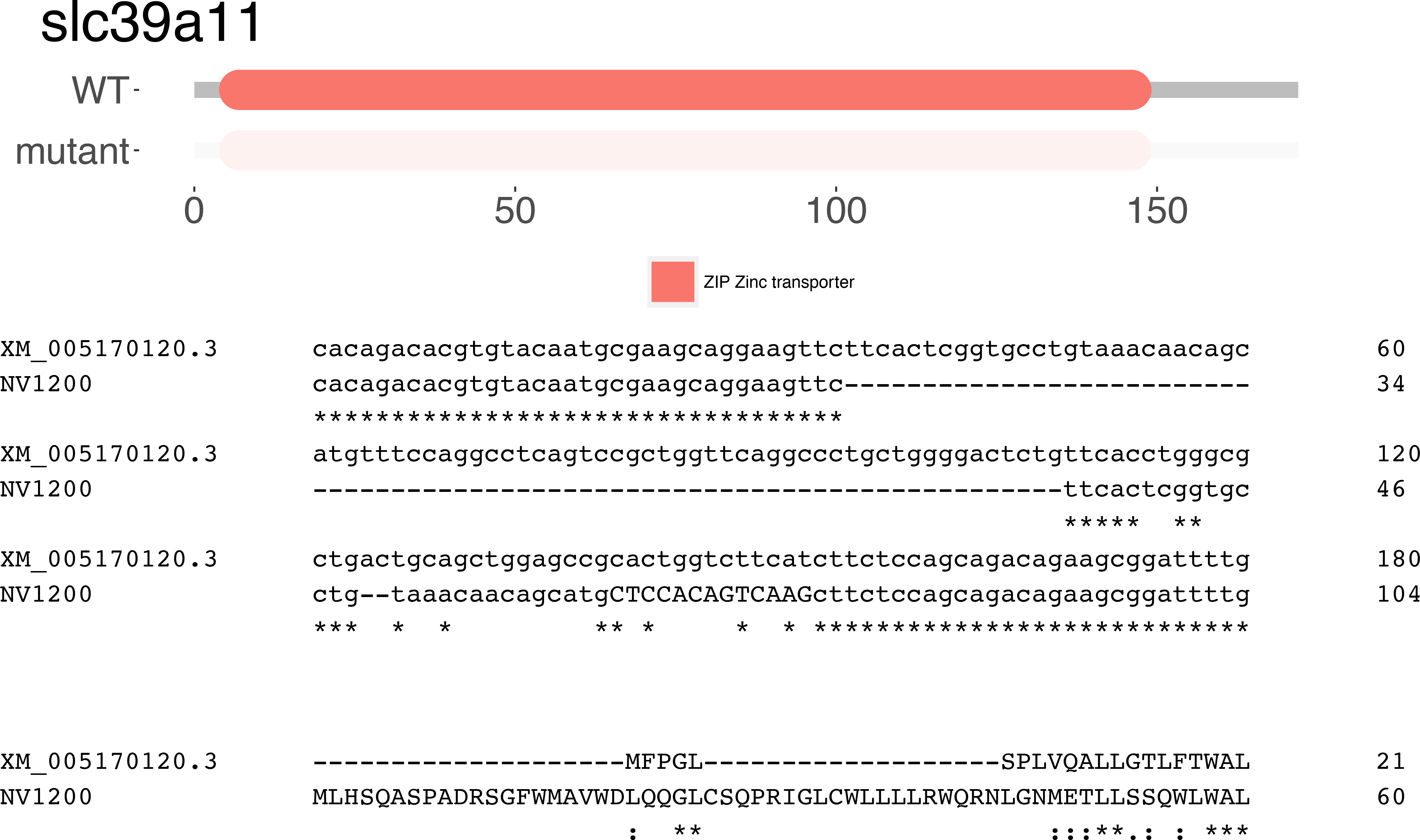

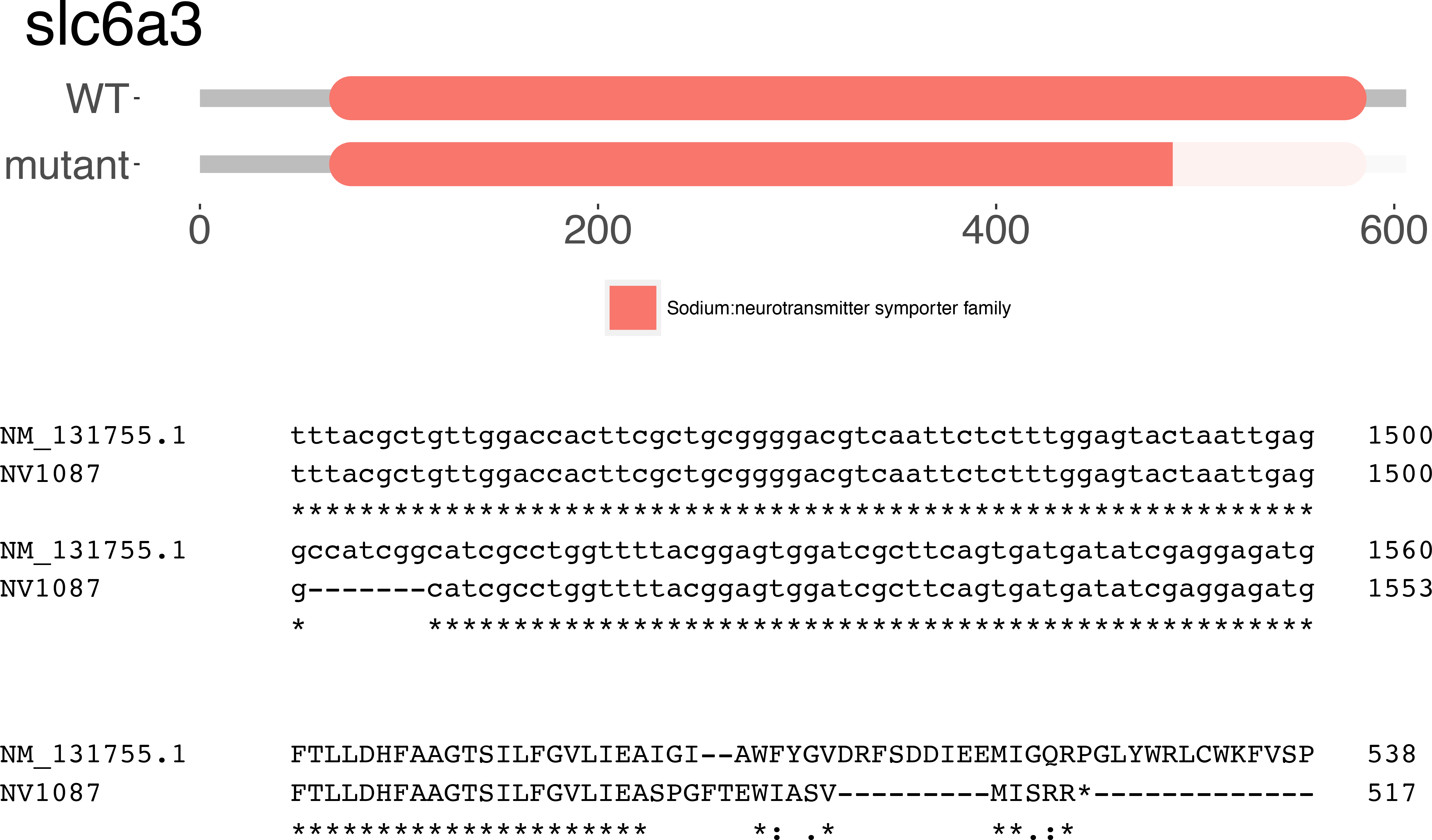

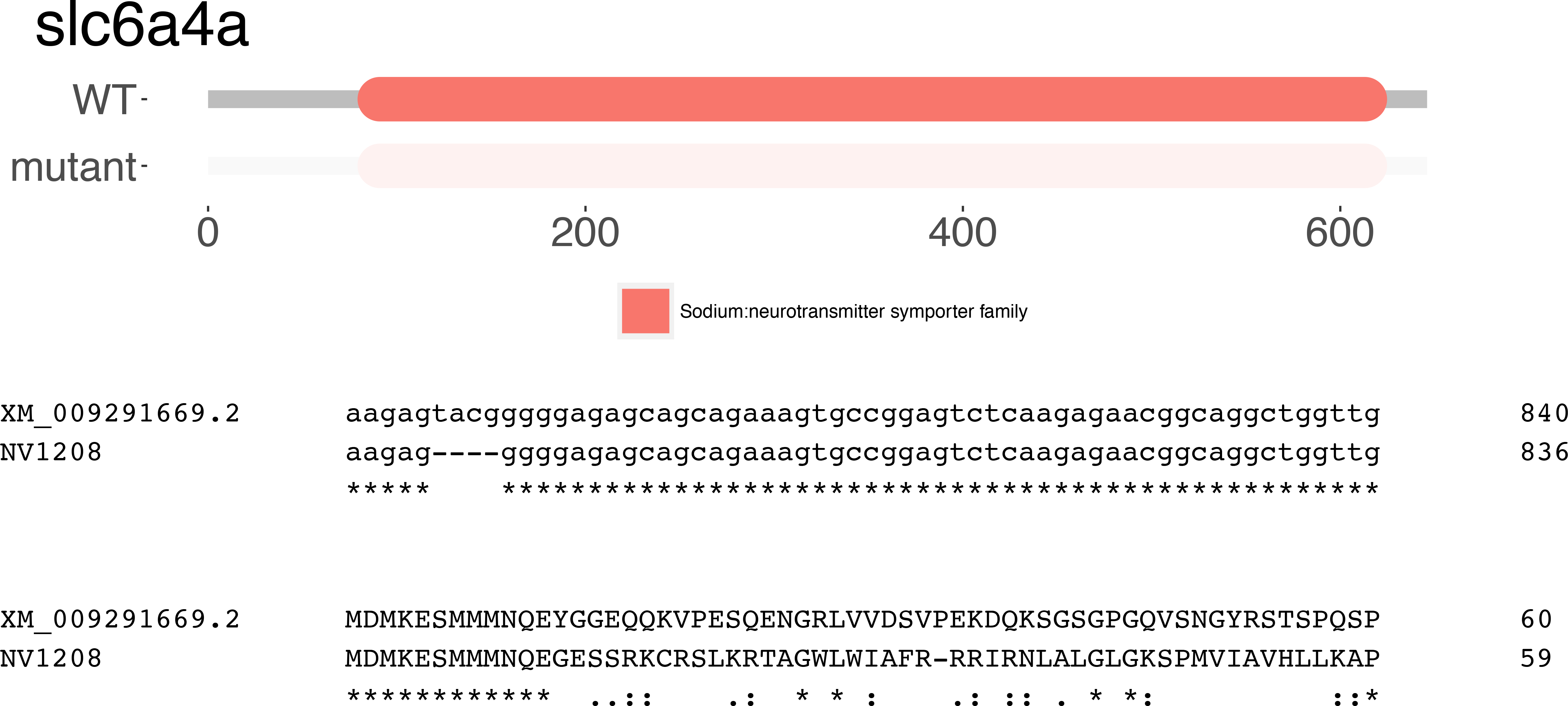

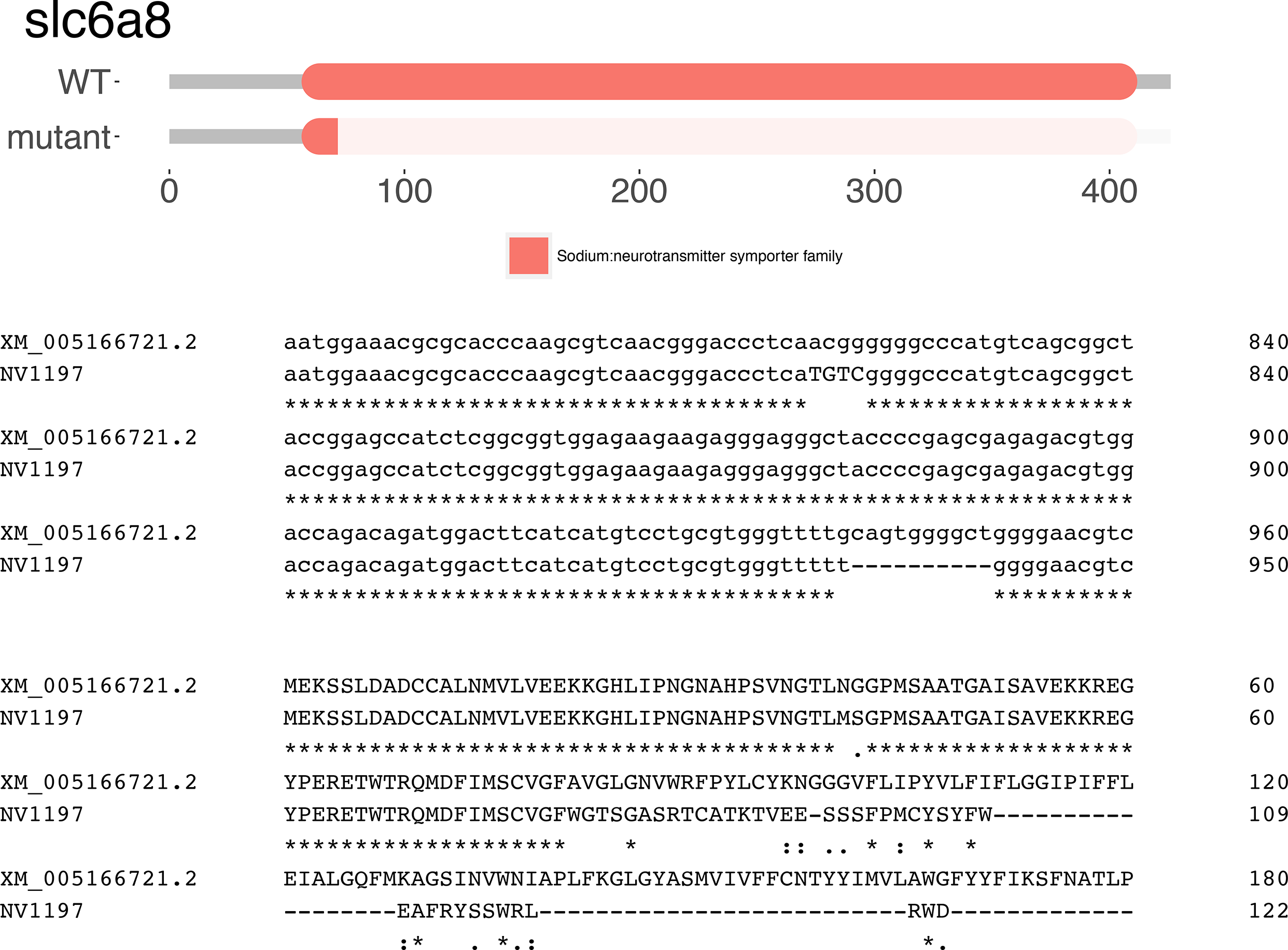

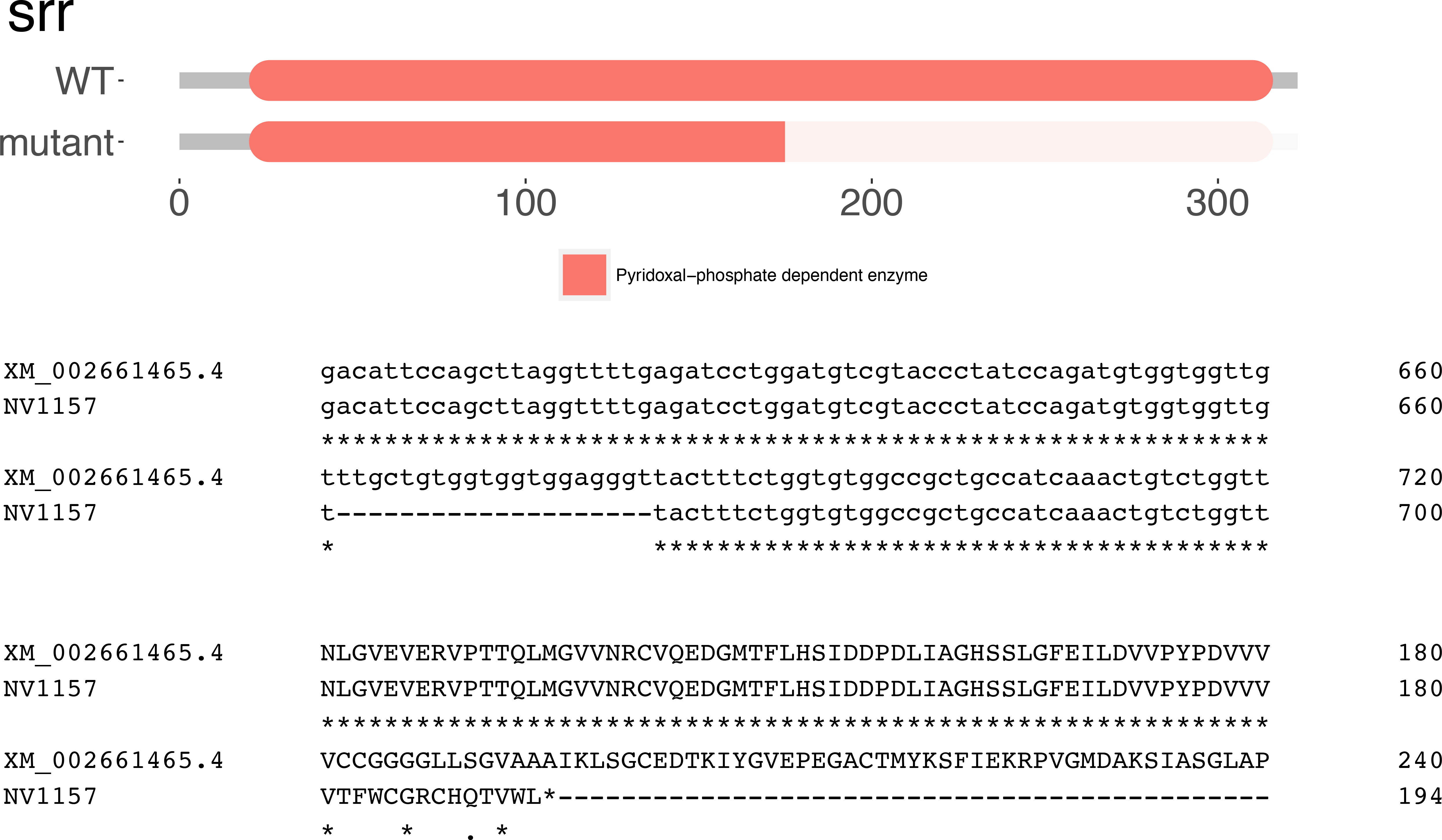

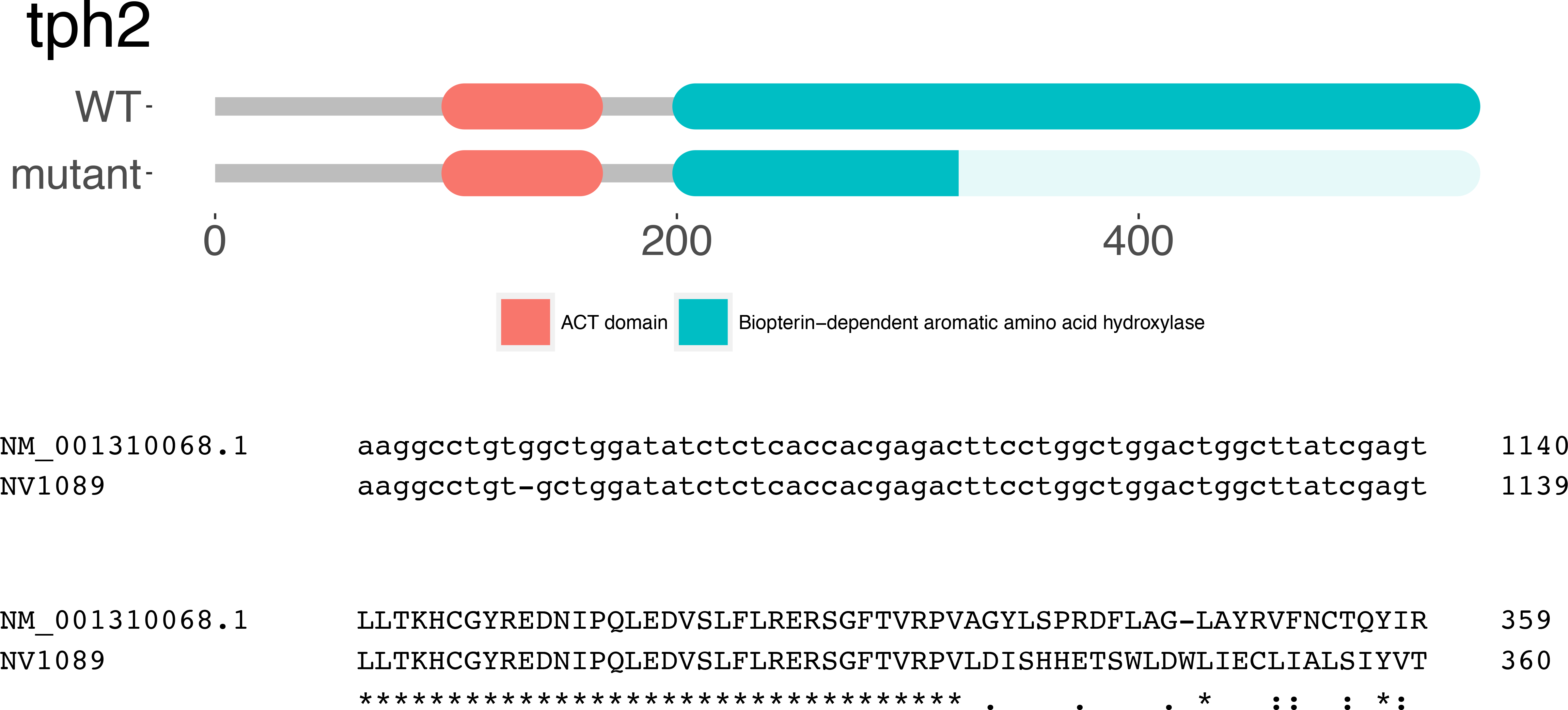

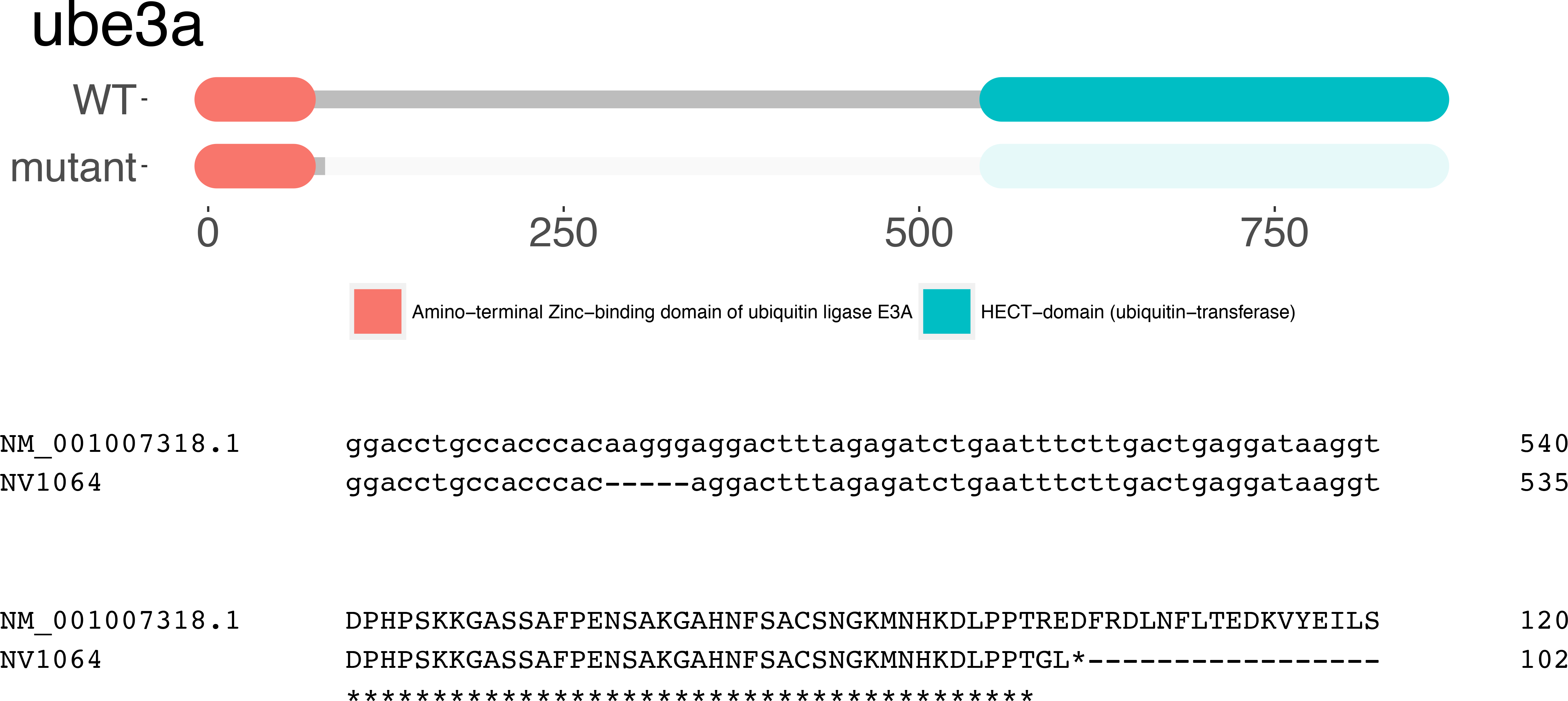

**Supplemental File 2.**
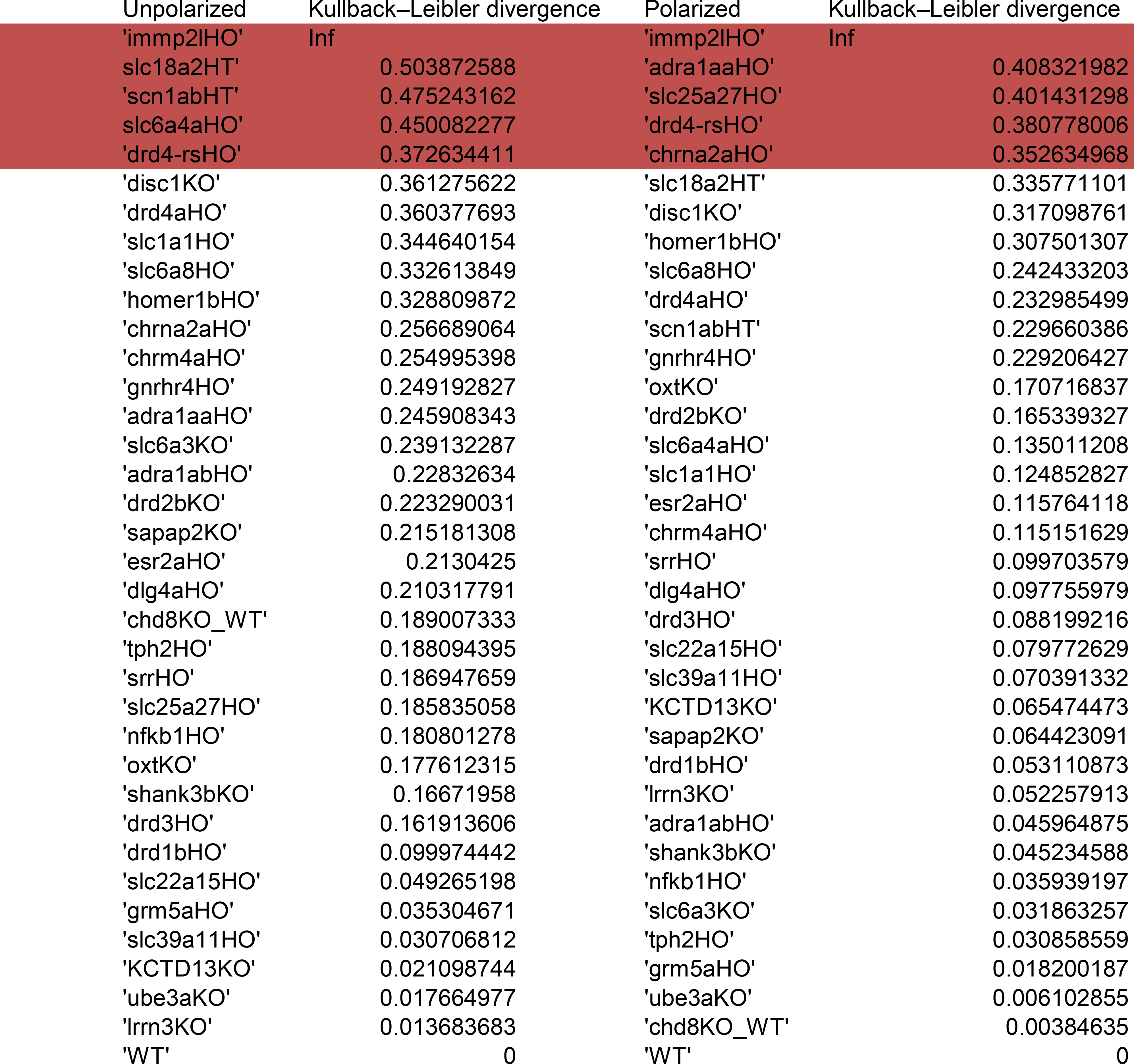

